# SPARC Reveals Distinct Osteoclastogenesis-Associated Malignant States in Metastatic Osteosarcoma

**DOI:** 10.64898/2026.07.07.736532

**Authors:** Shiyi Wu, William Cameron Walker, Bikesh Nirala, Carolyn A. Martin, Jason T. Yustein, Md. Abul Hassan Samee

**Affiliations:** Department of Integrative Physiology, Baylor College of Medicine, Houston, TX, USA; Department of Pediatrics, Emory University School of Medicine, Atlanta, GA, USA

**Author notes:** Correspondence: Jason T. Yustein, MD, PhD, Md. Abul Hassan Samee, PhD. These authors contributed equally to this work (Shiyi Wu, William Cameron Walker).

## Abstract

Single-cell studies of metastatic cancer have revealed extensive transcriptional heterogeneity and cellular plasticity as malignant populations establish themselves in distant organs. The growing availability of paired single-cell profiles from primary tumors and matched metastases offers new opportunities to investigate how malignant-cell states evolve during metastasis in relation to progression within the primary tumor. Static comparisons between these sites have identified differences in cellular composition and molecular programs, but can reduce heterogeneous populations to site-level summaries and overlook changes in gene-expression programs along possible routes of metastatic progression. Investigating these dynamics calls for trajectory inference that reconstructs state progression within each condition and transitions between conditions. However, existing trajectory inference tools have limitations in jointly modeling these relationships under biological ordering constraints. Traditional and widely used trajectory inference tools, such as Monocle 3 and Slingshot, do not impose direction constraints based on biological knowledge or the sampling design during graph construction. Optimal transport-based tools, such as moscot, incorporate temporal order to infer cross-condition correspondences, but these mappings alone do not reconstruct progression within each condition. More recent methods, including CStreet and CellRank 2, incorporate both transition types, but construct the two components separately through distinct modeling or graph-construction strategies. Here we introduce SPARC (Shortest Path Algorithmic Reconstruction of Cross-state Evolution), a unified graph-based optimization framework that evaluates both types of relationships under a common connectivity formulation to reconstruct complete, directionally constrained candidate trajectories. Across benchmarks, SPARC improved recovery of candidate metastatic precursors and complete progression routes in simulations, showed strong directed-trajectory reconstruction in conventional time-course settings, and performed competitively in real tumor lineage-tracing datasets. To examine metastatic progression in a model where lung lesions arise through spontaneous dissemination, we generated, to our knowledge, the first paired scRNA-seq dataset of primary osteosarcoma tumors and spontaneous lung metastases from the same mice, together with the first patient-derived osteosarcoma Visium HD dataset. The biological analyses revealed differences among metastatic tumor states in their association with osteoclast-like myeloid populations and candidate supportive signals, suggesting that certain malignant states may preferentially promote osteoclastogenesis. Together, SPARC provides a framework for investigating biologically ordered cell-state progression and identifying state-specific programs associated with metastatic disease.

## Introduction

Single-cell profiling and lineage-tracing studies have revealed that malignant populations differ in their metastatic potential and can undergo substantial transcriptional plasticity during tumor progression^1–6^. These findings raise a central question in metastasis research: why can some primary-tumor populations give rise to metastatic disease while others cannot, and how do molecular programs emerge and change as malignant cells progress from the primary tumor to distant organs?

An increasing number of studies now generate paired scRNA-seq datasets from primary tumors and metastatic lesions collected from the same patient or animal^5,7^. These datasets provide new opportunities to investigate metastatic progression against a shared biological background. Static comparisons of cellular composition and gene expression have identified important metastasis-associated programs, but site-level summaries can obscure the heterogeneous transcriptional changes occurring along different progression routes. For example, a molecular program may increase along one metastatic route but decrease along another, causing these distinct patterns to be obscured when populations are analyzed collectively. Understanding how molecular programs evolve during metastatic progression therefore requires trajectory inference that jointly reconstructs cell-state progression within each anatomical site and transitions between sites.

Traditional and widely used trajectory inference tools, such as Monocle 3, Slingshot and TinGa, have demonstrated strong performance across a range of trajectory structures in published benchmark evaluations^8,9^. They were developed to infer trajectories from single-cell snapshots and have been extensively applied to canonical developmental and differentiation processes^10,11^. In these settings, progression can often be interpreted along an immature-to-mature differentiation axis. However, their standard graph-construction procedures do not enforce direction constraints based on the biological ordering of sampled conditions. This limitation is particularly relevant to metastatic disease, where tumor-cell plasticity can allow some intermediate primary-tumor populations and metastatic populations to occupy similar transcriptional states. Although specifying a primary-tumor root determines where a trajectory begins, it does not constrain subsequent transitions to follow the prescribed primary-to-metastasis direction. Consequently, inferred paths may pass from primary to metastatic states and subsequently return to primary populations, conflicting with the biological ordering assumed for the analysis and complicating the identification of candidate metastatic precursors.

Methods developed for time-course data offer a way to represent transitions between ordered conditions. Optimal transport (OT) approaches, including Waddington-OT and moscot, estimate couplings between cell populations designated as an earlier source and a later target^12,13^. Applied to primary and metastatic samples, these couplings describe cross-site correspondences, but do not directly reconstruct progression within either site. Progression toward a candidate primary precursor and further state changes within a metastatic lesion therefore require an additional representation of within-condition relationships.

More recently developed methods, including CellRank 2 and CStreet, incorporate both within-condition and cross-condition transitions^14,15^. However, the two components are constructed separately using distinct modeling or graph-construction procedures. CellRank 2 combines neighborhood-based transition weights within time points with optimal transport-derived transition weights between time points. Their relative contributions are controlled by a user-specified mixing hyperparameter. In heterogeneous tumors, the relative importance of local progression and dissemination may vary among cellular populations and datasets, making a single global mixing parameter difficult to select and potentially insufficient to represent state-specific differences. CStreet applies one connection-frequency definition but uses different neighbor-search pools and separately determined edge-retention thresholds. These modeling asymmetries can change the relative support for local progression and cross-condition transitions, potentially affecting the inferred topology of primary-to-metastatic progression and the placement of its precursor and intermediate states. To address this challenge in the clinically relevant context of metastatic progression, we developed SPARC (Shortest Path Algorithmic Reconstruction of Cross-state Evolution).

SPARC is a graph-based optimization framework for trajectory inference within and across biologically ordered conditions. Building on the graph-abstraction rationale of PAGA^16^, it uses one global neighborhood reference in which each cell selects neighbors outside its own cluster across all included conditions. Both transition types are quantified using the same observed-to-expected connectivity measure and contribute to one trajectory optimization objective. The specified biological ordering constrains the permitted directions, and Dijkstra’s algorithm selects candidate trajectories with the lowest accumulated connection cost^17^. This common construction provides a consistent basis for positioning candidate precursors and intermediate states, without a separate coefficient for balancing within-condition and cross-condition connections.

Simulation benchmarks showed that SPARC reconstructs progression within and across conditions, with improved topology and precursor-to-metastatic trajectory recovery in the tested heterogeneous primary–metastatic setting. LARRY^18^ and independent tumor lineage-tracing datasets^1–3^ provided complementary evidence for evaluating the inferred state relationships. Together, these evaluations demonstrate the methodological value of placing local progression and cross-condition transitions under one reconstruction objective.

To examine how SPARC resolves progression toward metastasis, we generated paired scRNA-seq profiles of primary osteosarcoma tumors and their spontaneous pulmonary metastases from the same immunocompetent C57BL/6 mice in a syngeneic orthotopic model^19^. To our knowledge, this represents the first paired primary–spontaneous metastatic mouse osteosarcoma scRNA-seq dataset. The inferred trajectories distinguished metastasis-associated programs from changes also observed along local primary-tumor progression. We further generated patient-derived Visium HD profiles of primary and metastatic osteosarcoma specimens, which, to our knowledge, constitute the first patient-derived osteosarcoma Visium HD dataset. Integrating trajectory inference with spatial analysis revealed heterogeneity among metastatic malignant states in their associations with osteoclast-like myeloid populations and candidate osteoclastogenic signals. These findings suggest that metastatic tumor states may differ in their potential to promote osteoclastogenesis, providing a state-specific hypothesis for functional testing. SPARC is also applicable to other ordered studies in which local progression is essential to interpreting transitions between conditions.

## Results

### SPARC reconstructs biologically ordered trajectories through unified graph-based optimization

Metastatic progression can involve diversification within the primary tumor before dissemination and further transcriptional changes after colonization. The cells sampled at the two sites capture states along these possible processes, but their sequence is not directly observed. To reconstruct candidate routes from primary precursors through intermediate states to metastatic populations, SPARC evaluates within-site and cross-site relationships using the same connectivity measure and one complete-route optimization objective (Fig. 1a–c).

**Figure 1.**
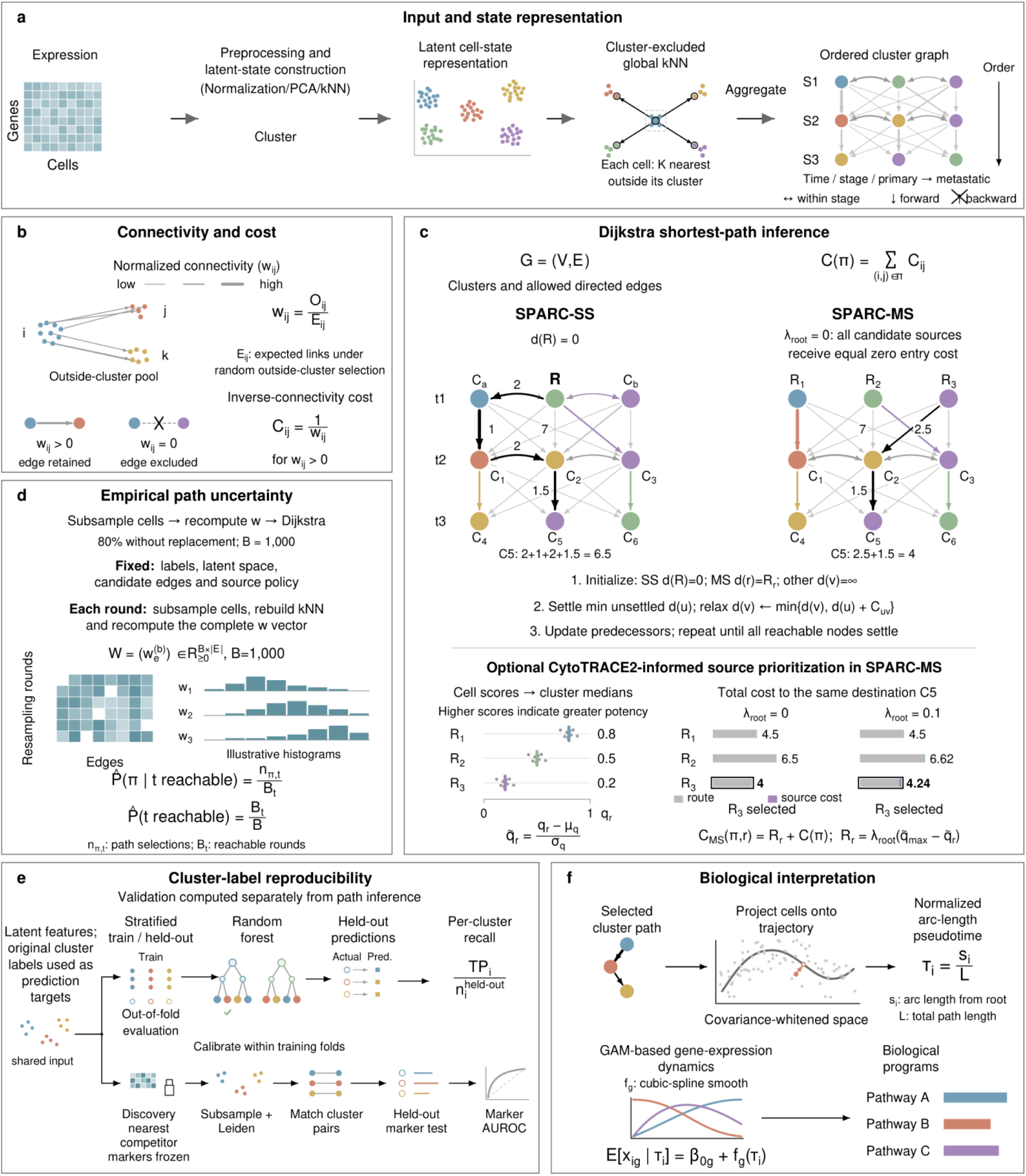
SPARC: a unified framework for trajectory inference within and across conditions. (a) Expression preprocessing, latent representation and cluster-excluded global neighbor selection produce the ordered cluster graph. (b) Connection strength w is the observed-to-expected number of union links between clusters; positive connections receive cost C = 1/w. (c) Schematic illustration of Dijkstra inference in single-source SPARC (SPARC-SS) and multi-source SPARC (SPARC-MS). The main multi-source analysis gives all earliest-condition clusters zero entry cost. The lower example illustrates an optional CytoTRACE2-based finite source preference. (d) Within-cluster cell subsampling, complete-vector connection-strength recalculation and exact Dijkstra inference over 1,000 rounds. Reachability uses all rounds and conditional path frequency uses reachable rounds. (e) Separate cluster assessments use held-out random forest (RF) recall and frozen nearest-competitor marker area under the receiver operating characteristic curve (AUROC) after subsampling and reclustering. (f) Projection onto a covariance-whitened cluster route supplies normalized arc-length pseudotime for generalized additive model (GAM) expression dynamics and pathway analysis.

SPARC takes molecular profiles, cluster assignments and a specified ordering of conditions as input. Conditions can represent collection times, anatomical sites or other ordered experimental states. Following expression preprocessing, cells are represented in a shared latent space. Each cell selects its nearest neighbors from the pooled population while excluding members of its own cluster. This cluster-excluded global neighborhood search provides a consistent reference for connections among clusters within the same condition and across different conditions. For any pair of clusters, SPARC compares the observed number of unique neighbor links with the number expected given the cluster sizes and eligible neighbor pools. The resulting observed-to-expected ratio defines a connection strength, w (Fig. 1b).

Biological ordering determines which connections can contribute to a route. Connections within a condition are allowed in either direction, whereas connections across conditions are directed from earlier to later groups. In a primary-to-lung analysis, for example, the model permits progression within the primary lesion, a transition from primary to lung, and further within-lung progression. It excludes lung-to-primary transitions under that prescribed ordering. Each positive connection is assigned the inverse cost, C = 1/w, so a stronger connection contributes a smaller cost. Dijkstra’s algorithm identifies the minimum-total-cost path from a designated source to each reachable destination (Fig. 1c). A candidate route is thus evaluated as a complete sequence of connections, including any primary or metastatic intermediates.

SPARC supports two source-selection settings. In single-source SPARC (SPARC-SS), the investigator specifies one initial state and reconstructs routes from that state to other reachable populations, including destinations within the starting condition. This setting is useful when a biologically motivated origin is available and local progression from that origin is part of the question. In multi-source SPARC (SPARC-MS), every cluster in the earliest condition is eligible to initiate a route. The principal SPARC-MS analysis assigns equal starting cost to all candidates and selects the source and complete path with the lowest total cost for each later destination. An optional CytoTRACE2-informed source preference can be included when developmental-potential estimates provide a relevant prior^20^. This preference changes the starting costs but does not exclude candidate sources.

To examine why specifying a starting population is insufficient to constrain progression, we inspected the paired mouse primary–lung profiles. Cell-state maps identify the populations at both sites (Extended Data Fig. 1a). The Monocle 3 and Slingshot trajectories included primary-to-lung transitions followed by lung-to-primary transitions, as identified from the root-to-endpoint path lists. The corresponding UMAP projections illustrate these inferred paths (Extended Data Fig. 1b). Specifying a primary origin alone does not impose the primary-to-lung direction on every subsequent transition.

The framework also supports analyses of molecular change along a selected route. Cells belonging to its constituent clusters are projected onto a piecewise-linear path^21^ in a covariance-whitened latent space^22^, and their projected positions define a route-specific pseudotime (Fig. 1f). Generalized additive models (GAMs) identify expression changes and where they occur along that ordering^21,23,24^, with false discovery rate (FDR) correction within each trajectory^25^. Direction-specific gene sets and pathway annotations characterize where molecular programs emerge along each inferred trajectory^26,27^. These analyses relate molecular programs to the states included in each route.

We checked how the routes changed when different cells were sampled^8,11,28^ and whether their clusters could still be distinguished from similar populations (Fig. 1d,e). For each cell subsample, we recalculated all connection strengths together and reconstructed the routes. Separately, held-out label prediction and nearest-competitor marker preservation tested the expression-defined clusters. The first assessment measures route stability under cell sampling; the cluster assessments test how well the original labels and marker differences are preserved. Methods and Supplementary Methods describe these tests, and Extended Data Fig. 2c–e shows the sampling-fraction analyses and cluster-reliability assessments.

### SPARC reconstructs directed trajectories supported by independent lineage evidence

We began with conventional time-course datasets because their ordered reference structures provide a direct test of whether a method recovers both the correct connections and their directions. Using Splatter^29^, we simulated linear, single-bifurcation, double-bifurcation and asymmetric branching trajectories across easy, standard and hard expression settings (Fig. 2a). We assessed directed-edge F1, which penalizes a reversed connection even when the same two states are linked. SPARC recovered nearly all reference directions across these structures, while comparator performance varied with topology and expression difficulty (Fig. 2b). Graph-similarity analyses^8,30^ gave a complementary view of structural recovery (Extended Data Fig. 2b). At smaller cell numbers, recovery was less consistent, indicating that adequate sampling remains important for representing intermediate populations (Extended Data Fig. 2a).

**Figure 2.**
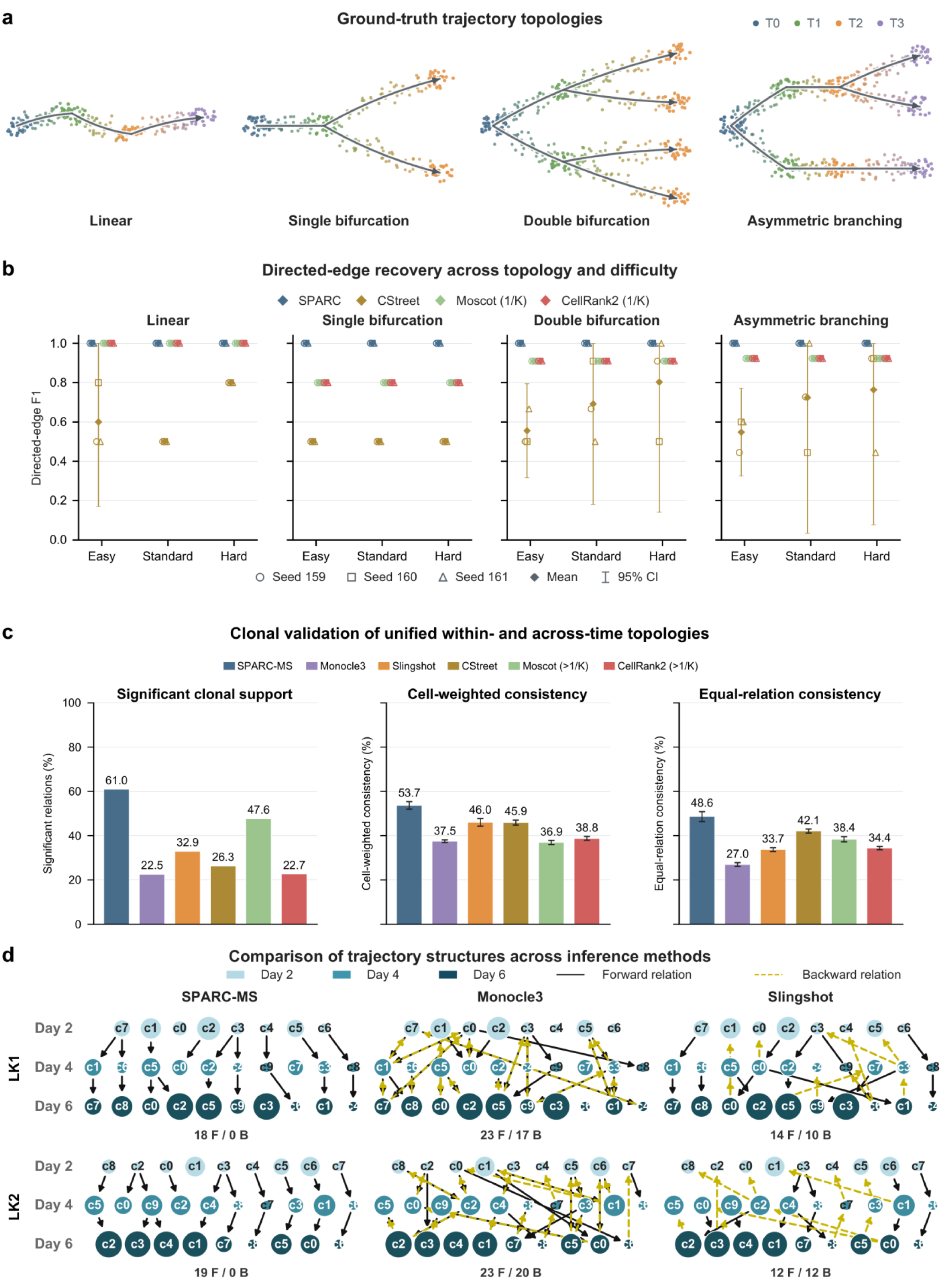
General trajectory recovery and independent LARRY clonal support. (a) Four time-ordered reference trajectory topologies. (b) Directed-edge F1 (the harmonic mean of precision and recall) for SPARC, CStreet, moscot and CellRank 2 across four topologies and three difficulty settings. Small symbols show three generation seeds; diamonds show means and intervals show t-based 95% confidence intervals bounded by the metric’s [0,1] range. (c) Pooled significant clonal support, cell-weighted consistency and equal-relation consistency for complete native outputs from six methods in LK1, LK2 and LSK. LK1 and LK2 denote culture-series identifiers; LSK denotes lineage-negative, Sca1-positive, Kit-positive progenitors. Cell-weighted and equal-relation error bars show sample standard deviation (SD) across five clone-evaluation partitions. Relation counts and within-day availability differ among methods; backward relations receive zero directional support. moscot and CellRank 2 reporting thresholds are specified in Methods. (d) Multi-source SPARC (SPARC-MS), Monocle 3 and Slingshot relation graphs for LK1 and LK2. Node size denotes cell number and color denotes sampling day; solid black and dashed yellow lines denote forward and backward relations. F and B give their counts. LSK maps and experiment-specific clonal comparisons are shown in Extended Data Fig. 3a,b.

Known simulated structures provide a useful reference, but real transcriptional similarities need independent biological evaluation. We therefore turned to the LARRY hematopoietic datasets, which jointly measure expression and heritable lineage barcodes at days 2, 4 and 6^18^. Two experiments, LK1 and LK2, share a day-2 origin before following separate cultures; LSK begins from a distinct progenitor population. We reconstructed trajectories using expression and sampling information only. The lineage barcodes were withheld until after inference and used to test whether the populations placed along inferred trajectories showed enriched clonal overlap.

To evaluate whether the relationships inferred by different methods were consistent with independent clonal information, we compared SPARC-MS, Monocle 3, Slingshot, CStreet, moscot and CellRank 2 using fixed reporting clusters (Fig. 2c). We retained the connections reported by each method, including within-day connections where available and directed across-day connections. Within-day pairs were evaluated as unordered relationships, whereas across-day connections were assessed in their inferred direction; backward-day connections received no directional support. Because these methods report different numbers and types of connections, clonal support was assessed within each method’s own output and interpreted in light of these differences.

Across LK1, LK2 and LSK, SPARC achieved the highest proportion of significantly clone-enriched relationships among all six evaluated methods: 36 of its 59 selected relations (61.0%) showed significant enrichment (Fig. 2c). The experiment-specific proportions were 50.0%, 57.9% and 75.0%, respectively (Extended Data Fig. 3b). These results indicate strong clonal support within SPARC’s selected graph.

To assess clonal consistency under different weighting schemes, we used cell-weighted and equal-relation summaries. The first emphasizes informative cell incidences, whereas the second gives each selected relation equal weight. SPARC also achieved the highest consistency among all six methods under both definitions, with pooled values of 53.7% and 48.6%, respectively (Fig. 2c). The two scores differ because highly represented clonal relationships contribute more heavily to the first summary. In the inferred directed graphs, Monocle 3 and Slingshot included backward-day relations in all three LARRY experiments, while SPARC-MS followed the prescribed sampling order (Fig. 2d; Extended Data Fig. 3a). Thus, independent barcodes and explicit direction checks provided complementary evidence about the selected state relationships.

We next examined whether the reported paths were sensitive to cell retention, cluster distinction or starting-source preference. The most frequent routes remained similar when 70%, 80% or 90% of the cells in each cluster were retained (Extended Data Fig. 2c). Most clusters also showed high held-out label recall and preservation of markers that distinguished them from their nearest competitors (Extended Data Fig. 2d,e). A modest optional CytoTRACE2 source preference^20^ preserved 57 of the 59 most frequent SPARC relations or routes reported in this sensitivity comparison and did not materially change the principal interpretation (Extended Data Fig. 3c). The main benchmarks used equal source costs.

The time-course evaluations tested recovery of directed branching structures and agreement between inferred state relationships and independent clonal measurements. Metastatic progression poses a further challenge because complete routes may include several primary-tumor intermediates, a cross-site transition and additional changes within the metastatic lesion. These alternatives can also be represented by very different numbers of cells. To determine whether SPARC recovers the correct precursor and complete route under these conditions, we designed a second benchmark with known progression topologies and metastatic descendant allocations.

### SPARC improves recovery of metastatic precursors and complete progression routes across simulated and lineage-traced tumor datasets

Different primary-tumor populations may contribute unequally to metastatic dissemination, resulting in imbalanced representation of alternative progression routes and potentially complicating the recovery of their precursor and intermediate states. To evaluate whether SPARC can accurately recover progression trajectories under different seeding patterns, we generated primary-to-lung datasets using two reference lineage architectures: branched metastatic progression and a deep asymmetric cascade (Extended Data Fig. 4a). Each architecture contained four primary-to-lung routes. In each lineage architecture, we examined two metastatic seeding patterns (Fig. 3a): homogeneous seeding, in which the four routes contributed equally to the lung population, and heterogeneous seeding, in which their contributions were unequal, with some routes producing substantially more metastatic descendants than others. By varying metastatic seeding while preserving the underlying lineage topology, we tested whether SPARC could recover the correct precursor states, intermediate relationships and complete progression routes despite differences in metastatic descendant abundance.

**Figure 3.**
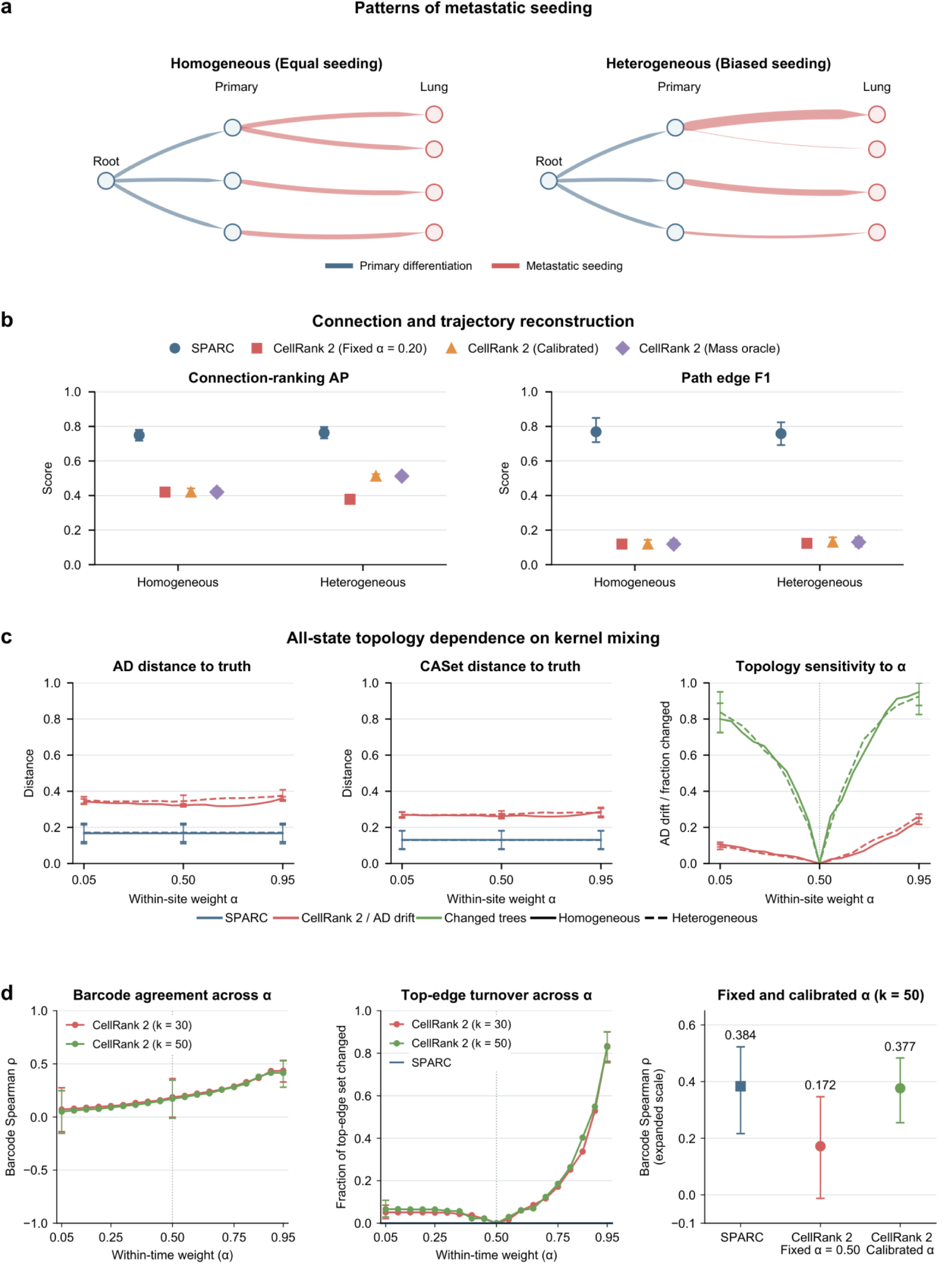
Recovery of mixed within- and cross-compartment progression. (a) Schematic illustration of equal and biased allocation among four primary-to-lung routes; red-arrow widths represent normalized generated Lung descendant abundance. (b) Held-out connection-ranking average precision (AP) and directed root-to-Lung-terminal path edge F1 for Homogeneous and Heterogeneous. SPARC is compared with CellRank 2 using k = 50 and fixed alpha = 0.20, calibration-selected alpha or a mass-truth oracle. The mass oracle selects alpha to minimize within-primary mass error and evaluates AP/F1 at that same setting. (c) All-state ancestor– descendant (AD) and common-ancestor-set (CASet) distance to truth across alpha, together with AD drift and the fraction of exact decoded trees differing from their alpha = 0.50 reference. Lower truth distance denotes closer reconstruction; drift and changed-tree fraction quantify sensitivity to the mixing weight. The simulations comprise 256 complete 8,000-cell/2,000-gene datasets: 96 calibration datasets from seeds 159, 161 and 162, and 160 test datasets from seeds 164–168. Reconstruction means and 95% intervals use five independent test-seed factorial means and 10,000 seed-bootstrap resamples. (d) Tumor Cas9 lineage validation across 14 units from 11 families, with k = 30/k = 50 sensitivity curves and the k = 50 fixed and family-held-out calibrated comparison. Intervals use 10,000 family-cluster bootstrap resamples.

To test whether performance depended on a particular model of RNA generation, we implemented the reference designs using PROSSTT, Splatter, TedSim and dyngen^29,31–33^. The resulting factorial benchmark contained 256 complete datasets spanning two topologies, two descendant-allocation settings, two expression regimes and eight random seeds for each simulator. We reserved 96 datasets generated with three seeds for calibration and evaluated reconstruction on 160 held-out datasets generated with five seeds. All methods received the complete expression inputs, supplied state labels and a shared expression representation. The lineage topology provided ground truth for testing connections, complete routes and ancestral relationships.

To determine whether a method recovers the complete progression sequence, we evaluated three complementary aspects of reconstruction. Connection-ranking average precision tested whether true state connections were scored above incorrect ones. Directed root-to-lung-terminal path edge F1 measured recovery of complete sequences and their directions. Ancestor–descendant and common-ancestor-set distances assessed how accurately states were positioned within the reconstructed topology^34,35^. CellRank 2 uses a mixing parameter to control the relative contributions of within-condition and cross-condition transitions, which may influence the inferred trajectory topology. To ensure a fair and rigorous comparison, we evaluated CellRank 2 under increasingly favorable parameter-selection settings. In addition to using a fixed mixing weight, we optimized this parameter using separate simulated datasets with known transition patterns. We further gave CellRank 2 access to the true within-primary transition mass of each test dataset, allowing it to select the mixing weight that best reproduced this information. This oracle setting provides an advantage unavailable in real applications, and allowed us to evaluate how well CellRank 2 could reconstruct complete progression routes even with privileged information for parameter selection.

Despite this advantage, SPARC showed higher connection-ranking precision and substantially better recovery of complete routes in both metastatic-allocation settings (Fig. 3b). In homogeneous datasets, connection-ranking average precision was 0.749 for SPARC and 0.420 for calibrated CellRank 2; directed path edge F1 was 0.769 and 0.119, respectively. In heterogeneous datasets, the corresponding average precision values were 0.764 and 0.512, and path F1 values were 0.758 and 0.131. The advantage was also apparent for links within the heterogeneous lung compartment, where average precision reached 0.804 for SPARC and 0.658 for the calibrated comparator. Together, these results demonstrate SPARC’s advantage in recovering the underlying topology of metastatic progression, including the correct placement of precursor and intermediate states along complete primary-to-lung trajectories, even under favorable parameter-selection settings for CellRank 2.

We next asked whether errors in individual connections affected the ancestral organization of the reconstructed state graph. Across all 160 test datasets, the mean ancestor–descendant distance to truth was 0.169 for SPARC and 0.342 for calibrated CellRank 2; the corresponding common-ancestor-set distances were 0.130 and 0.269 (Fig. 3c; Extended Data Fig. 4c). Lower distances indicate that SPARC more closely recovered which states lay upstream of others and which states shared ancestors in these simulated lineages. This matters for metastatic studies because a change in the inferred ancestral position of a primary state can alter which population is nominated for lineage tracing or perturbation.

We next examined how the mixing parameter affected CellRank 2’s inferred progression topology. Across 19 tested values, 158 of 160 test datasets showed changes in their decoded state trees, with corresponding shifts in the inferred ancestor–descendant relationships (Fig. 3c). Despite exploring a wide range of parameter settings, CellRank 2 consistently showed larger topological distance from the known reference than SPARC.

Because simulated truth may not capture every feature of metastatic evolution, we further assessed agreement with independent tumor lineage-tracing data. We analyzed 14 primary-metastasis units from 11 tumor families across KP-Tracer, macsGESTALT and Quinn xenograft datasets^1–3^. Expression-derived state connections were compared with Cas9-derived lineage scores within primary tumors, across primary and metastatic compartments, and within metastases (Fig. 3d). SPARC achieved a family-weighted mean Spearman correlation of 0.384, substantially exceeding CellRank 2 with a prespecified mixing weight of 0.50 (0.172). When CellRank 2’s mixing parameter was optimized using lineage information from other tumor families, its correlation increased to 0.377, approaching SPARC’s performance. Thus, SPARC achieved comparable lineage agreement without requiring the lineage-informed parameter tuning used to improve CellRank 2.

The simulations and lineage experiments answer complementary questions. A reference lineage allows us to measure whether the inferred intermediate sequence and ancestral topology are correct. Clone sharing in observed tumors assesses whether expression-derived relationships are compatible with independent lineage information. We next asked whether SPARC could reveal how molecular programs emerge and diverge along progression routes from primary tumors to metastatic lesions.

### SPARC distinguishes local progression from metastasis-associated transcriptional programs in osteosarcoma

To investigate this question in a biologically relevant model of spontaneous metastasis, we generated paired scRNA-seq profiles of primary osteosarcoma tumors and pulmonary metastases from the same immunocompetent mice in a syngeneic orthotopic model^19^. To our knowledge, this represents the first paired primary–spontaneous metastatic mouse osteosarcoma scRNA-seq dataset. Because lung metastases arise through dissemination from the primary bone tumor, these paired samples provide a valuable setting for investigating molecular progression from primary-tumor states to metastatic populations. We combined cells from three mice to reconstruct primary-local and metastatic trajectories within a shared expression-state representation, while retaining mouse identity as the biological replicate (Fig. 4).

**Figure 4.**
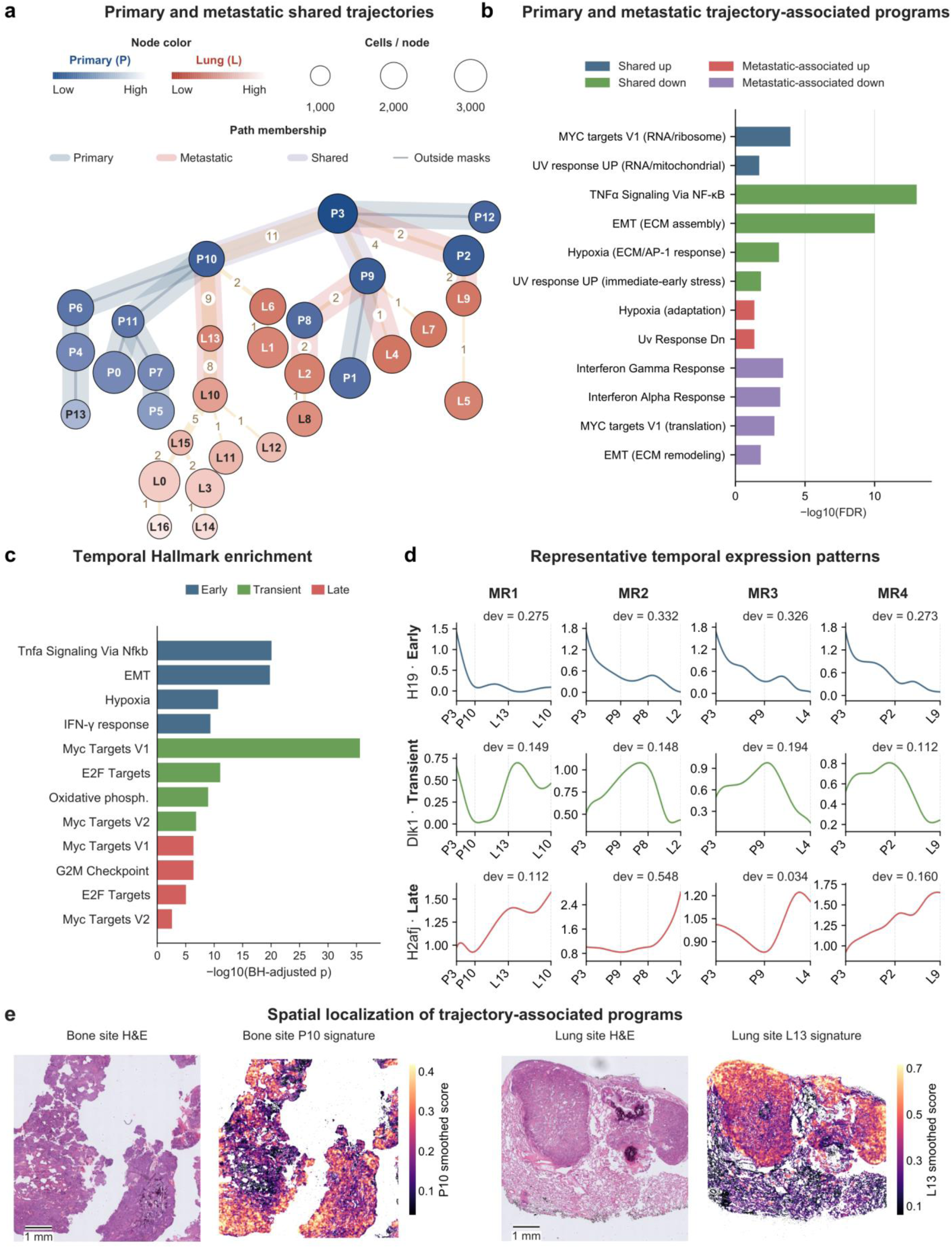
Local and metastatic progression programs and their route dynamics. (a) P3-rooted primary/lung trajectory map with 31 nodes and 30 union edges. P and L label primary and lung clusters, respectively. Node area denotes cell number and color denotes site-specific pseudotime. Edge widths and labels count displayed destination paths using each edge. Masks mark primary-local, metastatic-trunk and shared path membership. Primary-local program comparisons use endpoints P0, P13, P5, P1 and P12; metastatic trunks end at L10, L2, L4 and L9. Four representative metastatic routes (MR1–MR4) span four distinct primary-to-lung transition states. (b) Hallmark enrichment in shared increasing/decreasing genes and metastatic-associated genes across four metastatic trunks and five primary-local routes. Labels in parentheses were assigned by the authors to distinguish the functional roles of different gene subsets contributing to the same enriched pathway. UP and DN retain their original Hallmark gene-set designations; observed expression directions are indicated by the plotted categories. Enrichment uses the filtered ranked lists described in Methods. (c) Hallmark enrichment among early, transient and late gene-expression classes concordant across the four selected metastatic trunks, shown as −log10 Benjamini-Hochberg (BH)-adjusted P. (d) Generalized additive model (GAM) curves for H19, Dlk1 and H2afj across MR1-MR4, illustrating early, transient and late patterns. Labels give explained deviance; cluster positions mark the ordered route states. (e) Registered H&E images alongside ortholog-matched mouse P10 and L13 signature scores in patient Visium HD specimens Bone (primary bone tumor) and Lung (pulmonary metastasis). Each H&E image and its corresponding signature map share the same tissue field, orientation and spatial scale. These specimens are from different patients; color denotes spatially smoothed signature score. The Bone field shows the primary tumor. In the Lung field, compact tumor nodules occupy the upper part of the section, with alveolar lung tissue below and between the nodules. The corresponding Lung zonation is shown in Fig. 5c. H&E scale bars, 1 mm.

We first reconstructed candidate trajectories using malignant-cell states from both the primary tumor and lung lesions. CytoTRACE2 estimates were used to nominate a primary starting state, P3, as the root node^20^. From this starting point, SPARC reconstructed trajectories toward different primary-tumor states and lung metastatic populations, allowing us to compare molecular programs associated with local tumor progression and metastasis.

To identify molecular programs associated with metastatic progression, we compared gene-expression changes shared across primary-local and metastatic trajectories with those preferentially observed along metastatic routes. Both types of progression showed enrichment of MYC-related RNA-processing and ribosome-associated programs^36^ among increasing genes, while TNFα/NF-κB signaling and EMT-associated programs were enriched among decreasing genes (Fig. 4b). A distinct MYC-target subset associated with cytosolic translation decreased preferentially along metastatic routes. These shared changes suggest that some transcriptional programs associated with metastatic progression also accompany local tumor diversification. Hypoxia-related enrichment resolved into functionally distinct components. Shared decreasing genes represented ECM-associated responses and AP-1-mediated stress responses^37^, whereas metastatic-associated increasing genes included a CITED2-containing component associated with HIF feedback regulation and adaptation to hypoxic stress^38^. This pattern suggests a shift in the composition of hypoxia-related responses during metastatic progression. Metastatic routes also showed decreases in interferon-response programs. These findings are consistent with previous studies implicating MYC and hypoxia signaling in osteosarcoma progression^6,19^ and suggest that metastatic progression involves additional changes in stress adaptation and immune-related transcriptional programs. By comparing metastatic trajectories against local primary-tumor progression, SPARC distinguishes broader malignant-state changes from molecular programs preferentially associated with progression toward lung metastases.

We next examined the temporal patterns of gene expression changes along the inferred metastatic routes. Inflammatory and stress-response programs were enriched among genes expressed at early pseudotime positions (Fig. 4c), consistent with previous studies highlighting inflammatory signaling and cellular plasticity in metastasis-initiating osteosarcoma populations^6,39^. MYC, E2F and oxidative-phosphorylation programs showed transient enrichment, followed by stronger MYC, E2F and G2M activity at later positions, suggesting changes in metabolic and proliferative programs along metastatic progression. These patterns are consistent with previous evidence of transcriptional and metabolic plasticity during osteosarcoma colonization^6^.

This ordering helps formulate more specific experimental questions. A program enriched in early primary-route states could be tested in a candidate primary precursor, while one associated with later lung-route states could be investigated for a potential role in metastatic adaptation. We also compared predicted cell-cell communication^40^ during local and metastatic progression. Several sender and receiver cell types were shared, but the associated ligand and receptor genes differed (Extended Data Fig. 5c). Thus, apparently similar cell-type interactions may involve distinct candidate signals along the inferred progression contexts.

To validate the biological relevance of the mouse-derived progression programs in human osteosarcoma, we generated patient-derived Visium HD profiles of primary bone tumors and pulmonary metastases, constituting, to our knowledge, the first patient-derived osteosarcoma Visium HD dataset. We transferred the SPARC-derived mouse signatures to their human orthologs and examined their spatial expression in human tissues. Signatures associated with primary and metastatic cell states exhibited spatially structured distributions in the corresponding human specimens (Fig. 4e; Extended Data Fig. 5b,d), providing independent cross-species support for the molecular programs identified along the inferred mouse trajectories. Building on this cross-species validation, we next investigated whether programs associated with different metastatic routes occupied distinct spatial regions and cellular microenvironments in human tumors.

### SPARC identifies route-dependent tumor programs with distinct spatial distributions

The four metastatic routes differed in their inferred primary precursors, intermediate states and lung destinations. These differences raise a further biological question: do alternative metastatic trajectories show similar molecular changes, or are particular programs associated with distinct progression routes? We compared the selected MR1– MR4 trajectories at two levels, considering the overall programs represented along each route and the changes that occurred as cells advanced through its constituent states (Fig. 5a,b).

**Figure 5.**
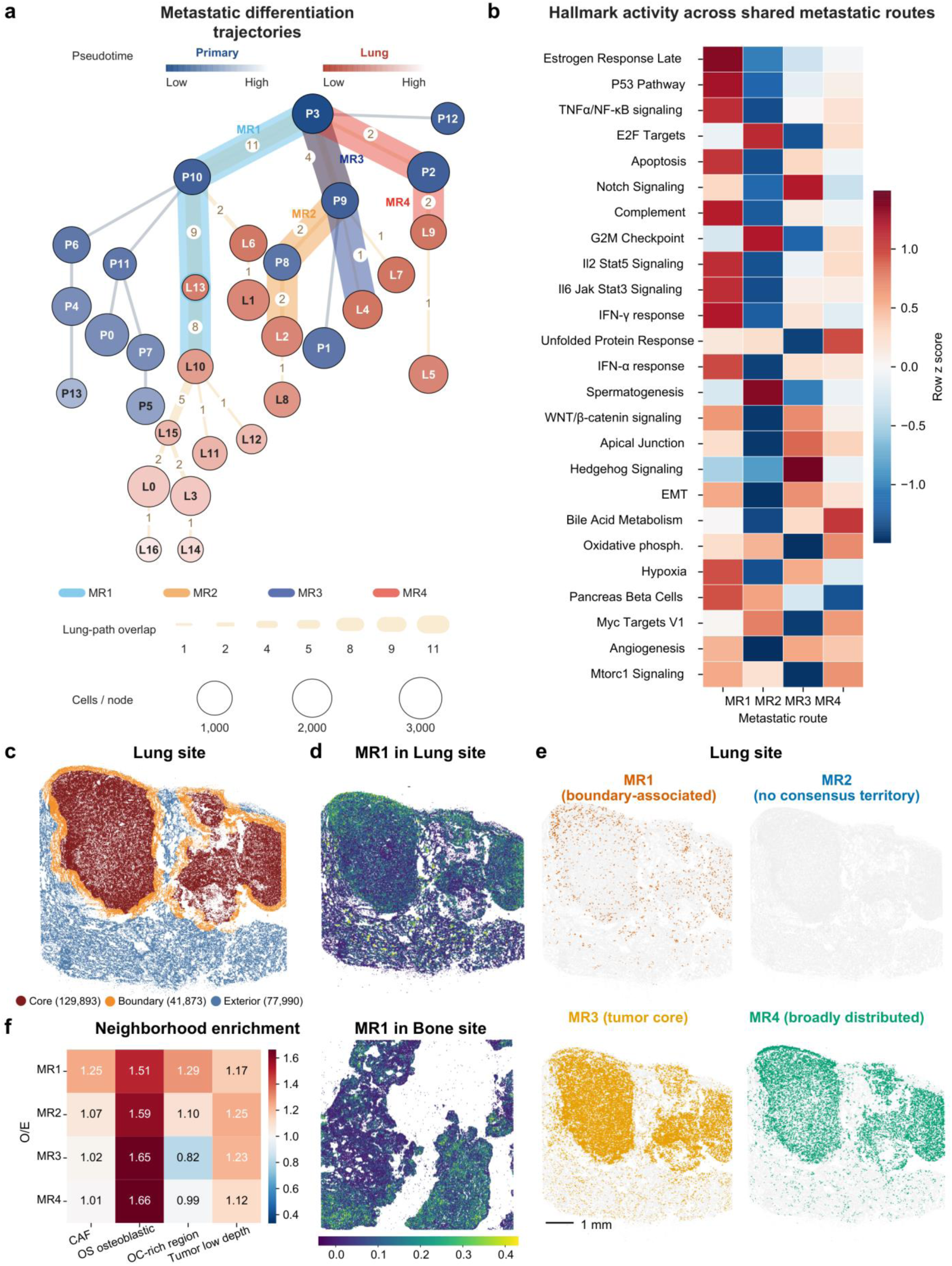
Metastatic route programs occupy distinct spatial environments. (a) The four selected routes MR1–MR4 highlighted on the P3-rooted graph, with node size indicating cell number and pale edge widths indicating lung-destination path overlap. (b) Mean Hallmark activity across the four selected trunks, standardized within each row. (c) Core, boundary and exterior zones in the Lung specimen. (d) Continuous MR1 signature scores in Lung and Bone. (e) Route-associated Lung territories: MR1 is boundary-associated, MR3 concentrated in the core and MR4 broadly distributed; MR2 has no consensus territory under the display rule. The corresponding registered H&E is shown in Fig. 4e. Scale bar, 1 mm. (f) Observed-to-expected abundance of annotated segmented objects in route-associated spatial neighborhoods. OC-rich region denotes the osteoclast-rich region. Signatures were defined from relative expression among mouse routes and transferred by ortholog matching. Bone and Lung represent different patients.

Average pathway activities differed among routes for proliferative, developmental and oxidative-metabolic programs, alongside recurring features of extracellular matrix remodeling, hypoxia, angiogenesis and inflammation (Fig. 5b). Because a high route-average score does not establish that a program increases during progression, we also evaluated gene-expression changes as a function of route-specific pseudotime. This distinction separated programs characteristic of the populations comprising a route from programs that changed systematically along its inferred sequence (Extended Data Fig. 5a).

Across the four routes, 770 genes showed strong significant dynamics in a common direction, 560 showed preferential dynamics in one route, and 1,526 showed opposing directions among routes. Mdk and Ptn decreased along all four selected routes, while Spp1 increased along MR1 and MR4 but decreased along MR2 and MR3. Ly6c2 showed preferential dynamics along MR1. These patterns demonstrate how pooling all primary and metastatic populations can obscure changes that follow different directions in alternative candidate trajectories. In particular, opposing trends can offset one another in a site-level average even when each route shows a consistent progression-associated pattern.

To examine the spatial distributions of these metastatic programs in human tumor tissue, we extended our cross-species analysis to the four mouse-derived route signatures. Using patient-derived Visium HD profiles, we found that the corresponding malignant programs occupied different regions of the human pulmonary metastasis (Fig. 5c–e). MR1-associated programs were enriched near the tumor boundary, MR3-associated programs were concentrated in the tumor core, and MR4-associated programs showed a broader distribution. MR2 lacked a consensus spatial territory under the predefined assignment criteria. MR1-associated activity was also detected in the primary bone tumor specimen from a different patient (Fig. 5d). These findings extend the mouse-derived transcriptional differences to spatially distinct malignant-cell programs in human osteosarcoma.

The spatial segregation of these programs prompted us to investigate whether they were associated with different cellular microenvironments. MR1-associated malignant cells showed greater spatial association with the osteoclast-rich region, whereas MR3-associated cells showed a contrasting neighborhood profile (Fig. 5f). Given previous reports of osteoclast-like populations in osteosarcoma lung metastases^41,42^, these findings suggest that the association with an osteoclast-rich microenvironment may differ among metastatic malignant states. We therefore examined myeloid-state progression and candidate tumor–myeloid signaling interactions to investigate the potential mechanisms underlying this heterogeneity.

### SPARC reveals heterogeneity in osteoclastogenic associations across metastatic trajectories

A fundamental question in osteosarcoma metastasis is how malignant cells originating in bone survive and expand after colonizing the lung, where they encounter a fundamentally different microenvironment. One possibility is that metastatic tumor cells remodel their surrounding microenvironment to create conditions favorable for their continued survival and progression. Classical osteoclasts reside on bone surfaces and mediate bone resorption during skeletal remodeling^43^. However, osteoclast-like cells (OLCs) have also been identified outside the skeletal compartment, including in chronic pulmonary injury, where local myeloid populations acquire osteoclast-associated features^44^. Previous single-cell and spatial transcriptomic studies have further identified osteoclast-associated myeloid populations at different differentiation stages in osteosarcoma lung metastases^41,42^. These observations raise the question of how ectopic osteoclastogenesis develops within pulmonary metastases and whether particular malignant-cell populations contribute to this process. Motivated by the distinct osteoclast-rich neighborhoods associated with the metastatic programs identified in Figure 5, we used SPARC to investigate candidate myeloid differentiation trajectories and their potential interactions with malignant cells.

To investigate how osteoclast-like states may arise in the lung metastatic microenvironment, we first examined the myeloid populations identified in mouse and human osteosarcoma. Canonical osteoclast markers and resorption-associated programs supported the identification of osteoclast-like populations in mouse primary tumors and lung metastases (Extended Data Fig. 6a,b). Reanalysis of published human osteosarcoma scRNA-seq data^41^ further identified corresponding populations in lung metastases, with elevated fusion, resorption and NFATC1-associated programs compared with monocytes and macrophages (Extended Data Fig. 6c,d). These findings provided the molecular basis for investigating possible differentiation relationships among the observed myeloid states.

To reconstruct candidate osteoclast-associated differentiation, we applied SPARC to the mouse lung myeloid populations. SPARC identified a progression route from monocyte-like cells through a macrophage-like intermediate to an osteoclast-like state (Mono1 → Mφ3 → OLC; Fig. 6a). Along the inferred trajectory, early AP-1-associated activity was followed by increased Nfatc1 expression and the emergence of fusion, acidification and matrix-resorption programs. This transcriptional ordering is consistent with established mechanisms of osteoclast differentiation, in which NFATc1 coordinates osteoclast commitment and maturation^45,46^. To examine whether corresponding differentiation-associated programs were present in human metastases, we mapped mouse-derived myeloid signatures onto patient Visium HD profiles. Monocyte/inflammatory, macrophage-intermediate, pre-osteoclast-like and mature-resorptive programs showed distinct spatial distributions in the human lung metastatic tissue (Fig. 6b), providing cross-species support for the biological relevance of the inferred myeloid progression.

**Figure 6.**
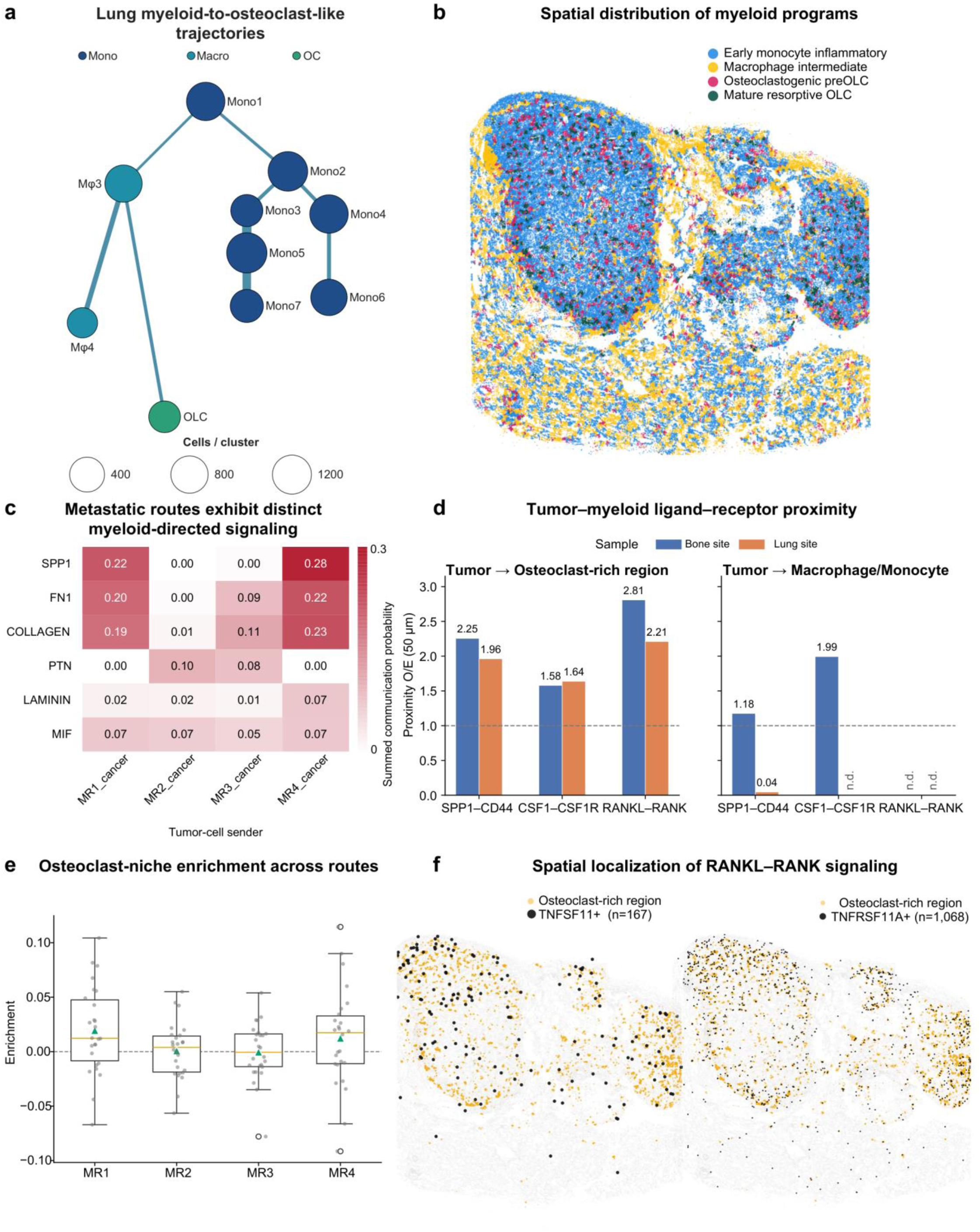
Myeloid progression and candidate signals in an osteoclast-rich niche. (a) Mouse lung single-source SPARC (SPARC-SS) topology containing Mono1 → Mφ3 → osteoclast-like cell (OLC). Node area denotes cell number and color denotes monocyte, macrophage or osteoclast-like annotation. Downstream analysis retained 1,975 route cells: 1,053 Mono1, 801 Mφ3 and 121 OLC. (b) Lung spatial distributions of early monocyte/inflammatory, macrophage-intermediate, pre-OLC and mature-resorptive signatures from the broader four-program myeloid reference. These signatures are defined in Methods. (c) Predicted tumor-route to myeloid communication probabilities for SPP1, FN1, collagen, PTN, laminin and MIF. (d) Tumor-to-myeloid ligand-receptor proximity at 50 µm in Bone and Lung for SPP1-CD44, CSF1-CSF1R and RANKL-RANK, with receiver groups comprising the osteoclast-rich region and macrophage/monocyte annotations; n.d. denotes an unavailable measurement. These specimens are from different patients. (e) Canonical osteoclast-niche enrichment for MR1-MR4 in 26 conventional Visium sections from 19 patients in external GSE293065. Gray dots denote sections, boxes show median and interquartile range, whiskers extend to 1.5 times that range and triangles show means. (f) Lung distribution of segmented objects assigned to the osteoclast-rich region, together with TNFSF11- and TNFRSF11A-expressing objects.

Having identified a candidate route toward osteoclast-like differentiation, we next investigated whether malignant cells associated with different metastatic trajectories expressed distinct signals potentially involved in this process. Comparing the four metastatic programs revealed stronger inflammatory, matrix-remodeling and predicted myeloid-directed communication signals along MR1 and MR4 than along MR2 and MR3 (Fig. 6c; Extended Data Fig. 7a,b). Among the candidate interactions, SPP1–CD44, CSF1–CSF1R and RANKL–RANK were particularly relevant because of their established roles in osteoclast-associated motility, myeloid survival and differentiation^45,47–50^. These differences suggest that metastatic malignant populations may vary in their potential to influence the local myeloid environment.

We further examined whether candidate osteoclastogenic signaling differed between the primary and metastatic sites. CellChat analysis^40^ identified RANKL–RANK communication at both sites, but with distinct predicted cellular sources: cancer-associated fibroblasts were the predominant senders in primary tumors, whereas malignant cells emerged as prominent predicted senders in lung metastases (Extended Data Fig. 8a). Osteoclast-like populations were the principal predicted receivers in both conditions. Consistent with these predictions, spatial analysis of human specimens identified ligand-expressing and receptor-expressing cells in proximity for SPP1–CD44, CSF1–CSF1R and RANKL–RANK interactions (Fig. 6d,f). The lung tumor boundary also contained a distinct macrophage compartment characterized by collagen and chemokine expression. Cross-species transcription-factor analysis revealed concordant changes in several osteoclast-associated regulatory programs between mouse and human myeloid states (Extended Data Fig. 8b). Together, these findings identify candidate molecular and spatial interactions through which metastatic tumors may influence osteoclast-associated myeloid states.

To determine whether these osteoclast-associated environments were preferentially linked to particular metastatic tumor programs, we compared the spatial neighborhoods of MR1- and MR3-associated malignant cells. MR1-associated populations showed stronger spatial associations with the osteoclast-rich region and mature-resorptive programs than MR3-associated populations (Extended Data Fig. 8c). We further assessed these relationships in an independent cohort comprising 26 spatial transcriptomic sections from 19 osteosarcoma patients^42^. MR1 showed stronger associations with macrophage/monocyte-rich regions and canonical osteoclast-associated programs than MR3 across most sections (Fig. 6e; Extended Data Fig. 8d,e). The association remained detectable after accounting for broader macrophage signatures, although evidence based on the more restricted mature osteoclast-marker panel was weaker. These results support heterogeneity in the myeloid environments associated with distinct metastatic malignant programs.

Together, these analyses reveal that the association between metastatic osteosarcoma states and ectopic osteoclast-associated environments varies across progression routes. By integrating inferred malignant and myeloid trajectories with human spatial transcriptomics, SPARC identifies specific metastatic tumor programs associated with candidate osteoclastogenic signals and myeloid differentiation states. These findings suggest that certain metastatic malignant populations may preferentially support ectopic osteoclastogenesis, providing a new hypothesis for understanding how osteosarcoma cells interact with the pulmonary microenvironment. Functional studies targeting these tumor populations and their candidate signaling interactions could determine whether they contribute to osteoclast-like differentiation and metastatic persistence.

### SPARC extends trajectory inference to treatment-response settings and spatial transcriptomics

Chemotherapy response presents another biological setting in which reconstructing progression both within and across conditions is essential. Drug-resistant populations may arise through the selection of pre-existing cellular states or the acquisition of adaptive programs during treatment. Distinguishing these possibilities requires understanding how molecular programs vary among cells within each treatment condition and how these states are connected across treatment stages. To investigate this setting, we applied SPARC to longitudinal single-cell RNA-seq data from high-grade serous ovarian cancer (HGSOC) collected before and after neoadjuvant chemotherapy.^51^ SPARC reconstructed candidate trajectories connecting treatment-naive and post-treatment malignant populations while capturing progression within each condition (Extended Data Fig. 9a). We focused on a previously reported chemotherapy-associated C7 stress-response program, which was detectable before treatment and enriched after chemotherapy. Although this program showed an overall increase following treatment, trajectory-specific analysis revealed heterogeneous dynamics, including sustained and non-monotonic expression patterns (Extended Data Fig. 9b). Gene-level analysis further distinguished early, intermediate and late stress-response programs across individual trajectories (Extended Data Fig. 9c,d). These findings demonstrate how joint reconstruction of within- and across-condition progression can resolve the molecular states and transitions associated with treatment response, providing candidate stages at which resistance-associated programs emerge or become enriched.

We further investigated whether SPARC could extend beyond scRNA-seq to spatial transcriptomic data, where molecular profiles can be analyzed together with their spatial tissue context. Using five patient-matched osteosarcoma conventional Visium comparisons, including four primary-to-lung-metastasis pairs and one primary-to-local-relapse pair, we constructed spatially informed expression representations and applied SPARC to reconstruct patient-specific progression trajectories (Extended Data Fig. 10a).^42^ Mapping inferred pseudotime onto the corresponding tissue sections revealed spatially organized matrix-remodeling and immune-response adaptation trajectories, allowing molecular progression to be examined in its anatomical context (Extended Data Fig. 10b,c). Trajectory-associated gene dynamics identified changes in osteogenic-matrix, inflammatory, antigen-presentation and osteoclast-associated programs, with several patterns recurring across patients (Extended Data Fig. 10d,e). To further examine applicability across spatial resolutions, we independently applied SPARC to myeloid populations in conventional Visium and single-cell-resolution Visium HD data. Both analyses recovered candidate progression toward osteoclast-like states, while Visium HD additionally resolved an earlier monocyte-like state and localized the inferred differentiation-associated progression within metastatic lung tissue (Extended Data Fig. 10f,g). These results demonstrate that SPARC can incorporate spatially informed molecular representations to reconstruct biologically interpretable trajectories across spatial transcriptomic platforms and resolutions.

## Discussion

Understanding metastatic progression requires identifying possible primary precursors, the intermediate transcriptional states through which they may progress, and the molecular changes associated with colonization of distant organs. Paired single-cell datasets provide snapshots of these populations, but do not directly record the histories that produced them. SPARC addresses this inference problem by evaluating progression within each site and transitions between sites using a common graph-based objective and an explicitly specified biological ordering.

The methodological distinction becomes important when several sequences can reach the same metastatic population. One route may pass through multiple primary intermediates before crossing to the lung; another may connect to the lung sooner and traverse additional metastatic states. The preferred sequence depends on the relative strengths of all constituent connections. SPARC evaluates those connections against the same cluster-excluded global neighborhood reference and compares the accumulated costs of complete routes. The methodological gain is a coherent basis for comparing alternative precursor and intermediate states across the combined progression topology.

In osteosarcoma, a local primary-tumor reference changes the interpretation of programs associated with metastatic populations. The inferred metastatic routes contained changes also observed during progression within the primary tumor, together with additional changes involving hypoxia-related and interferon-response annotations. Increased MYC-related program activity alongside reduced interferon responses along metastatic routes raises the possibility that progressing malignant states become less readily recognized by the immune system. This places metastatic-state differences against the progression already occurring in the primary lesion. Shared changes may accompany malignant-state progression more broadly, whereas additional changes suggest a metastatic context for further investigation. Their occurrence could reflect selection of a pre-existing primary state or adaptation to the distant environment. Lineage-traced tumors have shown that primary expansion and phenotypic plasticity can both contribute to metastatic progression^2^. SPARC organizes the observed states into candidate sequences in which these alternatives can be investigated, allowing program differences to be interpreted in the context of a proposed precursor and its subsequent state changes.

The human spatial observations add a tissue context to this progression analysis. Detecting corresponding malignant programs in patient-derived tumors establishes where these programs occur, while their association with osteoclast-rich neighborhoods identifies an environment in which tumor–myeloid interactions may be relevant. Osteosarcoma-derived cell lines were previously shown to induce an osteoclast-like phenotype in monocytes^52^, raising the question of whether this interaction is reused or remodeled in lung metastases. Pulmonary osteoclast-like differentiation has also been demonstrated in silica-induced fibrosis^44^, showing that such a myeloid program can emerge outside bone in another disease setting. The candidate myeloid trajectory and spatial interfaces identified here therefore motivate investigation of local differentiation and tissue adaptation in the metastatic niche. Tumor-derived signals, a myeloid environment favoring the malignant state and a shared tissue response remain possible explanations of the association. Distinguishing them requires functional evidence, including the cellular source of the relevant signals and the activity of the osteoclast-like population. The full human primary and metastatic maps are from different patients; larger paired cohorts could test recurrence of these program–environment relationships within individual disease courses.

Several limitations follow from the data and the assumptions needed to reconstruct routes. Single-cell RNA sequencing measures molecular states at collection and does not directly record each cell’s prior transitions^2,18^. Rare primary precursors may not be sampled, intermediate states may have disappeared, and transcriptionally divergent descendants may lie far from their ancestral states. In such cases, any expression-based method must reconstruct a route from incomplete evidence. The assumed condition order is also part of the model. For the primary-to-lung application, SPARC permits primary-to-lung transitions and excludes lung-to-primary connections. This is appropriate for investigating the specified progression direction; studies of reciprocal seeding or other histories would require a different biological ordering or graph design^53^.

Orthogonal clonal information offers a natural way to strengthen reconstruction when transcriptional intermediates are sparsely sampled or have diverged. In a future extension of the graph formulation, barcode-derived clonal relationships could inform edge weights, favoring lineage-compatible connections among transcriptionally and directionally supported populations. Such information could help distinguish genuinely related populations from those that resemble one another because they share a molecular program. Joint RNA–DNA assays such as DEFND-seq could provide additional clonal constraints in settings where experimental barcoding is unavailable^54^. These measurements would complement the RNA-state map, while additional sampling would still be needed to characterize missing molecular intermediates. In the current RNA analyses, lineage information was reserved for independent evaluation instead of constructing the graph. An extension using clonal information during reconstruction could be assessed on held-out clones or independent experiments to determine how much it improves the supported state relationships.

SPARC provides a common computational basis for organizing observed states into candidate paths across ordered conditions. Its value is greatest when the biological question depends on complete routes that include local progression and transitions between groups. Applied to metastasis, this representation places candidate precursor states in the context of downstream transcriptional changes and the tissue environments in which corresponding programs are observed. Combined with broader sampling and independent lineage or functional measurements, such route-based analyses can help convert static molecular profiles into testable hypotheses about the processes underlying disease progression.

## Methods

### SPARC trajectory inference

To reconstruct routes that follow the experimental ordering, SPARC combines expression-derived connections with direction constraints specified for the study. In a primary-to-lung analysis, connections are allowed within each site and from primary to lung. In a time course, they are allowed within each sampled condition and toward the next time point. Expression determines the strength of each connection, and the experimental design determines its permitted direction. Lineage barcodes, clone assignments and barcode-derived fate labels were reserved for evaluation after preprocessing, clustering, graph construction, root selection and path inference.

After dataset-specific quality control (QC), expression was normalized and represented by principal component analysis (PCA). All included conditions were placed in a shared latent space. Harmony correction was applied when supported by the biological replication and batch structure^55^. For lineage-tracing datasets, correction was performed within each time point, and the corrected coordinates were recentered to the original time-specific PCA centroid. Leiden clustering was performed separately within each time or condition^56^. The joint cell-neighbor graph then connected cells across the retained conditions. The expression representation, cluster labels and experimental ordering were fixed for the path-uncertainty analysis.

To compare within- and cross-condition connections using the same neighborhood reference, we built a cluster-excluded global k-nearest-neighbor (kNN) graph. Each cell searched for neighbors among cells from all included conditions outside its own cluster. For every cell in cluster *i*, the *K* nearest cells outside cluster *i* were selected in the shared latent space. The union of directed selections counted each cell pair once if either cell selected the other. Let *N* be the total number of retained cells, *n_i_* and *n_j_* the retained sizes of two clusters, and *U_ij_* the number of unique cross-cluster cell pairs observed between them. Under a reference model in which different cells independently select the prescribed number of distinct neighbors uniformly from cells outside their own cluster,

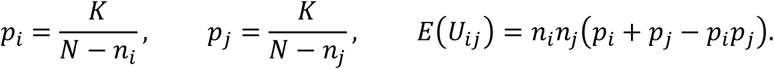

The connection strength was

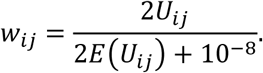

The factor of two accounts for symmetric adjacency in both observed and expected counts and cancels apart from the numerical stabilizer. PAGA introduced the graph-abstraction rationale for measuring connections between populations relative to their random expectation^16^. SPARC calculates an uncapped, exclusion-aware ratio from neighbor selections outside each cell’s own cluster. All cluster pairs permitted by the experimental ordering are retained as candidates, including pairs with a full-data weight of zero. Within-condition connections are reciprocal; cross-condition connections point forward. Positive finite weights yield traversable edges with inverse-strength costs. A zero weight is saved in the complete uncertainty vector, and its edge is unavailable in that round.

### Single-source and multi-source inference

SPARC-SS reconstructs routes from one specified starting cluster. Dijkstra’s algorithm finds the shortest path from that root to every other reachable destination^17^. Destinations may lie in the initial condition, allowing progression within that condition to be analyzed alongside progression to later conditions. In the primary-to-lung osteosarcoma analysis, P3 was chosen from primary clusters using their median CytoTRACE2 developmental-potential scores^20^ and remained fixed during resampling. An endpoint is a requested graph destination and can be an intermediate biological state.

SPARC-MS treats every cluster at the first sampled time as a candidate source. For each destination at a later time, multi-source Dijkstra identifies the least-cost route and its originating source:

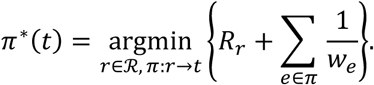

The main analysis used *R_r_* = 0 for all candidate sources. The selected routes identified which candidate sources were assigned paths to later-condition states; some candidates were assigned none. An optional developmental-potential preference assigns a finite starting cost to each candidate source. All initial-condition clusters remain eligible. In resampling round *b*, the median CytoTRACE2 score among retained cells in root *r* was standardized across roots to 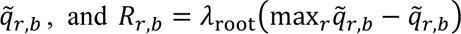. The main SPARC-MS setting was *λ*_root_ = 0; positive values were evaluated to assess how a user-specified source preference changes the routes. Source costs were specified from RNA-derived CytoTRACE2 information before clonal evaluation.

Users can enable the source preference when they want developmental potential to influence the choice among candidate origins. In the tested LARRY datasets, *λ*_root_ = 0.1 produced source costs on a similar scale to RNA route costs and preserved most modal routes, supporting this value as an initial setting for the optional source preference. Its effect depends on how source-cost differences compare with RNA cost differences between competing routes; each application therefore requires its own cost-scale and sensitivity assessment.

### Empirical path uncertainty

Cell subsampling has been used to evaluate trajectory stability and uncertainty in inferred pseudotime^8,11,28^. To assess how cell sampling changes the selected paths, 70%, 80% or 90% of cells were sampled without replacement within each fixed cluster for 1,000 rounds per fraction. The primary analysis used 80%. In each round, the same sampled cells from a cluster were used for all its connections. For every retained cell, the *K* nearest retained cells outside its own cluster were selected in the fixed latent representation. All exclusion-aware connection strengths were recalculated together and the complete edge-weight vector was saved. Each resulting graph was analyzed once with exact Dijkstra using the same starting-population choices, destinations and direction constraints. The analysis used the empirical distribution of *w* or 1/*w*. The resulting frequencies quantify path sensitivity for the fixed representation and clustering.

For each destination, reachability was the proportion of all rounds in which a permitted path existed. Among the reachable rounds, conditional path frequency was the fraction selecting a particular complete cluster sequence. The sequence selected most often was the empirical modal path. Paths were ranked by frequency, using a reproducible lexical rule to resolve ties. The top 20 paths and their cumulative frequency were reported, together with the smallest ranked set covering 95% of the conditional path frequency. This latter set was the empirical 95% path-coverage set. A destination that was never reachable had no conditional path distribution. To assess the effect of associations among edge weights, a control sampled each edge’s empirical marginal independently. Reciprocal representations of the same connection remained coupled, preserving the marginal distributions while removing associations among distinct edges.

Trajectory maps combined selected complete paths and retained their shared segments. Terminal-path usage counted how many displayed destination-specific paths contained each edge. It measures overlap among routes; the edge’s frequency across resampled graphs was reported separately. The displayed union of modal paths could contain paths selected from different sampled graphs, with each path shortest in the graph in which it was selected.

### Transcriptomic cluster reliability

To assess whether expression-derived cluster assignments were reproducible, we evaluated label prediction and marker preservation separately for each LARRY experiment and time point and each osteosarcoma condition. A random forest (RF)^57^ predicted the fixed original labels from the same 30-dimensional latent features used for clustering. Five stratified outer folds provided predictions for held-out cells. Within each outer training set, three-fold sigmoid calibration^58^ was performed without the outer test cells. Forests used 300 trees, a minimum leaf size of two and square-root feature sampling. Cluster-level recall counted the cells with an original label that received the same label in held-out prediction, divided by all cells carrying that original label. Confusion matrices, per-cell scores and calibration curves summarized assignment confidence. Outer folds held out cells within each experiment to assess the reproducibility of expression-derived cluster assignments.

Marker preservation assessed whether subsampling and reclustering retained the expression difference between a cluster and its nearest original competitor. Cells in each original cluster were divided into disjoint marker-discovery and evaluation subsets. Discovery-cell latent centroids identified the nearest competitor, and discovery cells alone defined the bidirectional marker sets and their scoring parameters. For each sampling fraction, 100 condition-specific cell subsamples were used to rebuild the neighbor graph and rerun Leiden with the original parameters. The resulting clusters were matched to the original groups by membership overlap among sampled cells. Frozen marker scores were evaluated only in evaluation cells assigned to the matched cluster and competitor. We reported nearest-competitor marker area under the receiver operating characteristic curve (AUROC), preservation of expression differences and unavailable comparisons. Marker discovery and scoring were held out from one another, while the latent representation and original clustering used the expression data. Recall and marker preservation were reported separately.

We applied the same assessments to the 14 tumor Cas9 units, comprising 121,885 cells and 172 fixed states. RF predictions used frozen 30-dimensional PCA coordinates separately within primary and metastatic compartments. Marker discovery used the 2,000 RNA highly variable genes in the formal expression inputs, with disjoint discovery and evaluation cells and a fixed nearest competitor. For each compartment, 100 unstratified 80% cell subsamples were drawn without replacement. Each subsample was used to rebuild the graph with 30 neighbors and rerun Leiden at resolution 0.6 and seed 159. States with fewer than 50 cells were merged into the nearest larger state in the same compartment, and clusters were matched by sampled-cell membership overlap. Marker AUROC was evaluated in held-out cells from distinct matched groups with at least ten cells in each. Unavailable and merged comparisons were excluded, and marker AUROC was summarized as the median over evaluable rounds (Extended Data Fig. 4d).

### Cell-level pseudotime and gene dynamics

TSCAN provided an earlier use of projecting cells onto cluster-level paths for pseudotime reconstruction^21^. For each selected route, cells belonging to its constituent clusters were projected onto the piecewise-linear curve joining cluster centroids in a covariance-whitened latent representation. Whitening used the regularized covariance matrix^22^, and each cell was assigned to its nearest line segment with the segment coordinate bounded by the two centroids. Cumulative projected arc length was scaled to [0,1] within the route. Pseudotime therefore represents relative transcriptional position along a selected route on a scale from 0 to 1.

GAM-based modeling of expression along pseudotime has been used in TSCAN and tradeSeq^21,24^. To identify expression changes along each route, we fitted Gaussian generalized additive models (GAMs)^23^ using pyGAM^59^, with ten cubic spline basis functions and a smoothing penalty of 0.8. Each route required at least 30 cells. Smooth-term P values were corrected within each trajectory using the Benjamini–Hochberg procedure^25^. Shared-trunk gene peaks were assigned to 20-bin classes. The representative *H19*, *Dlk1* and *H2afj* curves followed the four-route concordance criteria specified in Supplementary Methods. Enrichment used an analysis-specific gene background: the annotated domain for g:Profiler^60^ Gene Ontology (GO), fixed annotated-gene unions from gene-set files for mouse Hallmark/canonical overrepresentation, and the tested-gene background for paired spatial enrichment.

To compare the dynamics of individual genes across the four selected metastatic trunks, we classified the existing GAM results for all 22,489 tested genes. Adjusted GAM P < 0.05 defined significance. Strong dynamics required explained deviance at or above the route-specific 75th percentile, and weak dynamics required values at or below its median. Shared dynamic genes were strong and significant on all four routes with concordant direction. Route-preferential genes were strong and significant on exactly one route, weak on at least two routes and concordant in direction among significant signals. Opposing dynamics required a strong significant increase on at least one route and a strong significant decrease on at least two routes. Direction was determined from the difference in mean expression between the upper and lower pseudotime deciles. Quartiles were used if either decile contained fewer than ten cells.

### LARRY clonal validation and method comparison

To test whether inferred state connections were supported by independent lineage information, we analyzed the LK1, LK2 and LSK LARRY time courses at days 2, 4 and 6^18^. The full inputs contained 28,994, 36,740 and 72,946 cells, respectively. LK1 and LK2 share an initial day-2 pool and subsequently follow distinct culture series. Their results were reported separately, with this shared origin retained in the interpretation of biological replication. Leiden resolution was 0.4 for each day. Barcodes were matched to the expression-derived states for evaluation after inference.

SPARC-MS used the first 30 coordinates of the shared within-day Harmony representation, k_out = 49 outside-cluster neighbors and all day-2 clusters as zero-cost sources. Each of 1,000 80% cluster-stratified cell subsamples generated a complete vector of connection strengths and exact minimum-cost routes. The reported relation graph was the union of empirical modal routes to all later-day destinations.

Slingshot^11^ used the full 50-dimensional Harmony representation and the fixed reporting clusters with getLineages. A prespecified day-2 root oriented its root-to-terminal cluster lineages. Consecutive cluster pairs from these lineages defined the reported relations.

Monocle 3^10^ used the same 50-dimensional representation to construct Uniform Manifold Approximation and Projection (UMAP) with minimum distance 0.3, followed by learn_graph with partitions disabled. The graph root was the principal vertex nearest a representative cell from the prespecified day-2 root cluster. Principal vertices were labeled by the majority reporting cluster among their projected cells. Root-to-leaf principal-graph paths were converted to cluster sequences, collapsing consecutive duplicate labels, and consecutive pairs defined relations.

CStreet^15^ received the shared Harmony coordinates and fixed cluster assignments with additional normalization and log transformation disabled. Its within- and between-day calculations used ten internally reduced components, neighborhood size 15, five sampling trials and seed 159. The reported graph combined its native Otsu-thresholded across-time relations, with maximum outdegree ten, and per-day Otsu-thresholded within-day state relations.

moscot^13^ fitted unbalanced transport between each source day and the immediately following target day in the 50-dimensional Harmony space with regularization 0.005, marginal relaxation 0.95, mean cost scaling, convergence threshold 0.001 and at most 5,000 iterations. Cell-pair transport mass was summed over source and target clusters and normalized within each source cluster and adjacent-day window. The reported relations were across-time connections exceeding the equal-target reference described below.

CellRank 2^14^ constructed a RealTimeKernel from the adjacent-day moscot couplings and within-day connectivity, with connectivity weight 0.2 and 30 neighbors in the shared Harmony representation. Cell transition probabilities into each target cluster were averaged over source-cluster cells. Across-time blocks were normalized within the adjacent target-day window. For within-day blocks, self-transition mass was removed and the remaining entries were renormalized; an unordered relation was retained when either direction exceeded the equal-target reference.

For a source cluster distributing normalized mass uniformly over K target clusters, each target receives 1/K. We retained relations with p_st > 1/K to report connections receiving more support than this equal-target reference. The threshold adapts to the number of eligible target clusters: K is the number at the next day for across-time relations, and K_day−1 after excluding the source itself for within-day relations. This rule was fixed before barcode scoring. The permutation tests below assessed clonal significance separately.

The six methods used all cells from each experiment and the same reporting cluster labels. We evaluated the connections selected by each method, retaining their native directions across time and collapsing duplicate directions into unordered pairs within a day. moscot contributed across-time relations only. Per-method input dimensions, starting-population choices and relation-selection procedures are described above.

For an across-time relation, informative target cells carried clones observed somewhere at the source time; supported target cells carried clones observed in the proposed source cluster. For a within-day pair, we counted supported incidences in both endpoint clusters when the clone occurred in the other endpoint. An endpoint cell was informative only if its clone occurred elsewhere at that day, and each cell contributed at most once per endpoint and relation. Backward across-time relations retained their denominators but received zero support. Significant clonal support was the percentage of all selected relations with enrichment above one and a BH-adjusted stratified-permutation P below 0.05. Cell-weighted consistency pooled supported and informative incidences within each clone fold. Equal-relation consistency averaged the evaluable relation rates, counting every backward relation as zero.

SPARC’s pooled topology contained 59 relations, comprising 54 across-time and five within-day relations; 36 showed significant clone enrichment (61.0%). Cell-weighted consistency pooled supported and informative cell incidences, and equal-relation consistency averaged the evaluable relation rates.

Within-day and across-time relations entered one Benjamini-Hochberg family per method and experiment. Significance used 1,000 day-and-library-stratified clone-label permutations. For pooled cell-weighted and equal-relation summaries, evidence from LK1, LK2 and LSK was combined within each clone fold before averaging the five fold-specific values. Error bars denote sample standard deviation across these evaluation partitions; biological replication follows the experimental design.

Clone folds used seed 20260925, with remaining clones assigned using seed 20260925+7777. Permutation seeds were 20261003 plus the experiment-index offset. RNA-only inference used the full expression input, independently of the clone-evaluation partitions. Supporting data record supported and informative incidences, relation types, selection rules and experiment-specific results. Shared clones support common ancestry and clonal consistency across the connected states.

### Tumor Cas9 lineage-tracing benchmark

To assess agreement with tumor lineage evidence, we analyzed 14 complete primary-metastasis units from 11 independent tumor families in KP-Tracer^2^, macsGESTALT^3^ and the Quinn xenograft datasets^1^. Each unit used all cells. Counts were normalized to 10,000 per cell and log1p-transformed, and 2,000 highly variable genes and 30 principal components defined the expression representation. Primary and metastatic cells were clustered separately using 30 neighbors, Leiden resolution 0.6 and seed 159. States with fewer than 50 cells were assigned to the nearest larger state in the same compartment. SPARC used k_out=49 and the public sparc.run v0.3.0 implementation on the complete permitted graph.

Cas9 lineage information was used after RNA-derived scoring to quantify agreement for each state pair. In each primary-metastasis unit, the conditional lineage-transition matrix provided the forward entry for a primary-to-metastasis pair and the mean of the two directional entries for a within-primary or within-metastasis pair. Spearman correlations compared these lineage scores with each method’s expression-derived scores over the combined within-primary, primary-to-metastasis and within-metastasis pairs. SPARC used its cluster-excluded global kNN connection strengths. CellRank 2^14^ combined separately estimated within-compartment kNN transitions with an adjacent-compartment moscot coupling. The within-primary component received weight α, the primary-to-metastasis component received 1 - α and the within-metastasis component retained unit weight. We evaluated 19 values from 0.05 to 0.95, using k=50 for the main fixed and calibrated comparisons; the sensitivity curves also show k=30 as a neighborhood control.

Topology sensitivity was measured from the N_states −1 highest-scoring state pairs at each α by their fractional turnover from the α=0.50 set. This operation selected a fixed-size diagnostic edge set. For family-held-out calibration, α was selected using the highest mean all-block Spearman correlation among the other ten families, with ties resolved first by proximity to 0.50 and then by the smaller value, and applied to the held-out family. Quinn cell-number units were grouped as one family. Curves and intervals used equal family weighting and 10,000 family-cluster bootstrap samples with seed 159.

### Simulation-based topology and sample-size assessment

To measure recovery of known directed and branching structures, we generated time-ordered Splatter^29^ datasets with four topologies, three difficulty settings and seeds 159-161, giving 36 complete datasets. Directed-edge F1 and directed Hamming–Ipsen–Mikhailov (HIM) distance^8,30^ were calculated on the common supplied states. Cell-number robustness was assessed at seven sizes from 200 to 12,800 using five paired nested replicates. The time-ordered simulations used a supplied initial root.

To determine whether reconstruction depended on one expression-generating mechanism, we implemented two complex reference state lineages, each with four direct primary-to-lung routes, in four simulators. PROSSTT generated branchwise expression programs along pseudotime^31^. Splatter’s Splat differentiation-path model generated count distributions and differential expression along paths^29^. TedSim generated expression states through cell division and differentiation on a state tree^33^, and dyngen used stochastic gene-regulatory reactions^32^. Keeping the reference topologies fixed allowed route and topology recovery to be compared across these different assumptions. After native generation, a common observation layer applied the settings for measurement noise, expression-dependent dropout and batch shifts.

The generative mixed benchmark comprised 256 datasets from four simulators, two complex topologies, two transition-mass modes, two expression regimes and eight seeds. Each dataset contained 8,000 cells and 2,000 genes, with 4,000 cells at each site. Branched metastatic progression contained 15 states and the Deep asymmetric cascade 18 states; both contained four direct primary-to-lung routes. Homogeneous (homogeneous_020) allocated 1,000 Lung cells per complete subtree. Heterogeneous (metastasis_state_specific) used masses 0.85, 0.55, 0.25 and 0.05 to allocate 2,000, 1,294, 588 and 118 descendants. The deterministic largest-remainder rule assigned integer totals, and cells within each subtree retained the specified relative state allocation. For each descendant allocation, state populations were sampled and RNA observations generated before PCA and trajectory inference. Seeds 159, 161 and 162 supplied 96 calibration datasets; seeds 164–168 supplied 160 test datasets. SPARC fixed P0, k_out = 49 and 1,000 complete-vector rounds at 80% within-cluster cell retention. Main CellRank 2 used k = 50 and 19 alpha values. Condition-specific calibration minimized primary-state within-mass mean absolute error on the three calibration seeds and applied that selected alpha unchanged to the five test seeds: 0.20 in Homogeneous, 0.85 for Branched Heterogeneous and 0.55 for Deep Heterogeneous.

To distinguish accurate transition estimates from accurate reconstruction of full trajectories, we evaluated ranking average precision, fixed-budget pair F1/HIM, root-to-terminal recovery and all-state AD/CASet separately. For root-to-terminal comparisons, CellRank 2 routes minimized the sum of negative log transition probabilities from the supplied root to each configured lung terminal. For AD/CASet comparisons, its matrix was decoded into a fixed-root maximum-probability arborescence^61^ by maximizing the sum of log transition probabilities. These two comparisons used the respective path and all-state decoding objectives. Simulation intervals used five held-out seed clusters. The separate real Cas9 evaluation used family-cluster uncertainty, as described above. Changes in routes to fixed ground-truth lung destinations relative to alpha = 0.50 were reconstructed from the saved alpha-specific trees.

All-state comparisons adapted ancestor–descendant (AD) distance^34^ and common-ancestor-set (CASet) distance^35^ to expression-state trees by assigning one unique cell-state label to each vertex. Truth and prediction used the same complete state set and fixed root. Union-normalized AD was the size of the symmetric difference between strict ancestor-pair sets divided by their union size, with zero assigned to an empty union. Self-pairs were excluded. CASet averaged Jaccard distances between common-ancestor sets over all unordered state pairs; each state’s ancestor set included the state itself.

### Murine orthotopic osteosarcoma model and scRNA-seq sampling

Three independently established murine osteosarcoma cell lines (M1332, M408 and M1199) were used to generate syngeneic orthotopic tumors in immunocompetent C57BL/6 mice. The model system and establishment of tumor-derived cell lines were described previously by Nirala et al.^19^ Each mouse received an intratibial injection of 1 × 10^6^ cells. At the study-defined euthanasia endpoint, the primary tibial tumor and visually apparent spontaneous pulmonary metastatic lesions were collected and processed separately for scRNA-seq. Primary-tumor and lung-lesion profiles were retained as distinct anatomical conditions for subsequent analysis.

To estimate the cell-state organization shared across the three mice, quality-controlled cells were combined within the primary-tumor and lung conditions before trajectory reconstruction. Cells from multiple specimens have also been combined to reconstruct developmental manifolds and trajectories in mouse and zebrafish studies^10,62,63^. Here, the combined-cell trajectory describes shared cell-state organization, while the individual mouse remains the biological replicate.

### Murine primary-to-lung SPARC analysis

After quality control and malignant-cell selection, primary and lung cells were represented in shared Harmony-corrected PCA coordinates. The graph allowed reciprocal within-site and forward primary-to-lung connections. SPARC-SS fixed P3, k_out = 49 and the complete condition-permitted cluster-pair universe. One thousand complete-vector subsampling rounds were evaluated at each of 70%, 80% and 90%, with 80% used for the principal analysis. Every non-root cluster was eligible as a graph destination. The complete map contained 31 states and 30 union edges from paths to 17 lung and 13 non-root primary destinations. Primary-local programs used the five primary branch tips P0, P13, P5, P1 and P12; P7 was an intermediate on the route to P5. The P12 route contained 3,060 cells and 22,489 tested genes. The primary core contained 1,185 increasing and 786 decreasing genes; its intersections with the metastatic core contained 209 increasing and 480 decreasing genes. Metastatic-associated sets contained 469 increasing and 1,194 decreasing genes. Metastatic programs used four selected trunks: MR1, P3 → P10 → L13 → L10; MR2, P3 → P9 → P8 → L2; MR3, P3 → P9 → L4; and MR4, P3 → P2 → L9. Later lung extensions were analyzed separately. Gene-peak GAMs included root cells; the four-bin gene set variation analysis (GSVA)^64^ temporal analyses excluded P3 and assessed increasing, decreasing, transient-peak and transient-valley pathway profiles.

MR1–MR4 span inferred cross-site transitions from P10, P8, P9 and P2, respectively. Their complete node sequences were selected in 100%, 100%, 99.2% and 100% of the 1,000 rounds at 80% cell retention. All rounds were reachable for these destinations. This focused comparison covers four primary transition states. Extended gene-program analyses of the routes to L6 and L7 were performed separately from the four-trunk comparison.

### Human osteosarcoma specimens and Visium HD data generation

Three human osteosarcoma sections were used in the spatial analyses: a primary bone tumor from one patient (Bone; original specimen ID JY4), a pulmonary metastasis from a second patient (Lung; original specimen ID JY10), and the paired primary bone tumor from that second patient (original specimen ID JY8). Spatial transcriptomes were generated using the 10x Genomics Visium HD platform. Bone and Lung supported the full segmented-cell analysis and were from different patients. Only a small tumor-containing region of the paired primary specimen passed the criteria required for the mature-resorptive footprint analysis; this specimen was therefore limited to assessment of spatial presence in the paired specimens. JY23 was excluded because of insufficient data quality. Sequencing reads were aligned and mapped with 10x Genomics Space Ranger v4.0, using the platform’s native nucleus-guided segmentation to assign 2-µm HD bins to segmented objects.

### Cell segmentation and construction of the single-cell count matrix

Analyses were based on the Space Ranger segmented-cell output, in which built-in nuclear segmentation assigns the 2-µm capture bins to individual cells; each segmented cell constituted one observation. A spatial coordinate for each cell was derived as the centroid (mean pixel position) of its constituent 2-µm bins, using the sample-specific tissue-position and barcode-mapping files provided by Space Ranger. Pixel coordinates were converted to microns using the sample-specific scale factor (0.2297 µm px⁻¹ for both specimens); all reported distances are given in microns.

### Spatially aware quality control

Raw segmented cells exhibited low transcript capture (median 47–51 detected genes per cell). A modified neighborhood-based quality-control procedure, inspired by SpotSweeper^65^, identified local outliers relative to each cell’s tissue neighborhood. For each cell, the 36 nearest spatial neighbors (excluding the focal cell) were used to calculate the mean and sample standard deviation of each QC metric: log₁p-transformed total counts, log₁p-transformed detected-gene counts, and mitochondrial-count fraction. The focal value minus its neighbor mean was divided by the neighbor standard deviation to obtain a local z-score. A standard deviation below 10⁻¹⁰ gave a score of zero. A cell was flagged if its log-total-count or log-gene-count z-score exceeded |z| > 3 in either direction, or if its mitochondrial fraction z-score exceeded z > 3 (upper tail only). Cells with fewer than 5 detected genes were additionally flagged by a hard minimum threshold. Flagged cells were removed (logical OR of the two criteria), and genes detected in fewer than 10 cells were subsequently dropped. The procedure evaluates each cell relative to its spatial neighbors, accounting for local variation in transcript capture. After quality control, the annotated datasets retained 255,112 (Bone) and 249,756 (Lung) cells. For sensitivity analyses, a more stringent global-filter subset (≥20 detected genes; total counts within the 1st–99th sample percentile; genes in ≥10 cells) retaining approximately 130,000 cells per specimen (median 90–95 detected genes per cell) was also defined.

### Spatially informed clustering

Quality-controlled cells were clustered with BANKSY^66^ (Bioconductor package Banksy), which augments each cell’s own expression profile with a spatial neighborhood-expression term, enabling clusters that reflect both transcriptional identity and tissue context. Counts were library-size normalized over all quality-controlled genes without highly variable gene subsetting, and BANKSY was run with spatial mixing parameters λ = 0.2 and λ = 0.8, a geometric neighborhood of k geom = 15, 20 principal components, and Leiden clustering at resolution 0.6 (random seed 1000). The λ = 0.8 solution, which places greater weight on spatial context, was carried forward as the basis for cell-type annotation.

### Consensus cell-type annotation and expert review

BANKSY clusters were annotated by a four-method consensus approach. (i) Reference correlation: each cluster’s pseudobulk expression profile was compared with cell-type profiles from a mouse osteosarcoma single-cell RNA-seq reference (see below) by Spearman correlation over 3,000 shared highly variable genes. (ii) Canonical marker scoring: curated, site-aware osteosarcoma and tumor-microenvironment marker panels were scored as the product of mean expression and detection fraction per cluster. (iii) Differential expression: one-versus-rest Wilcoxon rank-sum tests (with tie correction; top 50 markers per cluster) were cross-referenced to the marker panels using scanpy.tl.rank_genes_groups.^67^ (iv) Spatial deconvolution: Tangram^68^ (tangram-sc) was applied in cluster mode with an RNA-count-based density prior, projecting reference cell-type labels onto BANKSY clusters. A consensus label was assigned to each cluster by majority vote across the four methods, with Tangram used to break ties. A domain expert subsequently reviewed all consensus labels, manually overriding a small number of clusters where warranted; these curated labels were used for all subsequent analyses. The nuclear segmentation yielded a combined macrophage/monocyte compartment, labeled Macrophage. The Lung analysis retained 249,756 nucleus-guided segmented objects. BANKSY clustering assigned 15,537 objects to an osteoclast-associated cluster, designated the osteoclast-rich region. Individual-object marker scoring and grouping of neighboring nuclear segments yielded an estimated 1,300–1,800 osteoclasts. For section-level normalization, this estimate corresponded to approximately 0.5– 0.75% of the total segmented-object count. Osteoclast-associated expression was concentrated within tumor nodules. Spatial neighborhood and ligand–receptor analyses used the cluster-level annotations.

### Reference dataset and cross-species gene-symbol conversion

The annotation reference was a mouse osteosarcoma single-cell RNA-seq atlas comprising 76,084 cells across 16 cell types, with gene symbols mapped to human orthologs. Because all SPARC-derived signatures are defined in mouse gene space, mouse symbols were converted to human orthologs prior to scoring: a curated map resolved non-trivial conversions (e.g. Cd24a → CD24; H2 MHC class-I/II genes → HLA-DRA, HLA-DRB1, HLA-A, HLA-B), after which remaining symbols were upper-cased and matched case-insensitively (ignoring hyphens and underscores) to the Visium HD feature space. Unresolved genes were excluded; node signatures with no recovered genes were assigned missing values.

### Module and signature scoring

Unless specified otherwise, module and signature scores were calculated with scanpy.tl.score_genes^67,69^ using use_raw = False, 25 control bins and random state = 0. Raw counts were normalized to 10⁴ per cell and log₁p-transformed. The control-set size was min(50, n genes) for the metastatic-route, metastasis-versus-local progression and OLC-stage scores where specified, and the Scanpy default of 50 otherwise. Per-cell scores were exported in files matched by cell barcode, with the source AnnData objects preserved.

### Spatial smoothing of scores and expression

To display spatial patterns, we averaged each cell’s score or log-normalized expression over its 36 nearest spatial neighbors, including the focal cell. Each feature used a shared color scale based on the pooled 1st–99th percentiles from the two specimens. Quantitative statistics used the unsmoothed per-cell values.

### Spatial recovery of SPARC tumor-trajectory programs

To examine where programs associated with mouse trajectory states were expressed in human tissue, we scored osteosarcoma-lineage cells for all 31 mouse cluster signatures. Each signature contained the top-20 one-versus-rest markers from the mouse SPARC object. A marker was counted as detected when its human ortholog occurred in the shared Bone and Lung feature space. We summarized signature recovery on the mouse trajectory topology rooted at P3. Site consistency required a higher mean score in Bone for primary P-nodes and in Lung for metastatic L-nodes. Representative signatures were shown as k = 36-smoothed spatial maps.

### Full-topology spatial program controls

Full-terminal increasing/decreasing modules were calculated with Pearson association on nine complete lung-terminal routes, including states beyond the shared-trunk anchors. Genes required route-specific adjusted P < 0.01 and concordant direction in at least five of nine routes. The top twenty human-matched genes by mean absolute Pearson correlation formed each module. These modules served as supplementary spatial controls. The primary-local versus metastatic gene classes used the five primary-local routes and four anchor-terminated trunks defined above.

### Tumor zonation and zone-resolved analyses

The Lung specimen was partitioned into tumor zones based on local tumor-cell density. For each cell, tumor density was computed as the fraction of tumor cells (OS differentiated, Tumor uncertain) among its 200 nearest neighbors, and cells meeting a density threshold of ≥0.5 were classified as tumor-positive. The resulting mask was rasterized on a 20-µm grid and morphologically processed to yield a spatially coherent tumor region: a tissue mask was formed by binary closing (40-µm kernel), hole-filling, and opening; the tumor region was generated by rasterizing, intersecting with the tissue mask, closing (100-µm kernel), hole-filling, dilation (60-µm kernel), and re-intersection with the tissue mask, retaining connected components of at least max(500, 5% of the largest component) bins. Tumor boundaries were identified from label transitions across each cell’s 20 nearest neighbors (with despeckle filtering), and a signed distance to the boundary was computed for every cell. Cells were assigned to three zones: core (>100 µm interior to the boundary; 129,893 cells), boundary (within ±100 µm; 41,873 cells), and exterior (>100 µm exterior; 77,990 cells). Within each zone, ligand–receptor proximity was recomputed at 50 µm as described above. Pairwise cell-type co-localization was quantified as the O/E ratio of the mean neighbor-type fraction over each cell’s 36 nearest neighbors (excluding the focal cell) relative to zone-wide cell-type abundance. Boundary-versus-exterior macrophage differential expression was assessed by per-cell Wilcoxon rank-sum tests on log₁p-normalized counts, requiring at least 50 cells per group and reporting genes with adjusted P < 0.05 and |log₂ fold change| > 0.5.

### Metastatic-route programs and route–microenvironment neighborhoods

To compare the tissue locations of programs associated with alternative metastatic routes, we defined signatures from the mouse MR1-MR4 pseudobulk expression matrix. Genes with positive expression in a route were ranked by that expression minus the mean of the other three routes, and the top 100 genes formed its signature. After mapping to human orthologs, the mouse-defined signatures were scored in independently collected patient Visium HD profiles. Each OS-lineage cell received all four scores and was assigned to the program with the highest score. Neighborhood enrichment compared the mean cell-type composition among each cell’s 36 nearest neighbors with the global tissue composition, expressed as an observed-to-expected ratio.

Within Lung, spatial neighbors within 25 and 50 µm were used for each OS-lineage cell to calculate the local fraction of objects assigned to the osteoclast-rich region, the local SPP1-positive fraction (raw SPP1 count > 0), and the mean mature-resorptive-OLC stage score. Continuous associations between route score and each neighborhood metric were quantified by Spearman correlation. For each route, neighborhood enrichment compared the fraction of route-dominant cells with at least one neighboring object assigned to the osteoclast-rich region, or at least one high-mature-OLC neighbor defined by the top score decile, against the corresponding fraction among OS-lineage cells assigned to the other routes. Observed-to-expected ratios and one-sided Fisher’s exact tests were reported with Benjamini– Hochberg correction. Sensitivity analysis removed objects assigned to the osteoclast-rich region from the sender population,and tumor-zone labels were inherited from the zonation analysis.

### External conventional Visium route–niche validation

All 26 conventional Visium sections from 19 patients in GSE293065 (9 primary bone tumors, 14 distant metastases and 3 local relapses) were scored for the four SPARC metastatic-route programs and niche signatures.^42^ Osteoclast-marker genes specified by the analysis configuration were removed from route signatures, and route-signature genes were excluded from the macrophage panel. Within each section, niche-high spots were defined by the upper quartile of the corresponding signature score. Route enrichment was the mean route score in those spots minus its mean in the remaining spots. Osteoclast sensitivity analyses used a canonical twelve-gene panel, a terminal four-gene panel (CTSK, ACP5, MMP9 and DCSTAMP), and route scores residualized by ordinary least squares against macrophage score and an intercept within each section. MR1-minus-MR3 enrichment contrasts were tested across sections with two-sided paired Wilcoxon signed-rank tests; the displayed P values are unadjusted. Myeloid, endothelial and T-cell comparisons used the same enrichment definition, with mean and standard error of the mean (SEM) summarized across sections.

### Mouse myeloid-to-OLC trajectory and cross-species spatial scoring

To reconstruct the relationship between mouse lung monocyte, macrophage and OLC states, we analyzed 7,681 cells in 12 fixed clusters using the 30-dimensional within-lung Harmony representation. Mono1, with the highest median CytoTRACE2 potency score (0.285), was the fixed source for SPARC-SS. Each cell selected its 49 nearest neighbors outside its own cluster in Harmony space. Connection strengths were calculated for all permitted within-lung pairs, with positive same-condition edges available in both directions. Dijkstra then used inverse-connectivity costs on the complete permitted graph. The selected route was Mono1 → Mφ3 → OLC. Program analysis excluded one cell originally annotated as Osteoclast that reclustering assigned to Mono1. The retained route contained 1,975 cells: 1,053 Mono1, 801 Mφ3 and 121 OLC.

Full annotation summaries contained 114 primary-site and 122 lung-site osteoclast-like cells in the mouse data. Our reanalysis of the published human cohort^41^ counted 8,806 osteoclast-annotated cells in primary bone tumors, 483 in lung metastases and 190 in recurrent samples.

Empirical path stability was evaluated at 70%, 80% and 90% cluster-wise cell-sampling fractions. For each fraction, 1,000 samples were drawn without replacement from every cluster; the outside-cluster neighbor graph and complete vector of connection strengths were recalculated, and exact Dijkstra inference was repeated with fixed clusters, source, endpoint and permitted edges. Mono1→Mφ3→OLC was reachable and selected in every round at all three fractions. Cells in the three route clusters were projected onto adjacent route segments after whitening the Harmony coordinates by the full-data covariance matrix. Arc length along these segments defined pseudotime. Running means summarized osteoclast-resorption, fusion-commitment and AP-1 immediate-early module scores, together with Nfatc1 expression, along the route.

RANKL communication was evaluated separately in primary and lung profiles with CellChatDB.mouse and cell-type grouping.^40^ Cell types with fewer than ten cells were excluded. Communication probabilities used truncated means with trim = 0.05, raw.use = TRUE and population.size = FALSE; filterCommunication used min.cells = 10. Pathway aggregation and network centrality supplied sender, receiver, mediator and influencer roles. For the paired role display, each role was divided by its maximum across both regions, giving primary and lung the same visualization scale.

The four-state spatial program map used a broader mouse myeloid reference partition: 3,485 early monocyte/inflammatory cells, 635 macrophage-intermediate cells, 801 osteoclastogenic pre-OLC cells and 121 mature-resorptive cells. One-versus-rest Wilcoxon tests within this 5,042-cell reference selected the top 30 positive markers per group with detection in more than 10% of group cells. Ortholog matching recovered 13, 22, 19 and 23 human genes, respectively. These signatures were scored in all Bone and Lung cells after normalization to 10,000 counts and log transformation, using Scanpy score_genes with use_raw = FALSE, control-set size min(50, signature size) and seed 0. The Lung map displayed the signature with the highest score after averaging each score over the 36 nearest spatial neighbors (Fig. 6b).

### Transcription-factor activity estimation and cross-species comparison

To compare changes in transcription-factor activity between mouse and human lung myeloid states, we used signed CollecTRI regulons^70^ retrieved through OmniPath^71^ and the decoupleR univariate linear model (ULM)^72^. The full annotation pools comprised 7,681 mouse cells (6,010 monocyte, 1,549 macrophage and 122 osteoclast-labeled cells) and 1,763 human lung-metastasis cells from GSE152048 (272 monocyte, 1,052 macrophage and 439 osteoclast-labeled cells). The log-normalized input matrices contained 22,489 mouse and 13,764 human genes.

Human TF–target pairs were deduplicated by source and target symbol. Exclusively inhibitory annotations received weight −1 and other retained interactions received +1. For mouse scoring, TF and target symbols in this network were converted to mouse-style capitalization and intersected with the measured genes. TFs with at least five represented target genes were scored using run_ulm; each cell’s activity score was the regression slope t-statistic. Scores were averaged within each species and annotation group, and a regulon shift was defined as mean osteoclast-labeled activity minus mean monocyte-labeled activity.

Cross-species comparison matched TF symbols after conversion to uppercase and included every match with finite contrasts in both species. Of 662 matched TF activity contrasts, 417 had the same sign and 245 had opposite signs.

### Spatial ligand–receptor proximity (SpaLRP)

Spatial co-localization of cognate ligand–receptor pairs was assessed with a nearest-neighbor proximity approach. A segmented object was classified as ligand- or receptor-positive if its raw count for the corresponding gene exceeded zero; multi-subunit complexes required all components to be detected. Annotation groups with fewer than 20 positive objects for a given gene were excluded. For each ligand-sender × receptor-receiver × annotation-group combination, the distance from each ligand-positive sender object to its nearest receptor-positive receiver object was computed, and the observed count was defined as the number of sender objects whose nearest receiver fell within each of five distance thresholds (15, 25, 50, 100, and 250 µm). A spatial-density reference count was approximated assuming uniform receiver density over the tissue bounding-box area: E = max(n sender · min(1, π·r²·n receiver / A), 0.5). Enrichment was tested with a one-sided Fisher’s exact test (alternative “greater”), comparing the in-range fraction of the focal sender annotation group against all other ligand-positive annotation groups,with Benjamini–Hochberg correction applied across all pairs within each sample × distance threshold. Results were considered significant at adjusted P < 0.05, with a minimum of 5 observed in-range pairs required.

### SPARC analysis of paired patient spatial transcriptomes

Five paired conventional Visium comparisons from GSE293065^42^ were analyzed independently: four primary bone tumor to lung metastasis pairs and one primary tumor to local relapse pair. Raw filtered matrices, Space Ranger tissue coordinates, image scale factors and H&E images were used. In-tissue spots with at least 200 detected genes were retained, counts were normalized to 10,000 and log-transformed, and up to 3,000 highly variable genes were selected within each pair. BANKSY^66^ incorporated expression from the six nearest within-section spatial neighbors with neighborhood contribution λ = 0.2. The resulting augmented representation was reduced to 50 principal components and supplied to SPARC. Spatial neighborhoods were restricted to individual tissue sections. Section identity was confounded with progression state, so Harmony was omitted.

For each pair, SPARC-SS used 30 components, K=29 cluster-excluded outside-cluster neighbors, Leiden resolution 0.8 and the complete condition-permitted candidate universe and random seed 159. Clustering was performed separately within the primary and progression sections. CytoTRACE2 was applied to raw counts from the primary section, and the retained primary graph state with the highest median spot-level score initialized the root. The fixed roots for Pairs 1 to 5 were Pri3, Pri2, Pri2, Pri2 and Pri6. The graph supplied to Dijkstra contains all permitted within-condition edges. One thousand complete-vector cell-subsampling rounds were run at each of 70%, 80% and 90%, with 80% used for the principal modal paths. Normalized pseudotime was mapped to H&E coordinates. Local arrows represent the fitted direction of increasing pseudotime among neighboring spots. Dynamic genes required a Benjamini-Hochberg-adjusted GAM P < 0.05 and explained deviance > 0.10. Pathway enrichment was performed separately for increasing and decreasing genes, and recurrence of predefined programs was summarized across independently reconstructed patient-specific terminal routes.

### Cross-resolution lung osteoclast analyses

Lung-local myeloid-to-osteoclast progression was analyzed independently in the Pair 4 conventional Visium lung section and in segmented single-cell-scale Visium HD objects from Lung. The conventional Visium SPARC-SS analysis used K=19 cluster-excluded outside-cluster neighbors and fixed Macrophage0 as the root after nomination by the marker contrast. It used complete marker-supported BANKSY clusters and an ordered macrophage-like, osteoclast-precursor and mature-osteoclast design. The earliest stable state resolved in the conventional data was macrophage-like, so Macrophage0 served as the fixed root without CytoTRACE2 scoring. In Visium HD, SPARC-SS used K=29 cluster-excluded outside-cluster neighbors and fixed LuM-C3 as the root after nomination by the monocyte-like program. It was applied to 6,483 lung-section objects using the BANKSY representation, 30-neighbor Leiden clustering at resolution 0.55 and random seed 159. Terminal graph leaves were discovered before post hoc osteoclast annotation. Terminal-state stability was evaluated across four Leiden resolutions and three random seeds. Gene-level agreement between platforms was evaluated on shared canonical genes using late-minus-early trajectory effects and Spearman correlation. Both applications used the complete condition-permitted candidate universe and 1,000 complete-vector cell-subsampling rounds at 70%, 80% and 90%.

### Statistics and reproducibility

Full segmented-cell spatial analyses used Bone and Lung as independent-specimen corroboration. Assessment of the paired patient specimens was confined to the presence of a mature-resorptive footprint in the limited paired primary region and the corresponding Lung specimen; abundance comparisons used Bone and Lung separately. The external conventional Visium validation included 26 sections from 19 patients, whereas direct paired-patient spatial SPARC was performed independently for five primary-to-progression pairs. Effect sizes, including observed-to-expected ratios and Spearman correlations, were treated as the primary readouts. Because cell-level sample sizes were large, P values were interpreted alongside effect sizes and biological replication. One-sided Fisher’s exact tests were corrected within their respective test families using the Benjamini–Hochberg procedure. Per-cell differential-expression analyses summarized within-specimen variation, with biological replication defined at the specimen or patient level.

Software versions and random seeds were retained separately for each analysis environment. LARRY clustering used seed 0; osteosarcoma and paired spatial inference used seed 159. RF sampling and calibration, graph subsampling and clone evaluation used the dedicated seeds described in Supplementary Methods.

## Data Availability

The paired mouse osteosarcoma scRNA-seq data generated in this study have been deposited in the Gene Expression Omnibus (GEO) under accession GSE343530, and the patient-derived osteosarcoma Visium HD data under GSE343536. Published human osteosarcoma scRNA-seq data are available under GSE152048^41^, and the external conventional Visium cohort under GSE293065.^42^

LARRY expression and clonal assignments are available under GSE140802, sample GSM4185642 (stateFate_inVitro).^18^

Processed KP-Tracer expression and lineage-tracing data are available in Zenodo (doi:10.5281/zenodo.5847462).2 For macsGESTALT, expression data are available under GSE173958, and clone assignments and reconstructed phylogenies in Mendeley Data (doi:10.17632/t98pjcd7t6.1).3 Quinn A549 xenograft expression and lineage-tracing data are available under GSE161363.^1^

Longitudinal HGSOC data are available under GSE165897.^51^

Synthetic count matrices can be regenerated using the PROSSTT, dyngen, Splatter and TedSim generation scripts, reference topologies, parameter files and random-seed manifests supplied with the study’s reproducibility materials.^29,31–33^ These materials, empirical path-frequency estimates, benchmark evaluation results and numerical figure source data are available from the authors during peer review and will be released on GitHub upon acceptance of the manuscript.

Supporting gene-level trajectory results, route definitions and numerical outputs for MR1–MR4 and the routes to L6 and L7 are available from the authors during peer review and will be released on GitHub upon acceptance.

## Code Availability

The SPARC package and analysis code are available to editors and reviewers during peer review and will be released on GitHub upon acceptance of the manuscript.

## Acknowledgments

We thank the non-author members of the Hassan laboratory for their technical assistance, helpful discussions and support throughout the study.

## Author contributions

S.W. conceived the study, developed the computational framework and analysis code, and performed all computational analyses other than the Visium HD spatial data analysis. S.W. interpreted the results, wrote the original draft of the manuscript, and prepared the figures and figure legends. W.C.W. performed the Visium HD spatial data analysis, generated the corresponding figures, and contributed to the interpretation of the spatial results. B.N. designed and performed the biological experiments and generated the experimental samples and data. C.A.M. contributed to the development of the analytical strategy and interpretation of the results. J.T.Y. supervised the experimental studies, provided resources, and contributed biological guidance and interpretation. M.A.H.S. supervised the computational work and the overall project, provided conceptual guidance and mentorship, acquired funding, and critically revised the manuscript. All authors reviewed and approved the final manuscript.

---

**Extended Data Fig. 1.**
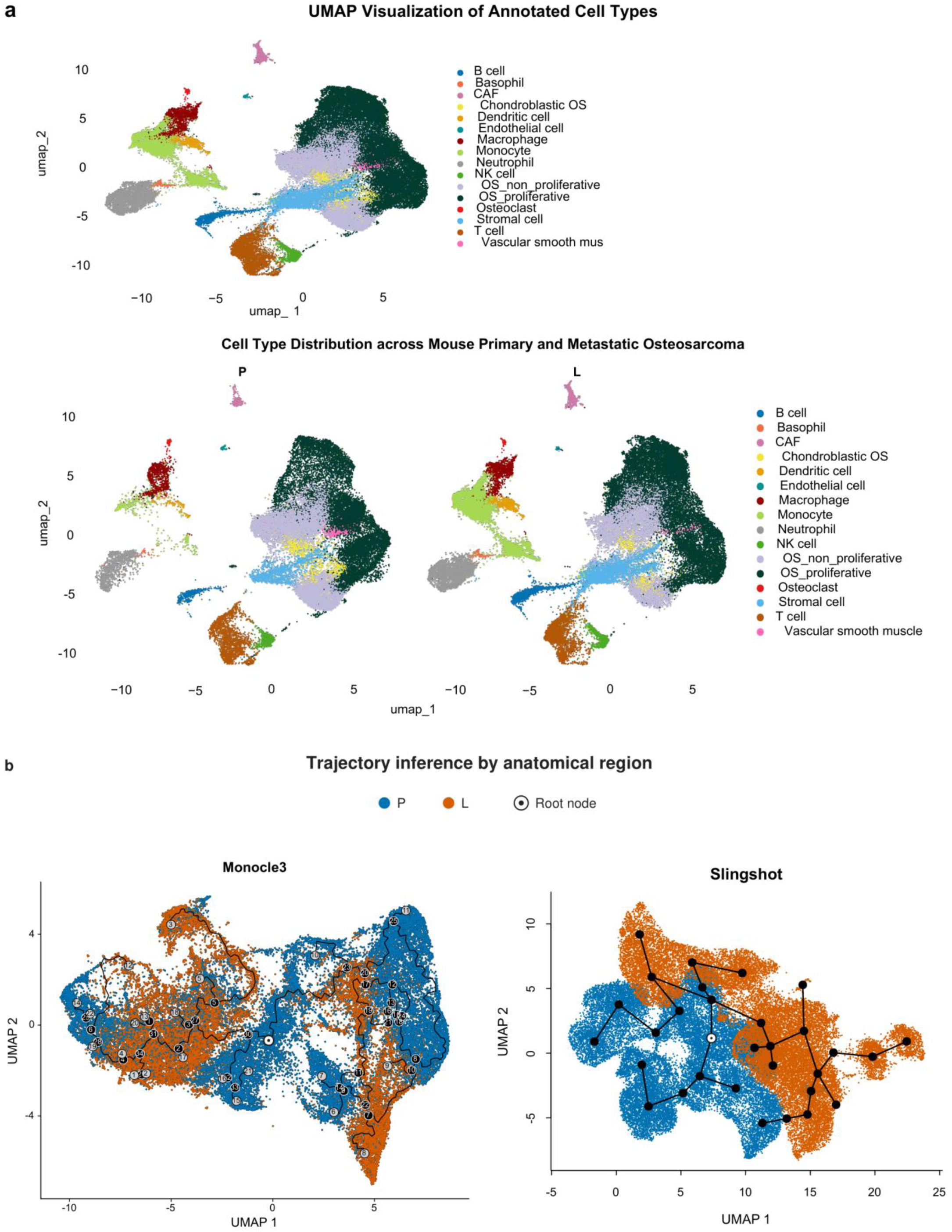
Mouse osteosarcoma cell-state context and alternative trajectories. (a) Mouse cell-state UMAP colored by reviewed cell type, shown jointly and separately for primary (P) and lung (L) origin. (b) Monocle 3 and Slingshot trajectory structures inferred from paired malignant-cell expression profiles. Blue and orange points denote primary (P) and lung (L) cells, respectively. Lines show the native Monocle 3 principal graph and Slingshot cluster tree. P3, the primary cluster with the highest median CytoTRACE2 score, was used to nominate the roots; ring-and-dot symbols mark the corresponding Monocle 3 principal-graph vertex and Slingshot starting cluster. The learned topologies contain primary–lung–primary paths. In the fixed Slingshot tree, such paths persist from every primary starting cluster.

**Extended Data Fig. 2.**
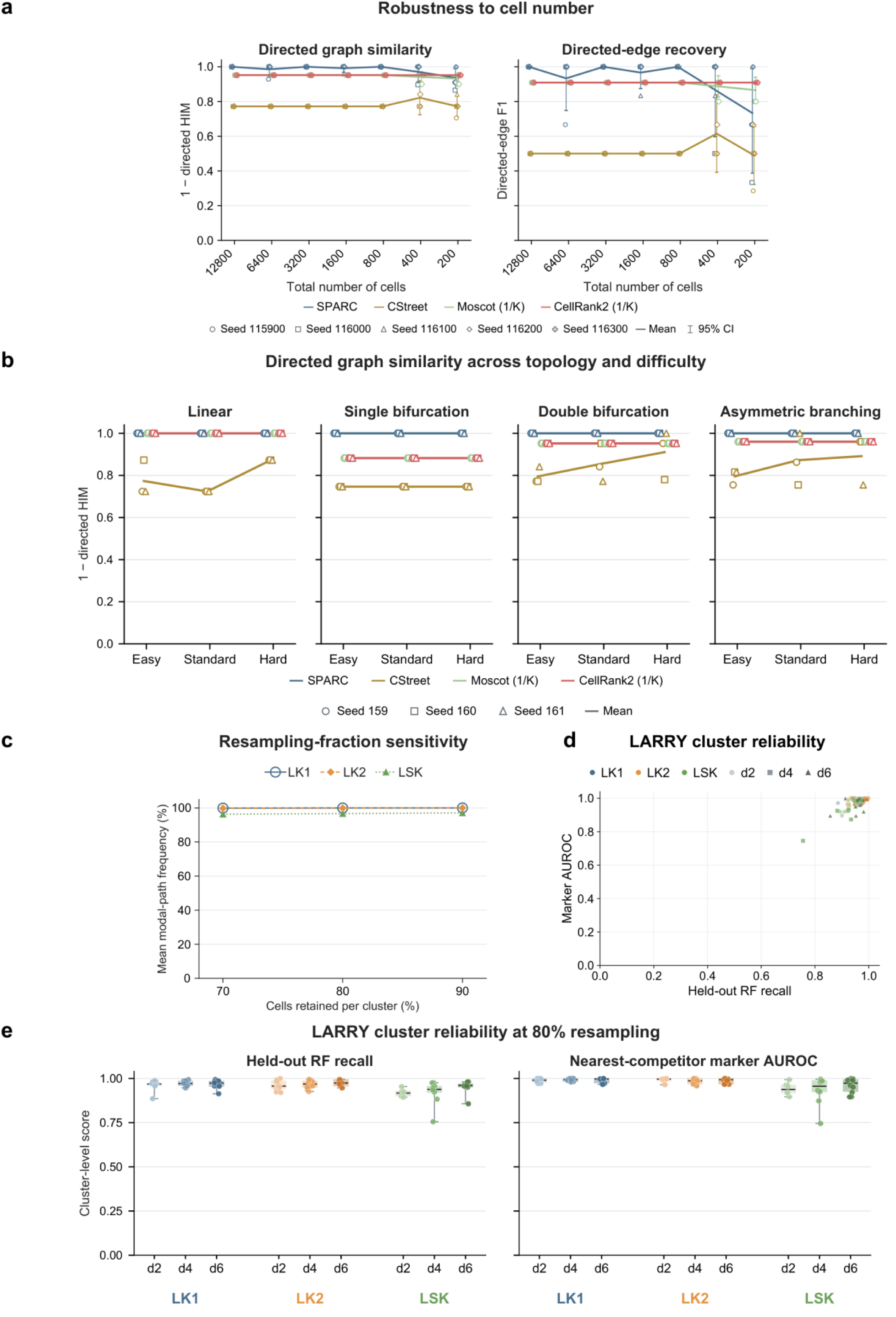
Cell-number robustness and transcriptomic reliability. (a) Directed Hamming–Ipsen–Mikhailov (HIM) similarity and directed-edge F1 across seven nested cell-number settings with five paired sampling replicates. Lines summarize each method and symbols show individual replicates. (b) Directed HIM similarity across four topologies, three difficulty settings and three generation seeds. (c) Mean empirical modal-path frequency under 70%, 80% and 90% cell retention within each fixed cluster for LK1, LK2 and LSK. LK1 and LK2 denote culture-series identifiers; LSK denotes lineage-negative, Sca1-positive, Kit-positive progenitors. Each fraction used 1,000 complete-vector graph-reconstruction rounds. (d) Held-out random forest (RF) recall versus nearest-competitor marker area under the receiver operating characteristic curve (AUROC); each point is a cluster, with experiment encoded by color and day by symbol. (e) Cluster-level RF recall and nearest-competitor marker AUROC by experiment and day at 80% resampling. Boxes and individual points summarize the cluster distributions.

**Extended Data Fig. 3.**
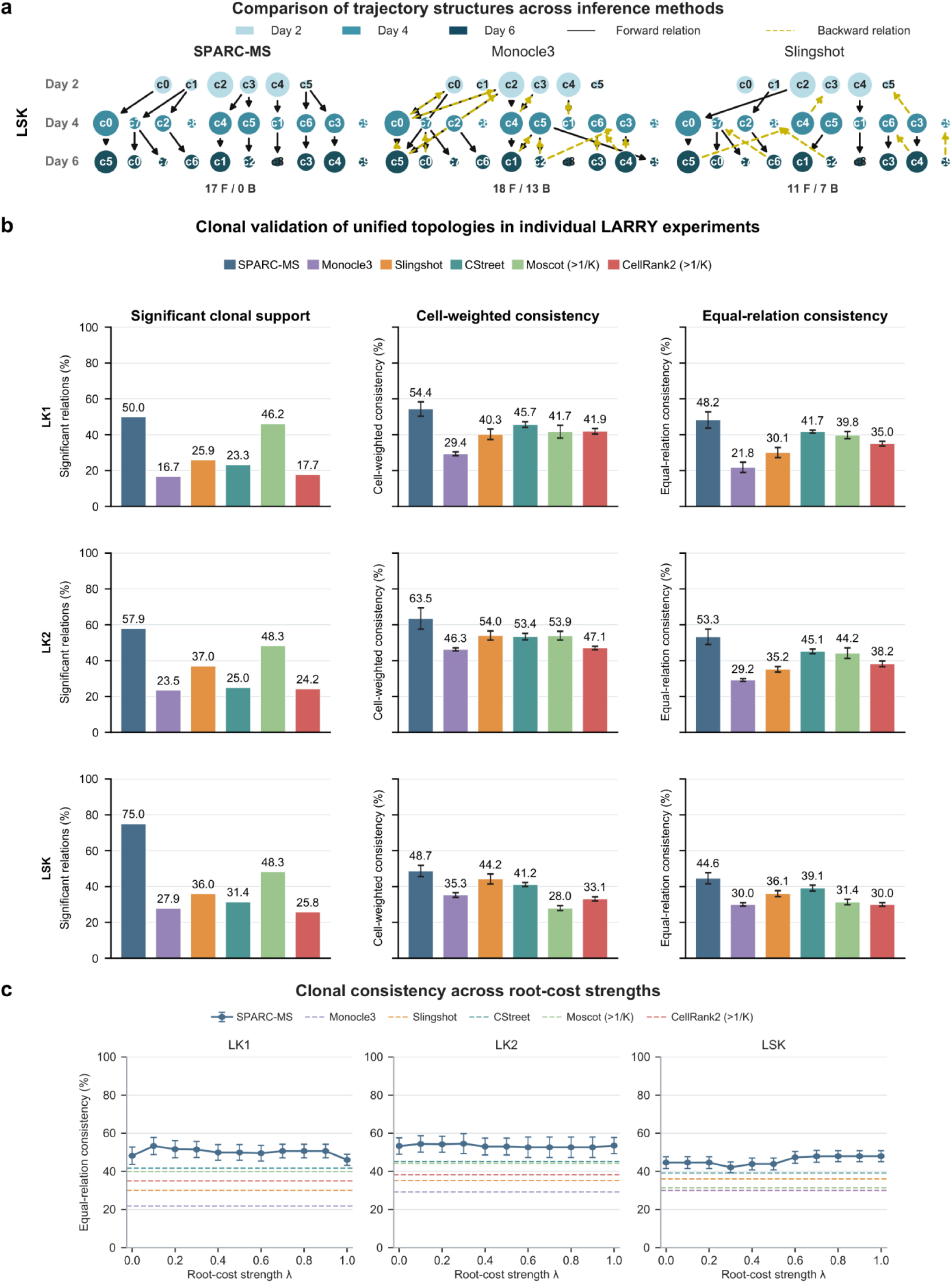
LARRY topology, clonal comparisons and optional source preference. (a) Multi-source SPARC (SPARC-MS), Monocle 3 and Slingshot LSK relation graphs. Node size denotes cell number and color denotes day; solid black and dashed yellow lines show forward and backward relations, with F and B counts. (b) LK1, LK2 and LSK comparisons of significant clonal support, cell-weighted consistency and equal-relation consistency for six methods. Fold error bars on consistency metrics show sample standard deviation (SD) across five clone-evaluation partitions. Complete native outputs retain their differing relation counts and within-day availability; backward relations receive zero directional support. (c) Equal-relation clonal consistency across optional source-preference strengths from 0 to 1 in increments of 0.1. SPARC-MS uses the same 1,000 complete edge-weight vectors at every setting, with round-matched CytoTRACE2 root medians. Points and error bars show means and sample SD over the five shared clone-evaluation partitions. Dashed lines show the corresponding comparator means from panel b. All candidate day-2 roots remain eligible; strength zero is the main SPARC-MS analysis. Clone labels enter evaluation after route inference.

**Extended Data Fig. 4.**
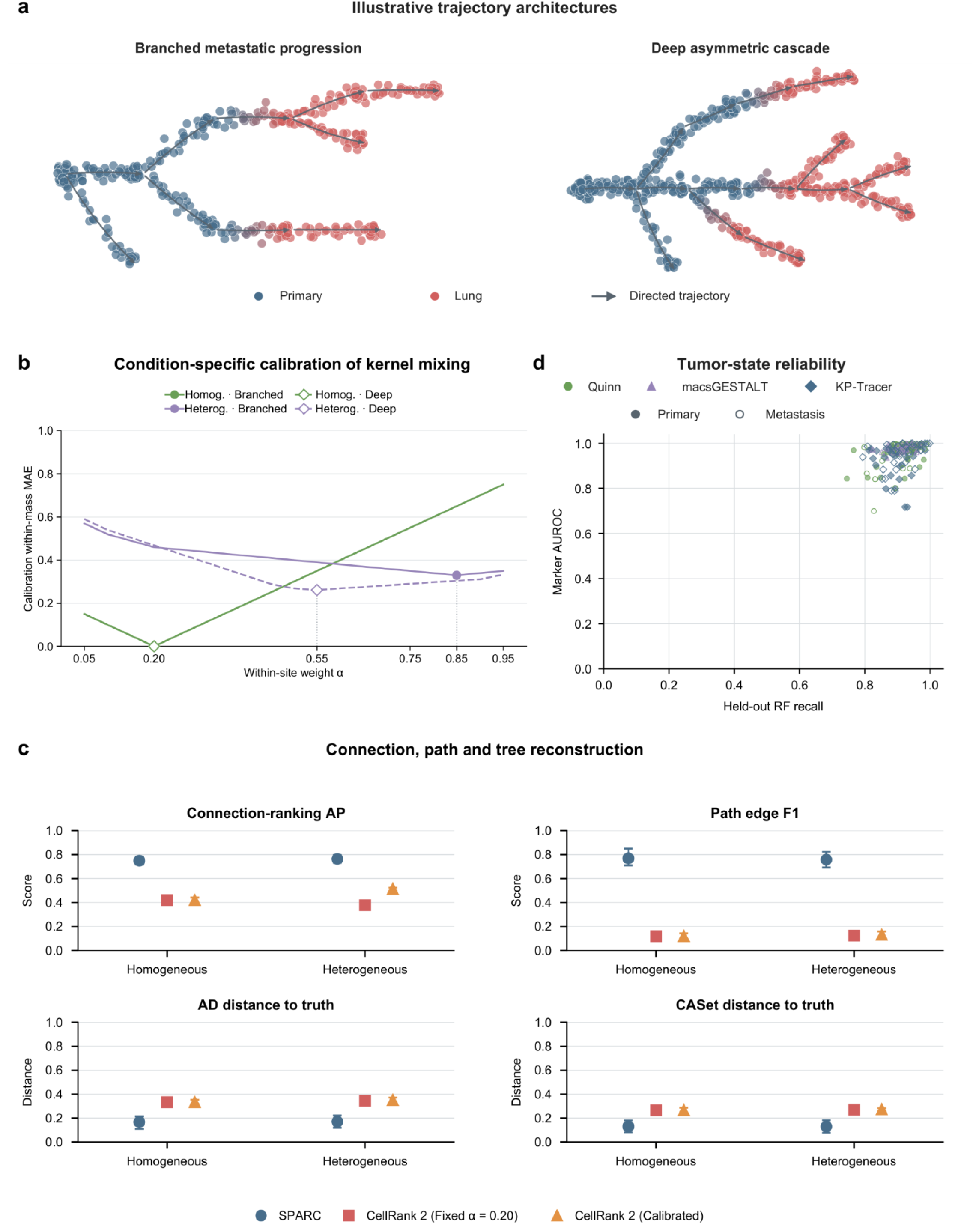
Generative trajectory reconstruction, RNA-state reliability and calibration. (a) Simplified illustrative layouts of Branched metastatic progression and a Deep asymmetric cascade, with schematic point-cloud coordinates and density. The actual simulations use complex 15-state and 18-state lineages, each with four direct primary-to-lung routes. (b) Primary-state within-mass mean absolute error across alpha on the calibration datasets, aggregated by topology and mode. Homogeneous selects alpha = 0.20; Heterogeneous selects 0.85 for Branched and 0.55 for Deep. (c) Comparison of connection-ranking average precision (AP), root-to-terminal path edge F1, ancestor–descendant (AD) distance and common-ancestor-set (CASet) distance for SPARC and fixed or calibration-selected CellRank 2. CellRank 2 used 50 within-compartment neighbors. The same calibrated configuration is transferred to all four readouts. Means and 95% intervals retain all 160 test datasets and use five independent seed-level factorial means. Higher AP/F1 and lower AD/CASet denote better reconstruction. (d) Real-tumor nearest-competitor marker AUROC versus held-out random forest recall; each point is an evaluable RNA-defined state. Symbols and colors identify dataset family and primary/metastatic compartment.

**Extended Data Fig. 5.**
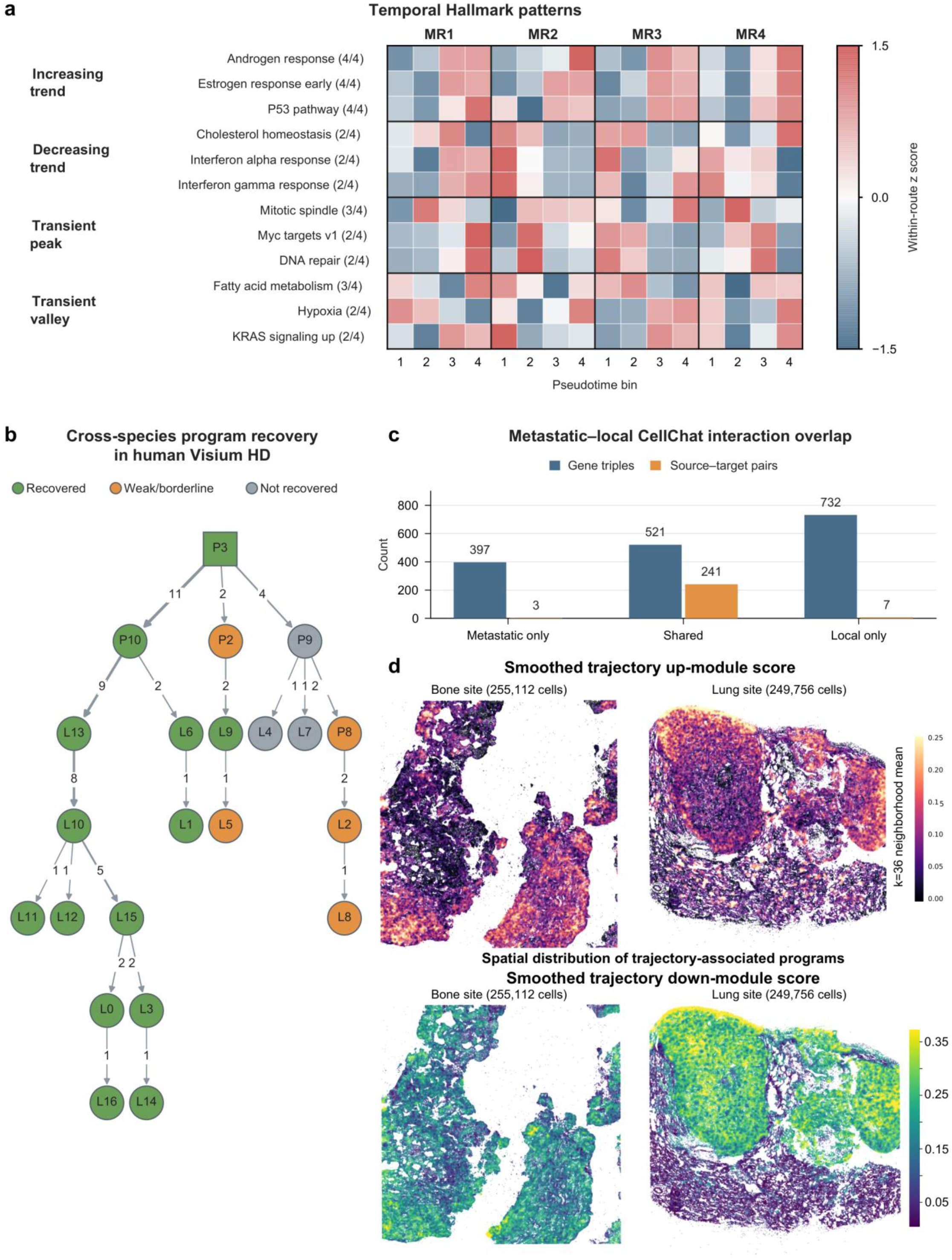
Local/metastatic programs and human spatial correspondence. (a) Four-bin Hallmark activity profiles along MR1-MR4, grouped as increasing, decreasing, transient-peak or transient-valley patterns. Colors show within-route standardized activity; numbers in parentheses give the number of routes supporting the indicated pattern out of four. Root P3 cells are excluded from this analysis. (b) Recovery of selected ortholog-matched mouse state programs in human Visium HD, displayed on the metastatic subgraph. (c) Overlap of CellChat ligand-receptor gene triples and source-target cell-type pairs between local and metastatic program analyses. (d) Increasing and decreasing full-terminal trajectory-module scores in Bone and Lung, smoothed over 36 spatial neighbors. These modules derive from nine complete lung-terminal routes including post-anchor states. Bone and Lung are from different patients.

**Extended Data Fig. 6.**
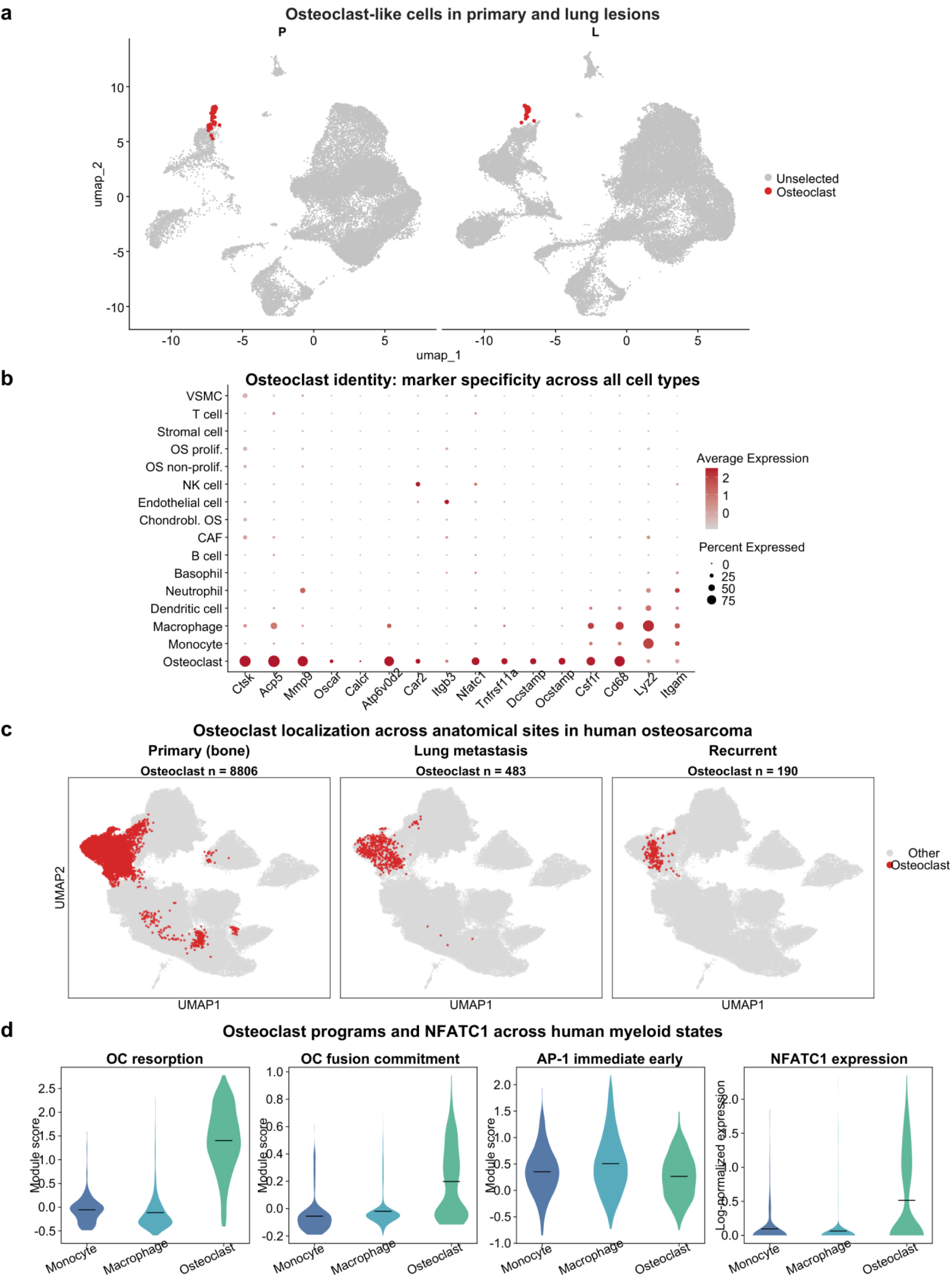
Osteoclast-like identity in mouse and human osteosarcoma. (a) Osteoclast-annotated mouse cells highlighted in primary and lung UMAPs. Full annotations contain 114 primary and 122 lung cells. (b) Canonical osteoclast and myeloid markers across mouse cell types; color shows mean expression and dot area detection fraction. (c) Our reanalysis of osteoclast-annotated cells in human primary bone, lung-metastatic and recurrent scRNA-seq samples from GSE152048, with counts of 8,806, 483 and 190. (d) Resorption, fusion/commitment, AP-1 and NFATC1 expression in human lung monocyte, macrophage and osteoclast annotations.

**Extended Data Fig. 7.**
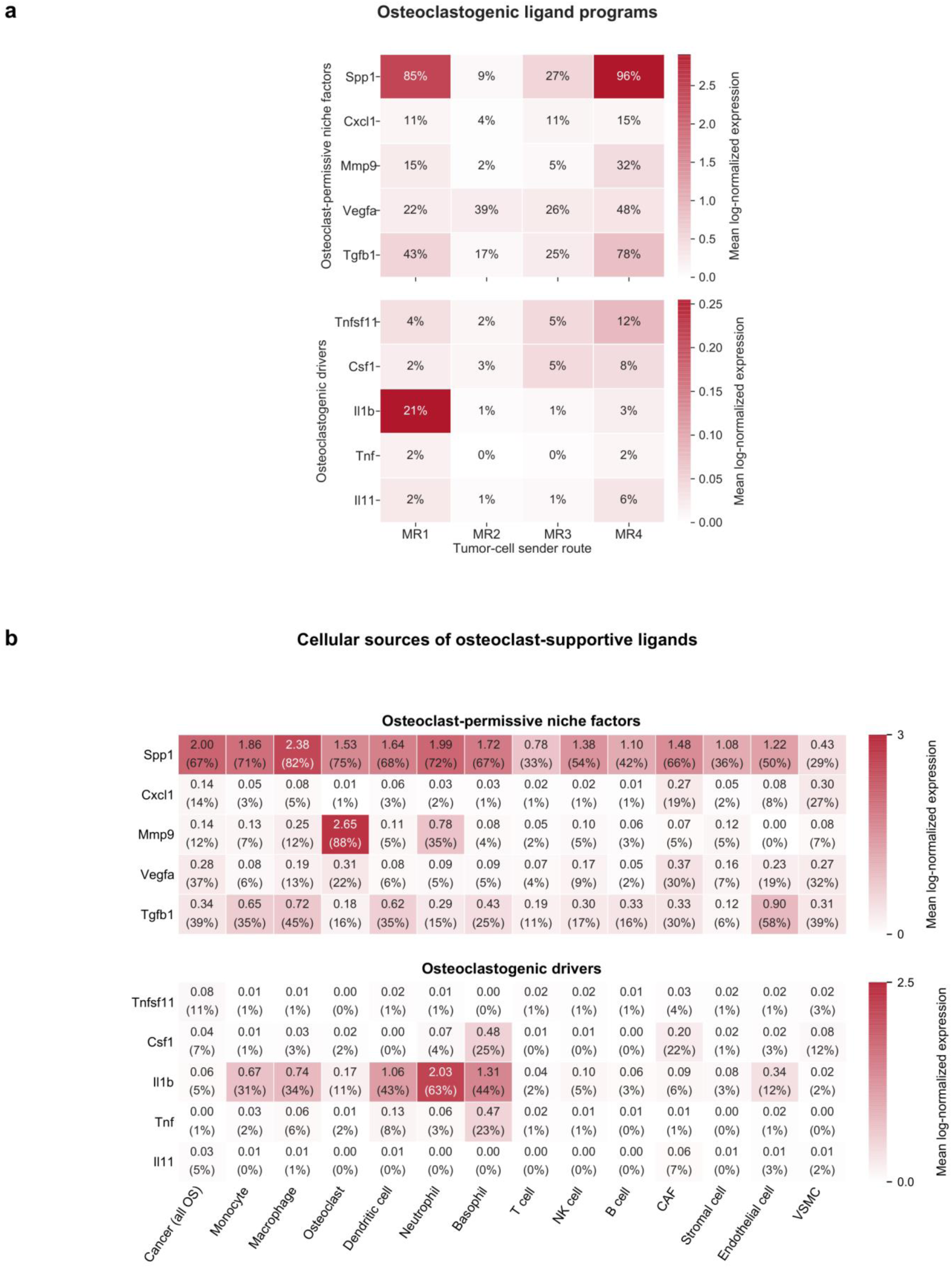
Route-associated ligands and their cellular sources. (a) Expression of osteoclast-permissive niche factors and osteoclastogenic ligand transcripts across MR1-MR4. (b) Expression of the same ligand classes across malignant, stromal and immune populations. Colors show mean log-normalized expression and numerical percentages show detection fractions. The two ligand classes have separate expression scales.

**Extended Data Fig. 8.**
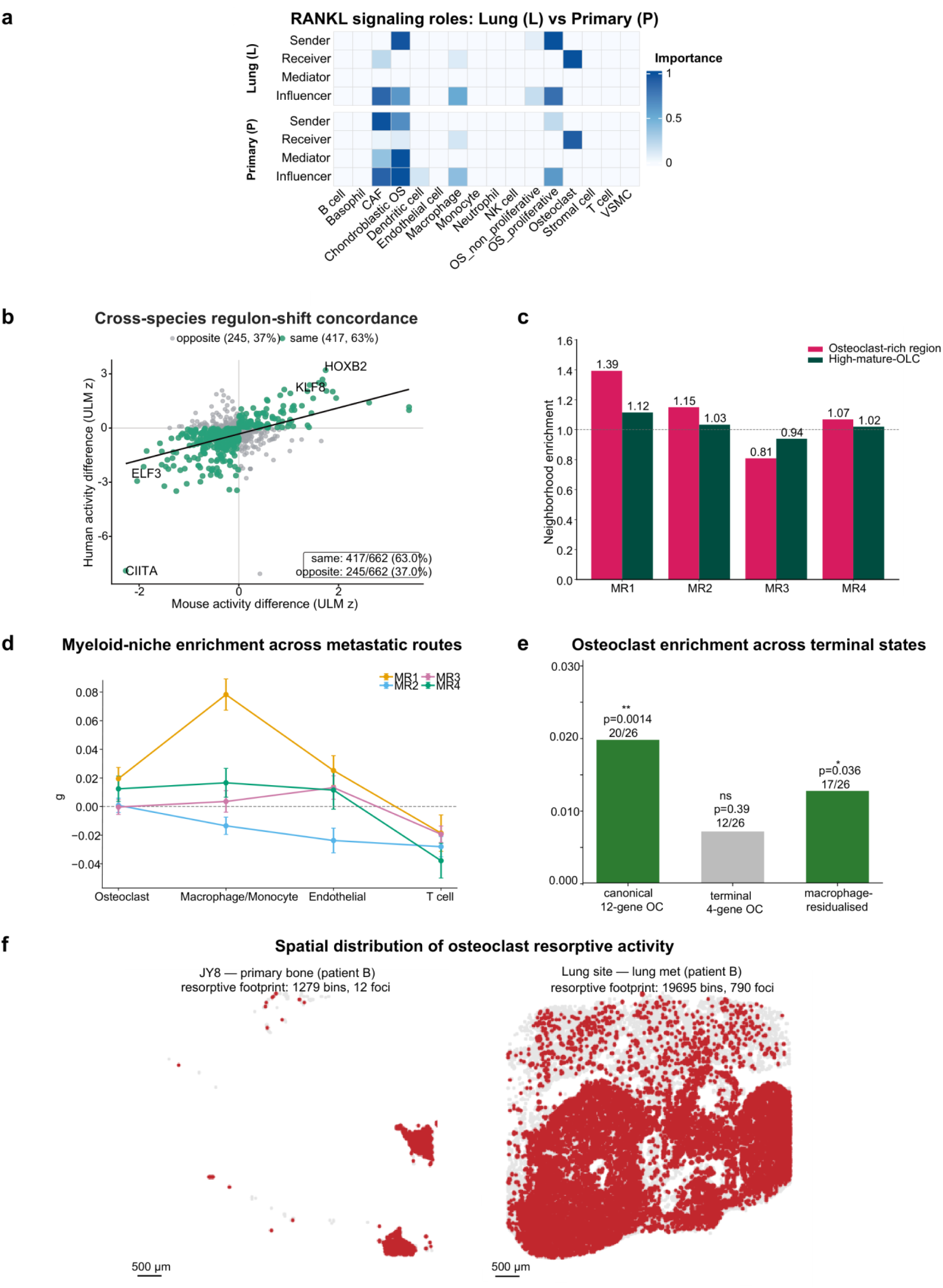
Niche signaling, regulatory concordance and spatial validation. (a) Predicted RANKL sender, receiver, mediator and influencer roles in mouse lung and primary conditions. Each role uses a common display normalization across the two regions. (b) Mouse-human concordance of univariate linear model (ULM) transcription factor (TF) activity contrasts. Each point is one symbol-matched TF; the axes show the difference between mean osteoclast-like and monocyte activity in each species. 417 of 662 contrasts share direction and 245 have opposite direction. Activity scores are ULM slope t-statistics. (c) Lung neighborhood enrichment of objects assigned to the osteoclast-rich region and objects with high mature-resorptive scores near MR1–MR4-associated malignant cells. The reference is tumor cells assigned to the other routes. (d) Mean route enrichment in osteoclast-, macrophage/monocyte-, endothelial- and T-cell-high spots in 26 external GSE293065 conventional Visium sections; error bars show standard error of the mean (SEM) across sections. (e) MR1-minus-MR3 osteoclast-niche enrichment using canonical twelve-gene, terminal four-gene and macrophage-residualized scores in the same 26 sections. Bars show mean section-level differences. Two-sided paired Wilcoxon signed-rank tests give P = 0.0014439, 0.3940727 and 0.0357244, respectively; MR1 has the higher value in 20, 12 and 17 of 26 sections. Displayed P values are unadjusted; **P < 0.01, *P < 0.05, ns P ≥ 0.05. (f) RNA-defined mature-resorptive footprints in the limited paired primary region and the Lung specimen: 1,279 bins in 12 foci and 19,695 bins in 790 foci. Scale bars, 500 µm.

**Extended Data Fig. 9.**
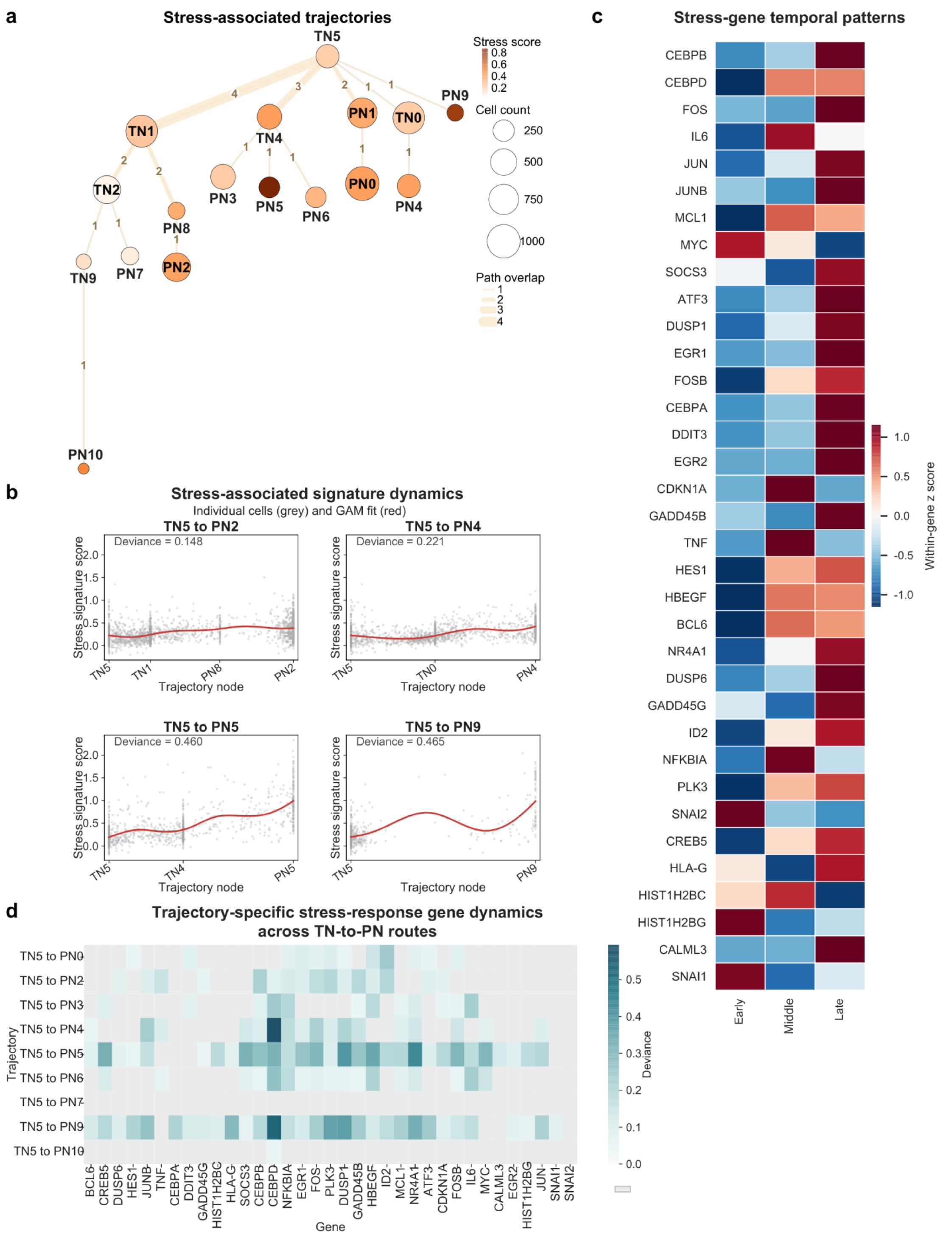
Route-specific stress-response dynamics before and after chemotherapy in high-grade serous ovarian cancer. (a) TN5-rooted single-source SPARC (SPARC-SS) trajectories with node area denoting cell number, node color denoting stress score and edge width denoting destination-path overlap. (b) Stress-signature scores in individual cells and generalized additive model (GAM) fits along four representative treatment-naive to post-treatment routes; labels give explained deviance. (c) Early, middle and late stress-gene profiles, standardized within each gene. (d) Route-specific stress-gene dynamics across treatment-naive to post-treatment paths, colored by explained deviance.

**Extended Data Fig. 10.**
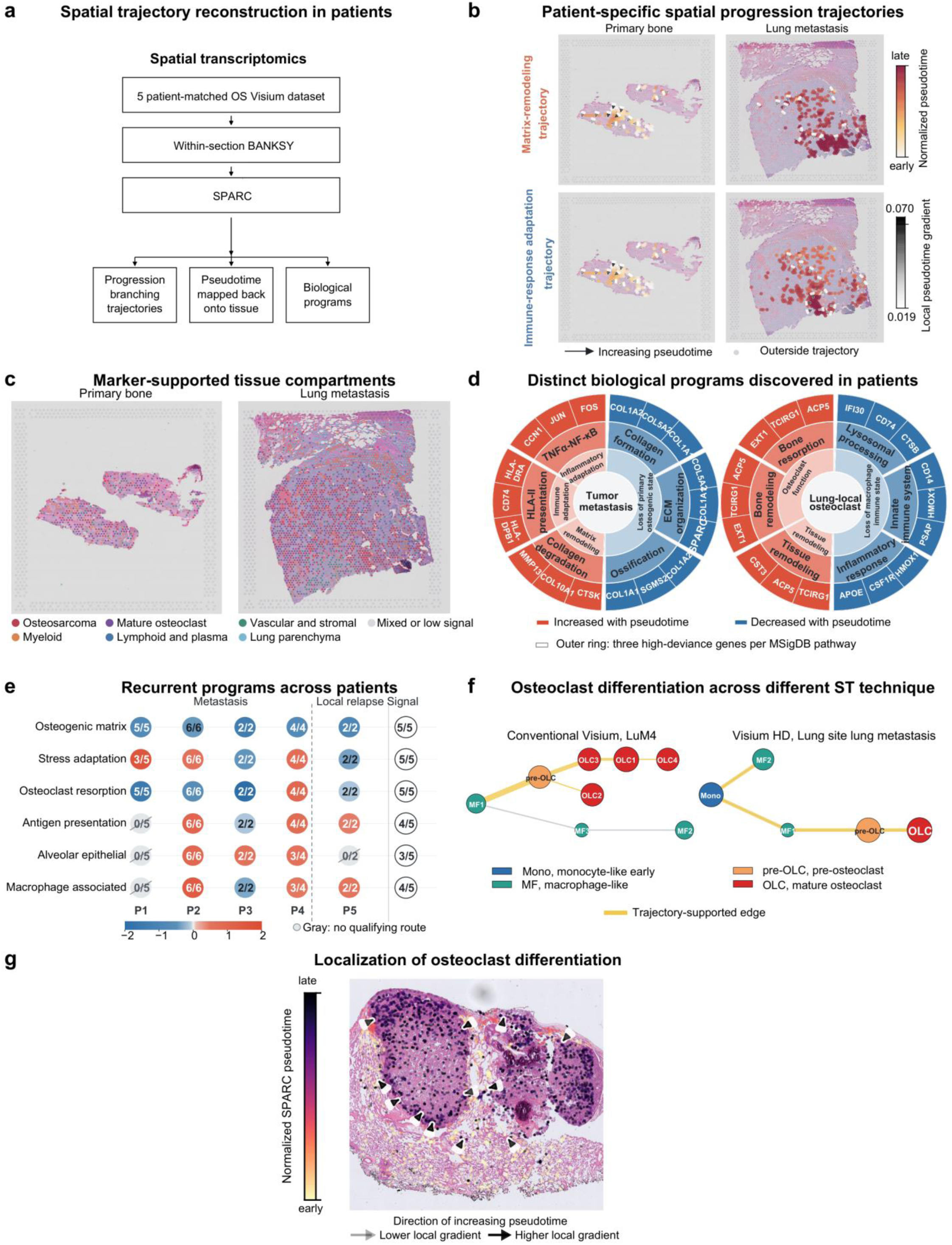
SPARC reconstructs tumor progression and osteoclast-associated trajectories from spatial transcriptomic data. (a) Schematic of SPARC analysis in five paired conventional Visium datasets. (b) Matrix-remodeling and immune-response adaptation trajectories in a paired primary/lung example, overlaid on tissue images. Points show route pseudotime and arrows the local increasing-pseudotime gradient. (c) Marker-supported tissue compartments. (d) Increasing and decreasing programs from tumor-metastasis and lung-local osteoclast trajectories; the outer ring lists three high-deviance genes per selected MSigDB pathway. Functional labels describe the contributing subsets of the KEGG antigen-processing-and-presentation pathway. (e) Program recurrence across four primary-to-lung pairs and one primary-to-local-relapse pair; fractions count qualifying patient-specific routes, with a separate patient-level summary. (f) Conventional Visium and Visium HD lung myeloid/osteoclast-associated maps, using the displayed monocyte (Mono), macrophage (MF), pre-OLC and osteoclast-like cell (OLC) identities. (g) Lung osteoclast-associated pseudotime on the tissue image; arrows indicate local increasing-pseudotime direction.

