## Supplementary Notes for "SPARC Reveals Distinct Osteoclastogenesis-Associated Malignant States in Metastatic Osteosarcoma"

### Supplementary Methods

### 1 Study design and expression representation

SPARC uses molecular similarity to connect cluster states within an experimentally specified progression. A primary-to-metastatic application asks how states can be reached from a nominated primary root. A time course asks which of the earliest observed states provides the least-cost route to a later state. These questions are implemented as single-source SPARC (SPARC-SS) and multi-source SPARC (SPARC-MS). In both modes, edges quantify transcriptomic support for connections between cluster states under the experimental constraints.

The inputs are a cell-by-feature matrix, a shared latent representation, cell identifiers, condition or sampling-time labels, and fixed within-condition cluster assignments. RNA datasets were normalized and represented by principal components. Harmony correction was applied only with the batch structure appropriate to the experiment[1]. For lineage-tracing time courses, cells were represented in a joint PCA space, Harmony was run separately within each time point, and corrected coordinates were recentered to the original time-specific PCA centroid. This procedure preserves the mean displacement between sampling times. For spatial datasets, BANKSY representations incorporated neighborhoods within individual tissue sections[2]. Dataset-specific settings are described below.

Leiden clustering was performed separately within each condition[3]. Cluster identifiers encode sampling time or site and a cluster index, such as D2_c7. Cell-type annotations were added for biological interpretation. Barcode sequences, clone identifiers and clone-derived fate annotations were excluded from clustering, latent representations and path inference.

### 2 Connection strength and the allowed directed graph

#### 2.1 Observed and expected cell-neighbor connections

Cluster-center distances summarize average molecular positions, while cell-neighbor links retain local connections across population boundaries. PAGA formalized a graph-abstraction approach based on intercluster links relative to a random expectation[4]. SPARC uses this rationale to measure within- and cross-condition connection strengths in one cluster-excluded neighbor graph. The expected-link calculation below is conditioned on excluding each query cell's own cluster. For a fixed cluster partition, the graph was constructed in latent space. We denote the conventional kNN neighborhood parameter by $k$ and the number of neighbors selected after excluding the query cell's own cluster by $k_{\text{out}}$. For each cell in cluster $i$, the algorithm selected its $k_{\text{out}}$ nearest cells among all cells outside cluster $i$, without a time-specific or condition-specific quota. The LARRY edge-resampling analyses used the first 30 latent coordinates and $k_{\text{out}} = 49$, preserving the 49 non-self neighbors represented by the stored Scanpy[5] setting of $k = 50$ when the query cell is included.

Directed selections were symmetrized by union. A cross-cluster cell pair was counted once if either cell selected the other. Let $N$ be the total number of retained cells, let $n_i$ and $n_j$ be the retained sizes of clusters $i$ and $j$, and let $U_{ij}$ be the number of unique unordered cross-cluster pairs in this union graph. Under a reference model in which different cells independently select $k_{\text{out}}$ distinct cells uniformly from outside their own cluster,

$$p_i = \frac{k_{\text{out}}}{N - n_i}, \qquad p_j = \frac{k_{\text{out}}}{N - n_j}.$$

For one cell in $i$ and one cell in $j$, the probability that at least one direction selects the other is $p_i + p_j - p_i p_j$. The expected unique-pair count and the symmetric adjacency count are therefore

$$E(U_{ij}) = n_i n_j (p_i + p_j - p_i p_j), \qquad V_{ij} = 2U_{ij}, \qquad E(V_{ij}) = 2E(U_{ij}).$$

Connection strength is

$$w_{ij} = \frac{2U_{ij}}{2E(U_{ij}) + 10^{-8}}.$$

The observed and expected terms use the same symmetric adjacency convention. The factors of two cancel apart from the numerical stabilizer. This exclusion-aware observed-to-expected construction follows the graph-abstraction rationale of PAGA[4]; the retained SPARC ratio is uncapped. The dimensionless ratio measures relative connection strength under the exclusion-aware null. Its reciprocal defines the traversal cost, so stronger connections have lower costs. The candidate universe was fixed from every condition-permitted cluster pair, including pairs with zero full-data connection strength. These candidates remain columns of the complete resampling vector. Only positive finite connection strengths are traversable in a particular graph; zero-weight connections are unavailable in that round.

#### 2.2 Selection and sensitivity of $k_{\text{out}}$

When an upstream graph reports a neighborhood size $k$ that includes the query cell, SPARC uses $k_{\text{out}} = k - 1$ to preserve the number of non-self neighbors, with all selected neighbors drawn from outside the query cell's cluster. When the reported size already counts non-self neighbors, that count can be carried forward directly.

For a new dataset, $k_{\text{out}}$ should be prespecified from the representation-graph design, independently of lineage labels and downstream agreement scores. Its selection should account for the eligible pool $N - n_i$ for each cluster $i$, latent-space density, cluster imbalance and separation, which together affect the distance to the $k_{\text{out}}$th external neighbor. The necessary feasibility condition is $k_{\text{out}} \leq \min_i (N - n_i)$ in every analyzed resample; in practice, the value should be comfortably below this bound so that isolated clusters are not forced to use very distant cells. The application settings below used $k_{\text{out}} \in \{19, 29, 49\}$; these values are recorded analysis choices for the corresponding representations.

A new application should repeat the complete inference over a prespecified smaller and larger value and compare edge support, reachability, selected source, modal route and conditional path frequency. In the full LARRY sensitivity analysis, $k_{\text{out}} \in \{49, 99, 199\}$ was evaluated in all three datasets using 1,000 paired 80% within-cluster cell-resampling rounds per setting. All 59 requested endpoints remained reachable. Relative to $k_{\text{out}} = 49$, all 59 empirical modal routes were unchanged at 99 and 57 of 59 were unchanged at 199. The temporally penalized equal-edge clonal-consistency score was unchanged across the grid for LK2 and spanned 1.50 and 2.70 percentage points for LK1 and LSK, respectively. These results support stability over the tested LARRY range. New datasets require the same prespecified sensitivity analysis, with any material changes reported directly.

##### 2.3 Permissible directions

Let the ordered conditions be $T_0, \ldots, T_m$. Supported within-condition connections are reciprocal, including those in $T_0$. Across successive conditions, only the earlier-to-later orientation is allowed. The candidate connection universe is fixed before resampling and includes every condition-permitted cluster pair, including zero-valued candidates. The full supported graph enters inference. Reciprocal within-condition edges permit directed cycles, so SPARC operates on a general directed graph.

Only positive finite weights are traversable, with $C_e = 1/w_e$. A connection with $w_e = 0$ is absent in that round, leaving a disconnected destination unreachable when every permissible route is broken. Resampling retains the fixed condition-permitted pair universe, including zero-valued candidates.

##### 2.4 Formal application graph settings

All applications used 30 latent coordinates and graph-resampling seed 159. The cluster-excluded graphs used $k_{\text{out}} = 49$ for the LARRY, osteosarcoma, HGSOC and myeloid OLC analyses, 29 for the paired spatial and osteoclast HD analyses, and 19 for the osteoclast Visium analysis. LARRY used SPARC-MS with every day-2 cluster as a candidate source at zero starting cost; the other applications used SPARC-SS with a fixed root. Complete dataset-specific cell counts, cluster counts and root assignments are retained in the accompanying reproducibility configurations. Each application generated 1,000 complete edge-weight vectors at each of 70%, 80% and 90% cell retention, with 80% as the primary fraction.

The time-ordered simulation analyses used $d = 30$, $k_{\text{out}} = 49$, a single supplied initial root and seed-specific expression inputs. Graph resampling used seed 159; simulation-generation and cell-selection seeds are recorded separately. Each formal simulation unit used its complete selected cell set. Its 80% uncertainty run contained 1,000 rounds. Simulation truth was read only during scoring.

The reproducibility configurations also record the selected-display source policies and path-resampling seeds. Internal cluster identifiers and displayed labels, and all inference destinations versus selected biological-route subsets, are supplied separately.

#### 3 Source policy and shortest-path inference

##### 3.1 SPARC-SS

A root is supplied before inference, based on the experimental question and, where appropriate, developmental-potential scoring. CytoTRACE2 was applied to RNA counts to obtain cell-level potency scores[6]. Median scores within eligible initial-condition clusters were used for root nomination. The osteosarcoma analysis fixed P3 as the root. A single-source Dijkstra search[7] retrieves a least-cost path to every reachable requested destination. All non-root clusters may be destinations, including other primary-site clusters. Destination is used here as a graph-analysis term and includes internal biological states.

##### 3.2 SPARC-MS

All clusters in $T_0$ are candidate sources. A multi-source Dijkstra search selects a source and a path for each later-condition destination by minimizing total path cost. With source cost $R_r$,

$$\pi^*(t) = \operatorname*{argmin}_{r\in\mathcal{R},\, \pi:r\to t} \left[ R_r + \sum_{e\in\pi} C_e \right].$$

For the main analysis, $R_r = 0$. Each later destination receives one least-cost source per round, and candidate sources with no optimal routes are not selected as origins for any destination. An equivalent construction adds a virtual source with edges of weight $R_r$ to all candidate roots and removes that virtual node from reported paths. With zero source costs and strictly positive biological-edge costs, traversing an edge between two candidate sources cannot improve upon starting at the second source. The optimized zero-cost implementation can therefore omit those initial-to-initial edges during the search without changing the optimum. They remain part of the conceptual allowed graph and are included when source costs differ.

##### 3.3 Optional developmental-potential preference for SPARC-MS route selection

Users may enable a developmental-potential preference when higher estimated potency should influence the selected origin. All $T_0$ clusters remain eligible sources, and the source preference is added to the RNA path cost.

In round $b$, the original per-cell CytoTRACE2 scores are aggregated over the cells retained in each root:

$$q_{r,b} = \text{median}_{i\in C_r\cap S_b}\, \text{CytoTRACE2Score}_i.$$

The scores are centered and scaled across the $m_{\mathcal{R}} = |\mathcal{R}|$ candidate roots:

$$\mu_b = m_{\mathcal{R}}^{-1} \sum_r q_{r,b}, \qquad \sigma_b = \left[ m_{\mathcal{R}}^{-1} \sum_r (q_{r,b} - \mu_b)^2 \right]^{1/2}, \qquad \tilde{q}_{r,b} = \frac{q_{r,b} - \mu_b}{\sigma_b}.$$

When $\sigma_b$ is numerically zero, all standardized values and source costs are set to zero. Otherwise,

$$R_{r,b} = \lambda_{\text{root}} \left( \max_{r'} \tilde{q}_{r',b} - \tilde{q}_{r,b} \right), \qquad \lambda_{\text{root}} \geq 0.$$

The root medians use exactly the cells that generated the corresponding complete edge-weight vector. Each round re-aggregates the original CytoTRACE2 cell scores over its retained root cells. A one-standard-deviation advantage in root potency reduces its starting cost by $\lambda_{\text{root}}$ path-cost units. The main analysis uses $\lambda_{\text{root}} = 0$.

**Cost scale and an initial preference strength.** For a user enabling the preference on the tested LARRY representation, $\lambda_{\text{root}} = 0.1$ provides a practical initial setting. Its nonzero source costs and baseline RNA route costs were of similar order of magnitude: For LK1, the median nonzero source cost was 0.220 and the median RNA cost was 0.215; 1/20 modal routes changed. For LK2, the median nonzero source cost was 0.227 and the median RNA cost was 0.170; 1/19 modal routes changed. For LSK, the median nonzero source cost was 0.109 and the median RNA cost was 0.426; 0/20 modal routes changed. Source-cost medians pool nonzero candidate-root costs across the 1,000 rounds and roots; RNA-cost medians pool the selected $\lambda_{\text{root}} = 0$ route costs across rounds and destinations. Modal-route changes compare the most frequent ordered route for each destination between strengths 0 and 0.1. Thus 57 of 59 modal routes were retained. Individual round–destination paths changed in 5.68%, 4.44% and 1.29% of LK1, LK2 and LSK cases, respectively. The coefficient expresses a cost trade-off whose influence depends on the potency and RNA-cost differences between competing routes. Applications with different route-cost distributions require their own scale and sensitivity checks.

**Source-preference sensitivity and clonal evaluation.** The displayed sensitivity analysis uses $\lambda_{\text{root}} \in \{0, 0.1, \ldots, 1.0\}$ and the same 1,000 complete edge-weight vectors at every strength. Round-matched root medians supply the corresponding entry costs, and exact Dijkstra selects routes under the resulting objective. Empirical modal routes define the selected relation graph. Clonal consistency is calculated using the same within-day and across-time relation definitions and five shared clone-evaluation partitions as the principal LARRY comparison. Points show means with sample SD over those partitions, and comparator reference lines are their corresponding fixed native-graph means. Clone labels are used after route inference for these evaluations. The principal LARRY result retains equal-cost sources; positive strengths describe the optional developmental-potential preference.

### 4 Empirical edge and path uncertainty

#### 4.1 Cluster-stratified cell subsampling

Cell subsampling has been used to evaluate trajectory stability and uncertainty in inferred pseudotime[8–10]. For each fraction $f \in \{0.7, 0.8, 0.9\}$, 1,000 samples were generated. The primary fraction was 0.8. Within each original cluster, $\max(2, \lfloor f n_i \rfloor)$ cells were selected without replacement; the analyzed clusters were large enough to satisfy this sampling rule. The same selected cells were used for every connection involving that cluster. Labels, latent coordinates, directional constraints, source policy and the allowed edge universe were held fixed. Sampled cluster sizes, cluster-excluded selections, union counts and exclusion-aware expected counts were recomputed in every round before calculating $w$.

Exact outside-cluster neighbor ranks were cached in the fixed latent space. Filtering a cell's outside-cluster rank list to retained cells and selecting its first $k_{\text{out}}$ retained neighbors reproduces an exact cluster-excluded neighbor search on that subsample when the cache is sufficiently deep. Cache depth was checked for every row and round, and an insufficient cache triggered failure. The cache stores cell-to-cell ranks from batched squared Euclidean distances; intercluster connectivity continues to arise from counted cell pairs. Full-data reconstruction checks verified the cached exact-neighbor calculations.

Each round generated a complete ordered vector $\mathbf{w}_b = (w_{1b}, \ldots, w_{Eb})$. All 1,000 vectors were saved per fraction, including zeros. Histograms and summaries were computed from these vectors without fitting a Gaussian or other parametric family. A zero mass, skewness or a small-cluster tail therefore remained part of the empirical distribution.

##### 4.2 Joint and independent-marginal path calculations

The principal analysis ran exact Dijkstra once on each saved complete vector after applying $C_{eb} = 1/w_{eb}$ to positive weights. A common sampling index across all edges preserves dependencies induced by the same retained cells. A control independently selected a saved row for each distinct connection and combined those values into a synthetic vector. Reciprocal orientations of the same connection shared the selected row. Differences between this control and the joint-vector analysis describe sensitivity to inter-edge dependence; the joint-vector analysis remains the principal uncertainty estimate.

For destination $t$, let $I_{tb}$ indicate reachability and let $n_t(\pi)$ count rounds selecting a particular ordered node sequence. Reported quantities were

$$\widehat{P}_t(\text{reachable}) = \frac{\sum_b I_{tb}}{B}, \qquad \widehat{P}_t(\pi) = \frac{n_t(\pi)}{B}, \qquad \widehat{P}_t(\pi \mid \text{reachable}) = \frac{n_t(\pi)}{\sum_b I_{tb}}, \quad B = 1000.$$

Reachability, conditional frequency and unconditional frequency were reported separately. A destination with zero reachable rounds has an undefined conditional frequency. The most frequent complete path was the empirical modal path. The 95% set was the smallest frequency-ranked prefix attaining cumulative conditional frequency at least 0.95. The top-20 listing and the empirical 95% path-coverage set were calculated independently. All reported frequencies characterize perturbation sensitivity for the fixed input design.

Results were compared across sampling fractions using selected path identities, source choices, reachability and conditional frequency summaries. A trajectory map combines modal paths that can arise in different sampled graphs. Terminal-path usage is the number of displayed destination-specific paths traversing an edge and serves as a route-overlap summary.

#### 5 Expression-based cluster reliability

##### 5.1 Held-out prediction of the original labels

LARRY LK1, LK2 and LSK were analyzed separately at each time point, and osteosarcoma was analyzed separately in primary and lung conditions. Random-forest inputs consisted of the first 30 coordinates of the original Harmony representation, with fixed original clusters as targets[11]. Barcode and outcome columns were not supplied. Five stratified outer folds were generated with seed 20260918. Each forest used 300 trees, minimum leaf size 2, square-root feature sampling and seed 20260918 plus the outer-fold index. Within the outer training set, three-fold sigmoid calibration was performed with an ensemble of fitted/calibrated models[12]. Outer test cells were excluded from both fitting and calibration.

For cluster $c$, recall was the number of outer-test cells with original label $c$ predicted as $c$, divided by all outer-test cells carrying label $c$. Per-cell class scores, a pooled out-of-fold confusion matrix and ten-bin top-label calibration summaries were retained. Low-confidence cells had maximum class score below 0.6; boundary cells had a difference below 0.2 between the two largest scores. These cutoffs summarize assignment ambiguity. Outer folds held out cells within each experiment, defining a within-study assessment of expression-derived label reproducibility.

#### 5.2 Reclustering and frozen nearest-competitor markers

Within each condition, 70%, 80% and 90% of all cells were sampled without replacement for 100 rounds per fraction. This sampling is condition-wide, in contrast to the cluster-stratified edge-weight experiment. The original latent representation was fixed. Neighbor graphs and Leiden clusters were recomputed with the original condition-specific settings: LARRY used 30 neighbors, 30 coordinates, resolution 0.4, seed 0 and the igraph implementation with two iterations; osteosarcoma used 50 neighbors, 30 coordinates, resolution 1.0, seed 159 and the leidenalg implementation. Sampling seeds began at 20260918. Original and resampled clusters were aligned by cell membership. Each original cluster was matched to the new cluster maximizing Jaccard overlap on the retained cells; matching was performed independently for each original cluster, so merged matches were possible. Jaccard overlap served only to align cluster identities; the molecular-reliability readout came from held-out marker preservation.

For marker discovery, each original cluster was divided approximately equally into discovery and evaluation cells using a reproducible hash-derived seed based on 20260922 and the cluster identifier. Its nearest competitor was the other original cluster with the nearest discovery-cell centroid in the 30-dimensional latent space. Gene expression came from the saved nonnegative log-normalized RNA matrix. Vector-associated features were excluded; eligible genes were detected in at least 1% of discovery cells. For each direction of the cluster-versus-competitor contrast, up to ten markers were selected by discovery mean-expression difference, requiring a difference of at least 0.15, a detection-fraction advantage of at least 0.05 and detection in at least 10% of the higher-expression group. At least three markers in each direction were required. All marker thresholds apply to the saved log-expression scale.

Gene means and standard deviations were fitted using discovery cells, with standard deviations floored at 0.10. The frozen per-cell program score was

$$M_i = \frac{1}{|G_+|} \sum_{g \in G_+} \frac{x_{ig} - \mu_g}{s_g} - \frac{1}{|G_-|} \sum_{g \in G_-} \frac{x_{ig} - \mu_g}{s_g}.$$

For each reclustering round, only retained evaluation cells were scored in the matched cluster $C'$ and matched nearest competitor $N'$. If these matched to the same new cluster, or either group contained fewer than ten evaluation cells, the contrast was unavailable. AUROC was the fraction of $C'$–$N'$ cell pairs in which $M_{C'} > M_{N'}$, with half credit for ties. Here AUROC has a defined binary classification interpretation. Mean-score difference and the fraction of marker directions preserved in the held-out expression contrast were also retained. Five detection-decile-matched random gene sets provided a descriptive marker control. The score evaluates preservation of the frozen nearest-competitor molecular contrast. The evaluation cells were held out from marker selection and scaling, but participated in the original representation and in the reclustering step.

##### 5.3 Expression-state reliability in tumor Cas9 datasets

The formal tumor Cas9 benchmark contained 14 complete units, 121,885 cells and 172 fixed states. RF prediction used the frozen 30-dimensional PCA coordinates separately within the primary and metastatic compartments, with the same five outer folds, three-fold sigmoid calibration, 300 trees, minimum leaf size two and square-root feature sampling specified above. Nearest-competitor marker discovery used only the 2,000 formal RNA highly variable genes. Discovery and evaluation cells were split equally within each fixed state, and bidirectional marker panels were frozen before transfer evaluation.

Each compartment underwent 100 unstratified 80% cell subsamples without replacement. Neighbor graphs were reconstructed with 30 neighbors, and Leiden used resolution 0.6 and random seed 159. States with fewer than 50 cells were assigned to the nearest larger same-compartment state. Original and resampled states were matched by maximal sampled-cell membership overlap. The held-out marker comparison required distinct matched groups and at least ten evaluation cells in each. Merged groups and unavailable marker panels were recorded as unavailable. The plotted marker AUROC was the median over evaluable rounds. Lineage files were reserved for the separate clonal benchmark.

#### 6 Cell-level pseudotime and molecular changes

##### 6.1 Whitened projection onto a selected route

TSCAN provided an earlier use of projecting cells onto cluster-level paths for pseudotime reconstruction[13]. Cells in the selected route's clusters were projected onto the ordered cluster-centroid polyline. For latent matrix $X$, covariance was regularized as $\widehat{\Sigma} = \mathrm{Cov}(X) + 10^{-6}I$. Eigenvalues were floored at $10^{-8}$ when forming the inverse square root. Cells and centroids were transformed by the same whitening matrix. For a cell $x$ and consecutive whitened centroids $a, b$, the segment coordinate was

$$u = \mathrm{clip}_{[0,1]} \frac{(x-a)^\top (b-a)}{\|b-a\|^2}.$$

The segment with the smallest projection residual was selected, with zero-length segments handled as points. The cell's distance was the cumulative length before that segment plus $u\|b-a\|$. Distances were normalized within the route. This construction yields a unitless relative position along the chosen molecular route.

##### 6.2 Gene dynamics and enrichment

GAM-based modeling of expression along pseudotime has been used in TSCAN and tradeSeq[13;14]. For routes containing at least 30 cells, the retained gene-expression workflow fits a Gaussian generalized additive model[15], $x_{ig} = \beta_{0g} + f_g(\tau_i) + \epsilon_{ig}$, using ten cubic-spline basis functions and smoothing penalty 0.8. Explained deviance summarizes the strength of the fitted expression pattern. Smooth-term significance estimates are corrected with the Benjamini-Hochberg procedure within each trajectory[16]. The formal application tables reported here were generated with CPU pyGAM[17] and use its smooth-term p-value.

Direction-specific dynamic-gene sets were tested against GO and KEGG annotations[18;19]. For an eligible background of $M$ genes, a pathway containing $G$ background genes and a selected list of $n$ genes with overlap $g$, the one-sided enrichment probability is

$$P(X \geq g) = \sum_{x=g}^{\min(n,G)} \frac{\binom{G}{x}\binom{M-G}{n-x}}{\binom{M}{n}}.$$

The background was analysis-specific: tested and mapped genes where configured for spatial trajectory enrichment, the union of annotated genes in the fixed GMT for retained mouse Hallmark/canonical ORA, and the annotated g:Profiler[20] domain for GO. Enrichment backgrounds were defined separately for each analysis, as specified below. Gene identifiers and species mappings were reconciled before enrichment using the recorded annotation resources, including MyGene.info[21]. Thresholds and gene-set versions specific to the spatial applications are given below. Gene-dynamics and enrichment results used the ordered route definitions, cell selections and analysis-specific backgrounds specified below.

#### 6.3 Complete topology and shared metastatic trunks

The complete osteosarcoma display used the P3-rooted 80% empirical modal paths, estimated with 1,000 complete-vector resampling rounds. It retained 31 nodes and 30 union edges from paths to 17 lung states and 13 non-root primary states. A requested graph destination is not necessarily a terminal biological state. Full-topology path-overlap labels count destination-specific paths using an edge.

For the primary-versus-metastatic program comparison, primary routes ended at the primary branch tips P0, P13, P5, P1 and P12. P7 is an intermediate on the P5 route. The P12 route contains 3,060 cells and 22,489 tested genes. Metastatic analysis used four shared trunks: MR1, P3→P10→L13→L10; MR2, P3→P9→P8→L2; MR3, P3→P9→L4; and MR4, P3→P2→L9. These anchors delimit the shared process before further branching. MR1 includes L13→L10. Post-anchor states remain available in the complete topology but are excluded from these four shared-trunk analyses. Primary, metastatic and common masks identify membership in the selected path sets within this complete topology.

MR1–MR4 provide a focused comparison across four distinct primary states at the inferred primary-to-lung transition: P10, P8, P9 and P2, respectively. Their complete sequences were selected in 100%, 100%, 99.2% and 100% of the 1,000 rounds at 80% cell retention. All rounds were reachable for these destinations. Additional branches from P10 and P9, including routes to L6 and L7, were examined in the extended gene-program comparison provided through the study repository. This extension supplies complete gene-dynamics and pathway-scoring tables while retaining the four-route definitions used for the main comparison.

All 22,489 eligible expression features were evaluated per selected route. The primary core contained 1,185 increasing and 786 decreasing genes. Shared sets contained 209 increasing and 480 decreasing genes, and metastatic-associated sets contained 469 increasing and 1,194 decreasing genes. Within the primary and metastatic groups separately, an up- or down-regulated core required adjusted

GAM $P < 0.05$ and the same direction on every selected route in that group. Common-up and common-down sets are the respective intersections of the two group cores. Metastatic-associated up and down sets are the metastatic core minus the corresponding primary core. These set differences define the route-program membership used for enrichment analysis.

For the comparative top-100 enrichment lists, genes additionally passed the retained sparse-expression gate of at least 20 expressing cells and 5% detection on every relevant trajectory. Lists were ranked by the recorded pooled mean explained deviance. Separate branch-core lists retained the original top-200 policy. Local mouse Hallmark[22] and canonical-pathway ORA used the union of genes annotated in the corresponding fixed GMT collection as the hypergeometric background; queries were intersected with that union. GO analysis used g:Profiler's annotated domain without a supplied custom background. These backgrounds differ from the tested-gene background used in the paired spatial trajectory enrichment and are reported separately.

For repeated pathway terms in Fig. 4b, functional descriptors in parentheses were assigned from the contributing gene subsets. The source tables retain the original pathway identifiers, gene membership and enrichment statistics.

#### 6.4 Gene temporal classes and representative GAM curves

Temporal classification used the four selected metastatic trunks, including root P3 cells, and 20 equal-width normalized pseudotime bins. Mean expression per bin was calculated for genes significant at adjusted GAM $P < 0.05$ and explained deviance $\geq 0.02$; empty bins were linearly interpolated. A gene had to pass this gate in at least two trunks. A peak in bins 0–5 was Early, in bins 6–13 Transient, and in bins 14–19 Late. A consensus label required at least two concordant votes and no tie for the leading class. Genes without a consensus label remained unassigned in the source table.

The representative display applied a stricter illustration policy: each candidate was significant and class-concordant in all four trunks. Candidates were ranked by decreasing minimum explained deviance across routes, with gene symbol as the tie-break. The first candidate whose fitted GAM peak matched the assigned class in all four routes was retained. H19, Dlk1 and H2afj represent Early, Transient and Late, respectively, in 12 curves. Representative curves were fitted with ten cubic-spline basis functions and smoothing penalty 0.8. These examples illustrate early, transient and late expression patterns along the inferred routes. Temporal-class Hallmark ORA used the class-specific gene lists and the fixed mouse Hallmark GMT annotated-gene union as background, followed by BH correction.

#### 6.5 GSVA temporal activity-shape heatmap

Pathway temporal shapes were analyzed separately from the gene-peak classes and gene-list ORA. Root P3 cells were excluded and the remaining cells in each shared trunk were partitioned into four stable rank/quantile bins. Retained route-bin pseudobulk expression entered R GSVA[23] with the fixed mouse MSigDB 2026.1 Hallmark, canonical-pathway and immunologic collections, `kcdf="Gaussian"`, gene-set sizes 10–500, `absRanking=FALSE` and `maxDiff=TRUE`; computation used `SerialParam`. The retained source records include the R session, pseudobulk cell counts, complete

GSVA matrices and selected pathway lists. For one pathway and route, let $\rho$ be the Spearman correlation of activity with bin index, $\Delta$ the mean of bins 3–4 minus bins 1–2, and $c$ the mean of bins 2–3 minus bins 1 and 4. Increasing required $\rho \geq 0.6$ and $\Delta > 0$; decreasing required $\rho \leq -0.6$ and $\Delta < 0$. A transient peak required $|\rho| < 0.6$, $c > 0$ and a maximum in bin 2 or 3; a transient valley required $|\rho| < 0.6$, $c < 0$ and a minimum in bin 2 or 3. A pathway required support for its block pattern in at least two of four routes.

For the compact illustration, at most three of the 50 Hallmark pathways per class were selected. At least two formally supporting routes also required Pearson correlation $\geq 0.5$ with the appropriate template and an original across-16-column row-z range $\geq 0.25$ within the route. Templates were $[-1.5, -0.5, 0.5, 1.5]$ and its reverse for increasing/decreasing, and $[-1, 1, 1, -1]$ and its negative for peak/valley. Candidates were ranked by clear-route count, mean similarity across all four routes, similarity in clear routes, median source amplitude in clear routes and pathway symbol. Displayed pathways satisfied both the classification criteria and the template-based selection criteria. All four routes, including nonsupporting routes, remain shown.

For display, activity was standardized separately for each pathway and route across its four bins using the sample standard deviation (ddof = 1). Colors denote these within-route standardized activity scores. The original GSVA values and across-16-column normalization were preserved as separate source tables.

Temporal classification and enrichment analyses used the specified shared-trunk routes, cell selections and expression inputs.

### 7 External clonal-lineage validation

#### 7.1 Data identity and independent evidence

LARRY LK1, LK2 and LSK[24] were analyzed separately at days 2, 4 and 6, with 28,994, 36,740 and 72,946 expression cells. LK1 and LK2 share the day-2 pool and are partially shared culture series, not wholly independent biological replicates. Each experiment used its complete expression input. Barcode sequences, clone identifiers and barcode-derived fate labels were excluded from expression representations, clustering, source selection and path inference.

#### 7.2 Native within- and across-time relation sets

The selected comparison evaluates the native chosen topologies of SPARC-MS, Monocle 3, Slingshot, CStreet, Moscot and CellRank 2. Across-time relations retain direction; opposite within-day directions collapse to one unordered pair. SPARC, Monocle 3 and Slingshot contribute their native same-day relations. CStreet contributes native across-time relations and per-timepoint Otsu-thresholded within-day graphs. Moscot is a between-time transport method with no native within-day topology and contributes zero within-day relations in every experiment. Each method contributes its complete native relation set to the comparison. Graph density and available relation types were retained in each method's evaluated output.

##### 7.3 Lineage-informative incidences

For a forward relation $i \to j$, an informative target cell in $j$ has a clone detected somewhere at the source day; it is supported if that clone is detected in $i$. For an unordered within-day pair $\{i, j\}$, supported incidences include cells in $i$ whose clone is detected in $j$ and cells in $j$ whose clone is detected in $i$. An endpoint cell is informative if its clone is detected somewhere outside its own cluster at that day. Each endpoint cell contributes at most once for a particular relation. The relation rate is supported divided by informative incidences, with an undefined rate when no informative incidences exist.

Native backward across-time relations retain their barcode-observability denominators but have zero supported incidences. They contribute zero to equal-relation consistency even when the fold lacks informative targets. Forward and within-day rates without informative cells remain unavailable. Shared clones support common ancestry and clonal consistency across the connected states.

##### 7.4 Three readouts, permutation tests and pooling

SPARC's pooled native output contained 59 relations: 54 across time and five within a day. Of these, 36 showed significant clone enrichment (61.0%).

For significant clonal support, the observed clone-sharing rate was tested against 1,000 permutations of clone labels within sampling-day and library strata. Barcode coverage and clone-size distributions were preserved by permuting existing labels. The empirical one-sided P value used the add-one correction. One BH family per method and experiment spanned the evaluable within- and across-time relations. A supported relation required adjusted $P < 0.05$ and observed-to-permutation enrichment greater than one. Metric A divides the number of significantly supported relations by every selected native relation, including unsupported backward and unavailable relations.

Let $s_{rb}$ and $n_{rb}$ be supported and informative incidences for relation $r$ in clone fold $b$, and $\mathcal{R}_b$ its evaluable relation set together with native backward relations scored zero. Cell-weighted consistency and equal-relation consistency are

$$B = \frac{1}{5}\sum_{b=1}^{5}\frac{\sum_r s_{rb}}{\sum_r n_{rb}}, \qquad C = \frac{1}{5}\sum_{b=1}^{5}\frac{\sum_{r\in\mathcal{R}_b} r_{rb}}{|\mathcal{R}_b|}.$$

The pooled B/C values combine all three experiments at the incidence or relation unit within each fold before averaging the five folds. Metric A pools the numbers of significantly supported and selected relations across the three experiments. Per-experiment results were also reported. Bars show percentages; B/C error bars show sample standard deviation across the five clone-evaluation folds.

All cells from a clone remain in one of five shared evaluation folds. Multitime clones were shuffled with seed 20260925; remaining clones used 20260925+7777, with cyclic fold assignment. Permutation seeds were 20261003 plus 1,000 times the experiment index. Inference was already fixed before fold/permutation scoring. The biological sampling units are the culture series described above; fold and permutation variation was calculated within these experiments.

### 8 Trajectory-method comparison

All methods received the complete LK1, LK2 or LSK cell input and the shared 50-dimensional within-day Harmony representation. The fixed reporting clusters supplied a common state vocabulary for evaluating each output. Method-specific dimensional reductions, source policies and relation-selection rules are specified in the following method descriptions. Slingshot and Monocle 3 used the prespecified day-2 roots D2_c2, D2_c6 and D2_c0 for LK1, LK2 and LSK, respectively. SPARC-MS treated all day-2 clusters as candidate sources with zero starting cost.

#### 8.1 SPARC-MS relation graph

SPARC-MS used the first 30 latent coordinates, $k_{\mathrm{out}} = 49$ and the complete condition-permitted cluster-pair universe. Each of 1,000 rounds sampled 80% of cells within every original cluster, recalculated the complete connectivity vector and ran exact Dijkstra. For every later-day destination, the most frequent ordered route was identified. The union of these modal routes defined the reported cluster-relation graph, including its selected within-day connections.

#### 8.2 Slingshot and Monocle 3

Slingshot[8] used all 50 Harmony coordinates and the fixed cluster labels with `getLineages`, rooted at the specified day-2 cluster. R seed 0 was used. A failed default distance calculation triggered the `dist.method="simple"` fallback. Consecutive cluster pairs in the returned rooted lineages defined the selected relations.

Monocle 3[25] received the same latent coordinates in its PCA slot and constructed a two-dimensional UMAP with minimum distance 0.3. `cluster_cells` and `learn_graph(use_partition=FALSE)` then produced the principal graph. The expression assay served as a container for this supplied-coordinate analysis. Within the root cluster, the cell nearest its UMAP centroid was located; its nearest principal vertex rooted the graph. Principal vertices were assigned the majority fixed reporting-cluster label of projected cells. Root-to-leaf shortest paths on the principal graph were converted to cluster sequences, with missing labels removed and consecutive duplicate labels collapsed. Consecutive pairs yielded the relation graph. Native backward relations were retained in the evaluation.

#### 8.3 CStreet

CStreet v1.1.2[26] used the Harmony coordinates as its input features and the supplied cluster assignments, with additional normalization and log transformation disabled. Within- and between-time calculations used ten internally reduced components, neighborhood size 15, five probability-sampling trials and seed 159. Across-time relations were taken from its native Otsu-thresholded state graph with maximum outdegree ten. For within-day relations, all non-self ordered cluster pairs were considered; missing native scores were assigned zero, and the Otsu threshold was calculated separately for each day. An unordered pair was retained when either native direction met that threshold. This uses the method's state-probability outputs for both relation classes.

##### 8.4 moscot and CellRank 2

moscot[27] fitted unbalanced transport between days 2 and 4 and between days 4 and 6 in the 50-dimensional Harmony representation. It used entropic regularization 0.005, marginal relaxation 0.95 for both distributions, mean cost scaling, convergence threshold $10^{-3}$ and at most 5,000 iterations. For source cluster $s$ and target cluster $t$, cell-pair coupling masses were summed to $M_{st}$. Within each source cluster and adjacent-day window, $p_{st} = M_{st}/\sum_u M_{su}$. The selected output consisted of across-time relations satisfying the above-uniform rule defined below.

CellRank 2[28] constructed a RealTimeKernel from these adjacent-day transport couplings, with within-time connectivity weight 0.2, 30 neighbors in the Harmony representation and seed 159. The reported output was derived from its cell-transition matrix. For each source cluster, transition probabilities into a target cluster were averaged over source cells. The resulting cluster-level entries were normalized over the next-day targets for across-time selection. Within a day, self mass was excluded and each row was renormalized over the other same-day clusters. An unordered pair was retained when either normalized direction passed its above-uniform threshold.

##### 8.5 Above-uniform relation selection

For the target set $\mathcal{T}_b$ with $K_b = |\mathcal{T}_b|$,

$$p_{st} = \frac{M_{st}}{\sum_{u\in\mathcal{T}_b} M_{su}}, \qquad \text{retain } s \to t \quad \text{if } p_{st} > \frac{1}{K_b}.$$

Here $M$ denotes transport mass for moscot and cluster-aggregated transition weight for CellRank 2. The reference distributes mass equally across target clusters and adapts to their number, providing a common reporting rule for the two probabilistic outputs. For within-day CellRank 2 relations, the target set excludes the source cluster, so $K_b = K_{\text{day}} - 1$. Selection was completed before barcode evaluation. The separate clone-label permutation analysis supplied the clonal-significance assessment.

Across-time relations retained native direction. Within-day pairs were represented as unordered relations, with opposite orientations collapsed. Native graph density, source policy and availability of within-day relations were retained in the comparison. For time-directed methods, zero backward transitions follow the prescribed temporal constraints. Independent lineage information evaluates the biological support for the inferred relationships.

#### 9 Tumor Cas9 lineage-tracing benchmark

##### 9.1 Datasets and expression-derived states

The tumor lineage-tracing benchmark comprised 14 complete primary-metastasis analysis units from 11 independent tumor families. Eight units came from the KP-Tracer lung adenocarcinoma study[29], two from the macsGESTALT pancreatic cancer study[30], and four cell-number versions came from the Quinn lung cancer xenograft study[31]. The four Quinn units arose from one tumor family and were grouped together for uncertainty estimation and calibration. Each unit retained all available cells.

Counts were normalized to 10,000 per cell and log1p-transformed. Two thousand highly variable genes were selected by the Seurat v3 procedure on raw counts, and 30 principal components defined the expression representation. Primary and metastatic cells were clustered separately using 30 neighbors, Leiden resolution 0.6 and seed 159. A state with fewer than 50 cells was assigned to the nearest state containing at least 50 cells within the same compartment, using the PCA centroid distance. Primary states were denoted $P_i$ and metastatic states $M_j$. SPARC used $k_{\text{out}} = 49$, the public `sparc.run` v0.3.0 implementation and the complete permitted graph. Within-compartment connections were reciprocal and cross-compartment connections were directed from primary to metastasis. These connections formed the graph supplied to SPARC.

##### 9.2 Direction-constrained Cas9 relation scores

Lineage information entered after expression-based relation scoring. For each unit, the published conditional lineage-transition matrix was converted to one score for every state pair. Let $L_{ij}$ denote its directional entry. The benchmark score was

$$g_{ij} = \begin{cases} L_{ij}, & i \in P,\ j \in M, \\ \frac{1}{2}(L_{ij} + L_{ji}), & i,j \in P \text{ or } i,j \in M. \end{cases}$$

Thus, primary-to-metastasis pairs retained the experimentally defined forward direction, while a within-compartment pair was treated as an unordered association. The evaluated universe contained every within-primary, primary-to-metastasis and within-metastasis state pair. SPARC supplied its cluster-excluded global-kNN connection strength $w_{ij}$ for the corresponding pair. Per-unit agreement was Spearman correlation between $g_{ij}$ and the expression-derived relation score over the full pair universe. Correlation over primary-to-metastasis pairs alone was retained as a prespecified control.

##### 9.3 CellRank 2 within-time weight grid

CellRank 2 v2.2.0[28] was run with independent within-compartment kNN transition matrices and an adjacent-compartment moscot coupling[27]. The main fixed and calibrated comparisons used 50 neighbors within each compartment; the alpha-sensitivity curves also show the 30-neighbor control. The moscot coupling used entropic regularization 0.005, marginal relaxation parameters 0.95, mean cost scaling, convergence threshold $10^{-3}$ and at most 5,000 iterations. The converged coupling and both within-compartment matrices were computed once per unit.

After aggregation to state pairs, let $q_{ij}^{P}$, $q_{ij}^{PM}$ and $q_{ij}^{M}$ denote the within-primary, primary-to-metastasis and within-metastasis components. For $\alpha \in \{0.05, 0.10, \ldots, 0.95\}$, the combined score was

$$q_{ij}^{(\alpha)} = \begin{cases} \alpha q_{ij}^{P}, & i,j \in P, \\ (1-\alpha) q_{ij}^{PM}, & i \in P,\ j \in M, \\ q_{ij}^{M}, & i,j \in M. \end{cases}$$

The all-block score therefore changes the relative placement of within-primary and cross-compartment relations as $\alpha$ varies. Multiplication of every cross-compartment score by the same positive factor leaves the cross-only rank correlation unchanged, providing the prespecified control. The $\alpha = 0.50$ setting was the prespecified CellRank 2 analysis.

##### 9.4 Topology sensitivity and family-held-out calibration

For unit $u$ with $K_u$ states, the topology-sized diagnostic set $E_u^{(\alpha)}$ contained the $K_u - 1$ highest-scoring CellRank 2 state pairs. Ties were resolved by source and target identifiers. Turnover from the prespecified reference was

$$T_u(\alpha) = 1 - \frac{|E_u^{(\alpha)} \cap E_u^{(0.50)}|}{K_u - 1}.$$

This definition provided a rank-based, fixed-size topology diagnostic. The SPARC edge ranking is independent of $\alpha$, so its turnover curve is zero by construction.

For family-held-out calibration of family $f$, $\alpha$ was estimated from the remaining families, which were first given equal weight, and the value maximizing their mean all-block Spearman correlation was chosen:

$$\widehat{\alpha}_{-f} = \arg\max_{\alpha} \frac{1}{|\mathcal{F} \setminus f|} \sum_{h \neq f} \frac{1}{|\mathcal{U}_h|} \sum_{u \in \mathcal{U}_h} \rho_u(\alpha).$$

Ties were resolved by proximity to 0.50 and then by the smaller value. The selected value was applied to every unit in family $f$. This calibration uses Cas9 evidence from the training families and was analyzed as a CellRank 2 sensitivity comparison. SPARC relation scores and routes were computed from RNA-derived connection strengths.

Curves and point estimates used equal family weighting. Ninety-five percent intervals were obtained from 10,000 family-cluster bootstrap samples with seed 159. Reported sample counts distinguish the 14 analysis units from the 11 independent tumor families.

#### 10 Mixed within- and cross-compartment simulation benchmark

##### 10.1 Formal factorial design and representation

PROSSTT[32], Splatter[33], TedSim[34] and dyngen[35] generated expression on two specified topologies: Branched metastatic progression (15 states) and a Deep asymmetric cascade (18 states). Both complex lineages contain four direct primary-to-lung routes. Each was evaluated under two transition-mass modes, two expression regimes and eight seeds, giving $4 \times 2 \times 2 \times 2 \times 8 = 256$ complete datasets: 96 for calibration and 160 for held-out evaluation. The modes were Homogeneous (`homogeneous_020`) and Heterogeneous (`metastasis_state_specific`). Each benchmark unit contained 8,000 cells and 2,000 simulated features, with 4,000 cells at each site. State labels supplied by the simulation were retained for scoring and cluster-level inference, so recovery is conditional on these state definitions supplied as the common inference input.

Counts were total-count normalized to 10,000, log1p-transformed, and represented by 30 principal components after Seurat-style highly variable feature selection (up to 2,000 features). The preprocessing seed was 159. PROSSTT used branch length 30, 12 modules, alpha 0.3 and beta

3. Splatter used a pool multiplier of 2.5, 100 path steps and differential expression probabilities 0.20/0.12 for standard/hard. dyngen used 100 housekeeping genes, 128 simulations, census interval 1, edge-balanced snapshot sampling and bandwidth 0.1. TedSim used 16,384 division-tree tips, 20 cell-identity factors, 10 differential factors, state-tree edge length 2 and maximum state walk 6. Simulator-specific sampling schedules and implementation details are provided with the simulation configurations and analysis code. The parameters above define the benchmark settings.

The standard/hard expression settings used signal scales 1.0/0.55, dispersion 20/7.5, dropout midpoint $-2/0$ and batch shift 0/0.35. Per-state abundances, the primary root P0, lineage edges and lung destinations are specified in the supplied 256-dataset configuration. Validated simulator-native state pools were reused where the generation parameters and state coverage were unchanged. For each descendant-allocation setting, state sampling, RNA observation generation, PCA and trajectory inference were performed on the resulting datasets. Formal SPARC used the `sparc.run` v0.3.0 implementation, single-source P0, $k_{\text{out}} = 49$, the complete allowed graph, and 1,000 joint-vector 80% within-cluster resampling rounds with seed 159. All non-root states were inference destinations; path scoring used the specified lung terminal subset. Truth edges were read only for evaluation.

##### 10.2 Lineage topology and prescribed transition-mass estimands

The complex reference state lineage specified the four Lung descendant subtrees and the precursor/intermediate states used for evaluation. The prescribed transition-mass matrix and generated abundance pattern were related design inputs. Homogeneous assigned 1,000 Lung cells to each complete subtree; its eligible non-final source states retained true within-block mass 0.20. Heterogeneous direct-edge masses were 0.85, 0.55, 0.25 and 0.05. They generated complete-subtree totals of 2,000, 1,294, 588 and 118 Lung descendants, respectively. Integer totals used the deterministic largest-remainder rule; cells within each subtree followed the specified relative state allocation. These allocations entered RNA observation generation before inference. The simplified illustrative layouts in Extended Data Fig. 4a summarize the organization of these lineages, with schematic point-cloud coordinates and density. The complete reference lineages, including all truth edges and state counts, are retained in the simulation configurations.

For a primary state, its within mass was divided among its true within-primary neighbors, and remaining mass among its configured lung targets. Lung states divided their mass among within-lung neighbors. These rules define a separate row-stochastic truth matrix. Full configuration records preserve the topology, abundances, within masses and cross targets for every unit.

##### 10.3 CellRank 2 mixing weights and calibration split

CellRank 2 used independently computed within-compartment connectivity kernels and an adjacent-compartment moscot coupling in the same PCA representation. The main simulation comparison used 50 within-compartment neighbors. SPARC selected 49 neighbors per cell from the global pool of cells outside the query cell's cluster. Moscot used entropic regularization 0.005, marginal relaxation 0.95, mean cost scaling, convergence threshold $10^{-3}$ and at most 5,000 iterations. Components were calculated once and recombined for 19 weights from 0.05 to 0.95:

$$T_\alpha = \begin{pmatrix} \alpha C_P & (1-\alpha)\Pi_{P\to L} \\ 0 & C_L \end{pmatrix}.$$

The final lung compartment retained its within-compartment kernel at unit weight. The reference weight 0.20 is the homogeneous-mass control; 0.50 is the topology-drift reference.

Seeds 159, 161 and 162 formed the 96-dataset calibration set, and seeds 164–168 formed the 160-dataset test set. The same eight seed values were used across the entire retained factorial design. For each simulator/topology/transition/expression condition, alpha minimized mean absolute error in total primary within mass on calibration seeds; ties selected the smaller alpha. That alpha was applied unchanged to the five test seeds. Alpha was selected by within-primary transition-mass MAE. The 32 condition-specific choices were 0.20 for Homogeneous, 0.85 for branched Heterogeneous and 0.55 for deep-asymmetric Heterogeneous. The mass oracle selected the mixing weight with the lowest true within-primary mass error for each test dataset; AP and path F1 were then evaluated using that weight. Metric-specific truth oracles select a mixing weight using the evaluation truth for each metric and are reported as diagnostic analyses. The held-out set contains 160 factorial datasets generated with five independent seed values. Means and 95% intervals used 10,000 seed-cluster bootstrap samples with seed 159, retaining condition pairing.

##### 10.4 Relation ranking and ordered-path recovery

The ranking universe contained unordered within-primary and within-lung state pairs, plus directed primary-to-lung pairs. Self-edges were excluded. Within-pair probability scores used the larger of the two directions; cross scores retained primary-to-lung direction. SPARC ranking used original connection strengths, while transition-calibration metrics used its row-normalized matrix. Both complete and block-specific average precision were calculated against the specified lineage-pair labels.

For a fixed-budget ranking diagnostic, the top $N$ pairs were retained, where $N$ was the number of true pairs in that scoring universe; score ties used source/target identifiers. With both sets of size $N$, pair F1 equals TP/$N$. Reciprocal within-pair orientations were restored for the directed HIM calculation. This truth-budget-matched diagnostic is distinct from the native, non-budget-matched LARRY topology comparison.

Root-to-terminal path recovery compared each inferred ordered path with the configured directed truth path. CellRank 2 paths were extracted from its aggregated state-transition matrix by minimizing the sum of edge costs $-\log\{\max(p_{ij}, 10^{-15})\}$ from the supplied root to each terminal, using positive-probability non-self edges. The all-state arborescence decoder described below was used for AD/CASet. Edge F1 was $2|E_{\text{pred}} \cap E_{\text{truth}}|/(|E_{\text{pred}}| + |E_{\text{truth}}|)$, averaged over the specified lung terminals. Exact sequence match was recorded separately. Primary-state within-mass MAE was the alpha-calibration loss, as specified above.

##### 10.5 All-state AD/CASet and alpha-dependent tree changes

The selected all-state comparison decoded each CellRank 2 matrix into a fixed-root maximum-probability arborescence, maximizing $\sum_{(i,j)\in T} \log p_{ij}$. Self-edges, zero-probability edges and incoming root edges were excluded. Node coverage, arborescence structure and root reachability were checked for every decoded graph. The union of SPARC paths to all non-root states was checked on the same state set. The tree-distance metrics were calculated on these validated benchmark trees. The comparator tree is defined by this maximum-probability arborescence objective. The optimum-branching step used Edmonds' algorithm[36] as implemented by NetworkX.

The tree-distance comparison adapts mutation-labeled tumor-tree measures to one uniquely labeled expression state per vertex, using the same complete state set $V$ and fixed root in truth and prediction. Following the ancestor-pair comparison of Govek et al.[37], define

$$\Phi(T) = \{(u, v) \in V^2 : u \neq v,\ u \text{ is ancestral to } v \text{ in } T\}.$$

The reported AD is the union-normalized variant

$$d_{\mathrm{AD}}(T, T') = \frac{|\Phi(T) \triangle \Phi(T')|}{|\Phi(T) \cup \Phi(T')|},$$

with zero assigned to an empty union. Self-pairs are excluded from both sets. For CASet-style distance[38], each state's ancestor set includes itself. For each unordered state pair, common-ancestor sets in truth and prediction were compared by Jaccard distance, and these distances were averaged across all state pairs. Equivalently, with $A_T(v)$ containing the focal state and all its ancestors, $C_T(u, v) = A_T(u) \cap A_T(v)$ and

$$d_{\mathrm{CASet}}(T, T') = \binom{|V|}{2}^{-1} \sum_{u<v} \frac{|C_T(u, v) \triangle C_{T'}(u, v)|}{|C_T(u, v) \cup C_{T'}(u, v)|}.$$

An empty common-ancestor union contributes zero. Lower values are better for both metrics. Per-unit oracle alpha minimizes the particular reported distance.

Topology drift used union-normalized AD from the same unit's alpha-0.50 tree; the fraction changed used exact directed-edge-set inequality. Across the 19-weight grid, 158 of 160 test CellRank 2 trees changed at least once. Seed-cluster uncertainty was retained for accuracy and drift. Pooled across the two modes, mean AD was 0.169 for SPARC and 0.342 for calibration-selected CellRank 2; mean CASet was 0.130 and 0.269, respectively. Mean AD drift from alpha 0.50 was 0.099 at alpha 0.05 and 0.249 at alpha 0.95. These measurements assess state-topology recovery under the supplied labels and decoding objectives. Metric-specific truth oracles use evaluation truth and remain separately marked as diagnostic analyses.

The relation-block analysis evaluated direct lineage-adjacency ranking within Primary, from Primary to Lung and within Lung separately. In Heterogeneous, within-Lung AP was 0.804 for SPARC and 0.658 for calibrated CellRank 2. The paired SPARC-minus-CellRank difference had a 95% seed-bootstrap interval of 0.137–0.158. These block-specific measurements complement the pooled connection, path and tree comparisons.

### 11 Time-ordered simulated topology and cell-number robustness

#### 11.1 Expression simulation

Splatter path simulation[33] generated four state graphs: linear progression, one bifurcation, two bifurcations and asymmetric branching. The topology-by-difficulty factorial contains four topologies, three difficulty settings and seeds 159, 160 and 161. Each dataset contains 6,000 selected cells and 2,000 genes, drawn according to a fixed state-abundance design from a 12,000-cell simulation. These selections define the formal benchmark dataset; all selected cells are passed to the compared methods. Common path parameters are 100 steps, skew 0.5, nonlinear probability 0.1 and sigma factor 0.8.

The easy, standard and hard settings respectively use differential-expression probabilities 0.20, 0.10 and 0.05; factor-location parameters 0.20, 0.10 and 0.05; factor-scale parameters 0.30, 0.40 and 0.40; common biological coefficients of variation 0.05, 0.10 and 0.20; and BCV degrees of freedom 100, 60 and 30. Down-regulation probability is 0.5. Additional dropout is absent in the easy and standard settings; the hard setting uses experiment-level dropout with midpoint 0 and shape −1. The hard double-bifurcation design also changes state abundance. Difficulty therefore jointly perturbs differential expression, biological variability and dropout. State-abundance vectors and directed truth graphs are supplied in the accompanying compact parameter record.

A separate cell-number analysis used 200, 400, 800, 1,600, 3,200, 6,400 and 12,800 cells from a common double-bifurcation source. Five paired replicates used fixed selection seeds 115900, 116000, 116100, 116200 and 116300 and nested subsets across cell counts. Each method received the identical subset in a replicate/size unit; SPARC graph-resampling seed was 159. These paired replicates measured sensitivity to sampled cell composition at each cell count.

#### 11.2 Directed topology metrics

SPARC, CStreet, CellRank 2 and moscot were compared on identical cells and exact temporal-state labels. The simulations evaluate connection recovery conditional on supplied state definitions. Native method edge selections were used without matching their counts to SPARC. For simulated CellRank 2 outputs, outgoing adjacent-time transition probabilities were normalized over the target time and retained above the uniform target-cluster probability. This is the same $p_{st} > 1/K_b$ definition used for the LARRY moscot and CellRank 2 comparisons. Predictions were scored as directed pairs, with self-edges removed and duplicate pairs collapsed. Edge precision, recall and F1 use the specified directed truth edges; a reversed edge counts as a mismatch.

HIM was used for trajectory-topology comparison following its application in trajectory-inference benchmarking[9], with the directed-graph formulation of Jurman et al.[39]. For binary source-by-target adjacency matrices $A$ and $A^*$ on $n$ labeled nodes,

$$H = \frac{\sum_{i \neq j} |A_{ij} - A^*_{ij}|}{n(n-1)}.$$

Direction is represented for the spectral component using the undirected bipartite lift

$$B(A) = \begin{pmatrix} 0 & A^\top \\ A & 0 \end{pmatrix}.$$

Laplacian frequencies are the square roots of the nonnegative eigenvalues of this lifted graph. One zero frequency is omitted; additional zeros from disconnected components are retained. The normalized spectral density is a sum of Lorentzian kernels,

$$\rho_A(\omega) = K_A \sum_j \frac{\gamma}{(\omega - \omega_j)^2 + \gamma^2}, \qquad \int_0^\infty \rho_A(\omega)\, d\omega = 1.$$

The bandwidth $\gamma$ is calibrated separately for each node count so that the empty and complete directed graphs have spectral distance one. With $I = \left[\int_0^\infty (\rho_A - \rho_{A^*})^2 d\omega\right]^{1/2}$, reported similarity is $1 - \sqrt{(H^2 + I^2)/2}$. The reported metric uses this directed Hamming normalization and bipartite spectral lift throughout.

### 12 Reproducibility and interpretation

The reproducibility materials include simulation configurations for all 256 mixed-benchmark datasets, parameter and random-seed records, source policies, selected-route definitions, gene-dynamics and pathway-classification tables, numerical figure source data and analysis code. Access to expression matrices and patient data is described in the Data sources and access section.

Experiment-specific scripts supplied inputs and parameters to the same SPARC shortest-path implementation. Inference provenance records the cell and cluster order, input paths and checksums, source policy, candidate roots, destination list, sampling fractions, seeds and complete edge-weight columns. Package and environment records are retained per run, with CPU and GPU environments recorded separately. GPU use is reported at the operation level, including exact neighbor-distance calculation and Leiden execution where applicable.

Uncertainty units followed the study design: cell resampling measured within-experiment sensitivity, clone folds measured evaluation variation and independent experiments defined biological reproducibility. Figures retain these units explicitly. Source citations describe the underlying algorithms and datasets, while the combinations and evaluation definitions above specify the SPARC implementation.

### 13 Murine Orthotopic Osteosarcoma Model and scRNA-seq Sampling

#### 13.1 Orthotopic tumor generation

Three independently established murine osteosarcoma cell lines, M1332, M408, and M1199, were used to generate syngeneic orthotopic tumors in C57BL/6 mice. The model system and establishment of tumor-derived cell lines were described previously by Nirala et al.[40]. Each mouse received an intratibial injection of $1 \times 10^6$ cells. At the study-defined euthanasia endpoint, the primary tibial tumor and visually apparent pulmonary metastatic lesions were collected. Primary and lung lesions were isolated separately and processed as single-cell suspensions for scRNA-seq. The three cell lines were treated as independent biological model systems, and primary-tumor and lung-lesion profiles were retained as distinct anatomical conditions.

##### 13.2 Murine primary-to-lung SPARC configuration

Following experiment-specific scRNA-seq quality control and cell-type annotation, malignant cells from the primary and lung compartments were represented in a shared Harmony-corrected principal-component embedding. The primary and lung objects were supplied to SPARC as ordered conditions, with primary constrained to precede lung. Organism-dependent settings were configured for mouse. CytoTRACE2 was calculated from raw counts, and root nomination was restricted to retained primary-tumor graph states. The state with the highest median cell-level CytoTRACE2 score initialized the root. P3 was fixed as the single source. All condition-permitted cluster pairs were retained as candidates; positive finite weights define traversable edges in each graph. Subsequent graph construction, minimum inverse-connectivity path optimization, Mahalanobis-whitened projection, trajectory-specific GAM fitting, and direction-specific enrichment followed the algorithmic procedures defined in the preceding sections of this Supplementary Note.

The anatomical ordering is a constraint derived from the experimental design. SPARC trajectories therefore represent transcriptomically supported state progression from primary to lung under the specified anatomical ordering.

#### 14 Spatial transcriptomic applications

##### 14.1 Registered histology display and Lung zonation

Registered H&E images from the Space Ranger v4 outputs are displayed alongside the frozen P10/L13 spatial signature scores. Full-resolution pixel coordinates are converted to micrometres using the sample-specific `microns_per_pixel`; registered hires-image pixel coordinates use `tissue_hires_scalef`. Each histology/signature pair shares the same tissue field, orientation and display scale. Image crops retain the source pixel colours. Signature maps use the stored $k = 36$ spatially smoothed scores and their established per-sample first-to-ninety-ninth-percentile colour range. The Lung RNA-defined region uses the cell annotations `OS_differentiated` and `Tumor_uncertain`. Their local fraction is calculated among the 200 spatial nearest neighbours, and fractions of at least 0.5 identify the initial region. The established procedure uses a 20 $\mu$m grid, tissue-mask closing at a 40 $\mu$m radius, hole filling and 20 $\mu$m opening, followed by region-mask closing at 100 $\mu$m and a 60 $\mu$m margin. Retained connected components contain at least the larger of 500 grid bins and 5% of the largest component. The resulting Lung region defines the core, boundary and exterior assignments for 249,756 segmented objects in Fig. 5c. Both histology/signature pairs retain their original display fields, orientation, colours, scale bars and signature values.

#### 14.2 Node programs versus full-terminal spatial-module controls

Node-specific P10/L13 signatures assess expression-state recovery in human tissue. The retained recurrent up/down modules have a different source: Pearson statistics on nine complete lung-terminal routes, including post-anchor states. Genes required route-specific BH-adjusted Pearson $P < 0.01$ and concordant direction on at least five routes; the top 20 human-matched genes by mean absolute correlation formed each module. These full-topology modules provide supplementary tissue context; the four shared-trunk GAM analyses define the local-versus-metastatic comparison. The primary/metastatic comparison uses the five primary and four shared-trunk route sets defined above.

#### 14.3 Modified neighborhood-based QC for segmented Visium HD objects

The full segmented-cell maps used the primary bone tumor specimen Bone (original specimen ID JY4) and pulmonary metastasis specimen Lung (original specimen ID JY10), which came from different patients. The limited primary specimen paired with Lung had original ID JY8 and was used only for the mature-resorptive footprint comparison. Quality control for the segmented Bone and Lung objects used a neighborhood-based procedure inspired by SpotSweeper[41]. For each cell $i$, the 36 nearest spatial neighbors $\mathcal{N}_i$, excluding the focal cell, supplied the mean $\mu_i$ and sample standard deviation $s_i$ of each QC metric:

$$\mu_i = \frac{1}{36} \sum_{j \in \mathcal{N}_i} x_j, \qquad s_i^2 = \frac{1}{35} \sum_{j \in \mathcal{N}_i} (x_j - \mu_i)^2, \qquad z_i = \frac{x_i - \mu_i}{s_i}.$$

If $s_i < 10^{-10}$, the score was set to zero. Metrics were log1p total counts, log1p detected-gene counts and mitochondrial-count percentage. Cells were flagged for $|z_i| > 3$ on counts or genes, or $z_i > 3$ on the mitochondrial metric. A hard minimum of five detected genes was also applied. Cells failing either the local-outlier or hard-minimum criteria were removed, followed by genes detected in fewer than ten cells. This implementation uses mean/standard-deviation scaling and two-sided count/gene filtering. The retained annotations contained 255,112 segmented objects in Bone and 249,756 in Lung.

#### 14.4 Osteoclast-rich spatial annotation and counting units

The Lung analysis retained 249,756 nucleus-guided segmented objects. BANKSY clustering assigned 15,537 objects to an osteoclast-associated cluster, designated the osteoclast-rich region. Individual-object marker scoring and grouping of neighboring nuclear segments yielded an estimated 1,300–1,800 osteoclasts. For section-level normalization, this estimate corresponded to approximately 0.5–0.75% of the total segmented-object count. Osteoclast-associated expression was concentrated within tumor nodules. Spatial neighborhood and ligand–receptor analyses used the cluster-level annotations.

#### 14.5 Lung zonation and boundary-associated myeloid analyses

The Lung section was divided into core, boundary and exterior zones from the local tumor-cell density and the signed distance to the tumor boundary, as described in the manuscript Methods.

Core cells lay more than 100 $\mu$m inside the boundary, boundary cells within $\pm$100 $\mu$m, and exterior cells more than 100 $\mu$m outside it. The zone inputs contained 129,893, 41,873 and 77,990 cells, respectively.

Within each zone, cell-type neighborhoods were summarized by the mean fraction of each neighbor type among 36 same-zone nearest neighbors, excluding the focal cell, divided by that type's zone-wide abundance. Ligand–receptor proximity was recomputed within each zone at 50 $\mu$m using the expression-positive cell definition, expected-pair calculation and Fisher-test procedure described in the manuscript Methods. Boundary-versus-exterior macrophage expression was compared by per-cell Wilcoxon rank-sum tests on log1p-normalized counts, requiring at least 50 cells per group. The evaluated macrophage groups contained 1,806 boundary cells and 9,536 exterior cells. Reported genes met adjusted $P < 0.05$ and $|\log_2 \mathrm{FC}| > 0.5$. These comparisons describe variation within the single Lung specimen.

#### 14.6 Paired conventional Visium data

The paired spatial analysis used conventional Visium data from GSE293065[42]. Raw 10x filtered matrices, Space Ranger tissue-position tables, image scale factors, and H&E images were used. Five paired comparisons were analyzed independently: four primary bone tumor to lung metastasis comparisons and one primary tumor to soft-tissue local relapse comparison. The abbreviation Loc denotes local relapse.

The paired conventional Visium inputs after spatial-SPARC quality control were:

- Pair 1, Pri1 → LuM1: primary accession GSM8875097, progression accession GSM8875083; retained primary/progression spots 778 / 3,799.
- Pair 2, Pri2 → LuM2: primary accession GSM8875098, progression accession GSM8875094; retained primary/progression spots 712 / 3,297.
- Pair 3, Pri3 → LuM3: primary accession GSM8875099, progression accession GSM8875095; retained primary/progression spots 1,061 / 3,122.
- Pair 4, Pri4 → LuM4: primary accession GSM8875100, progression accession GSM8875096; retained primary/progression spots 823 / 3,395.
- Pair 5, Pri5 → Loc5: primary accession GSM8875101, progression accession GSM8875082; retained primary/progression spots 1,805 / 615.

#### 14.7 Quality control and BANKSY representation

Each patient pair was processed independently. Only in-tissue spots were retained, and spots with fewer than 200 detected genes were removed. Raw counts were preserved in a separate layer. Counts were normalized to a total of 10,000 per spot and log-transformed. Up to 3,000 highly variable genes were selected within each patient pair, using the two sections as the batch variable.

BANKSY[2] was used to construct a spatially informed molecular representation. Spatial neighborhoods were restricted to individual tissue sections. For every spot, zeroth-order neighborhood expression was calculated from the six nearest within-section spatial neighbors using uniform weights. The neighborhood contribution was $\lambda = 0.2$. The resulting expression-plus-neighborhood matrix was jointly reduced to 50 principal components within each pair. Tissue-section identity was fully confounded with progression status, so Harmony was omitted. An expression-only $\lambda = 0$ representation served as a control; the spatially augmented representation defined the main analysis.

##### 14.8 Pair-specific SPARC inference and spatial projection

SPARC used the first 30 components of the BANKSY-derived representation, $K = 29$ cluster-excluded outside-cluster neighbors, Leiden resolution 0.8, the complete condition-permitted candidate universe, and random seed 159. Clustering was performed separately within the primary and progression sections. CytoTRACE2 was applied to raw counts from the primary section only, and the retained primary graph state with the highest median spot-level score initialized the root. The fixed roots for Pairs 1–5 were Pri3, Pri2, Pri2, Pri2 and Pri6, respectively. Minimum-cost paths, cell-level projection, normalized pseudotime, and GAM-based gene dynamics were computed as described above. Dynamic genes required Benjamini–Hochberg-adjusted $P < 0.05$ and explained deviance $> 0.10$.

For tissue mapping, all in-tissue spots were displayed as a low-opacity background, and spots assigned to a selected SPARC trajectory were colored by normalized pseudotime. Local pseudotime arrows were calculated separately within each section. For a candidate trajectory spot, up to 30 local same-trajectory neighbors were used in a distance-weighted linear regression of pseudotime on H&E coordinates after normalizing distance by the median Visium spot spacing. Arrow orientation follows the positive fitted pseudotime gradient, opacity represents gradient magnitude, and length is fixed for display. These arrows summarize the fitted local spatial gradient of pseudotime.

##### 14.9 Tissue-compartment and neighborhood analysis

Marker-supported dominant molecular compartments were assigned to Visium spots in GSE293065 for visualization. Within each section, every marker gene was standardized across spots after normalization and log transformation, and Z-scores were clipped to $[-4, 4]$. A program score was the mean of its marker Z-scores. A dominant label required a leading score of at least 0.10 and a margin of at least 0.12 over the next program; all other spots were labeled Mixed/low signal. The programs represented osteosarcoma, myeloid, mature osteoclast, lymphoid/plasma, vascular/stromal, and lung-parenchymal compartments. These labels summarize the dominant molecular program in each mixed Visium spot.

For every Pair 4 lung-metastasis trajectory spot, the focal spot was excluded and its six nearest spatial spots were identified. Neighbor-label fractions were summarized across four rank-based pseudotime quartiles. Continuous trends were tested by Spearman correlation, with Benjamini–Hochberg correction across compartments separately within each trajectory.

##### 14.10 Branch programs and recurrence across patient pairs

Trajectory enrichment used each trajectory's complete set of significant dynamic genes. Increasing and decreasing genes were tested separately with a one-sided hypergeometric test against MSigDB 2026.1 Hallmark, Gene Ontology, and canonical-pathway collections. The background was the full set of genes tested by the trajectory-specific GAM. Gene sets were required to contain 10–1,000 background genes and at least three dynamic-gene overlaps. Benjamini–Hochberg FDR $< 0.05$ defined significance. Enrichment quantified overlap between trajectory-associated dynamic genes and curated gene sets.

For Extended Data Fig. 10d, the KEGG antigen-processing-and-presentation components were labeled HLA-II presentation for the tumor-progression trajectory and lysosomal processing for lung-local osteoclast progression. Functional labels reflect the contributing genes, with three high-deviance representatives displayed for each selected pathway.

Patient-level recurrence was summarized for predefined osteogenic-matrix, stress-adaptation, osteoclast-resorption, antigen-presentation, alveolar-epithelial, and macrophage-associated modules. Within a terminal trajectory, a module had qualifying signal when at least one module gene had adjusted GAM $P < 0.05$ and explained deviance $\geq 0.10$. The trajectory-level effect was the median late-minus-early change among qualifying genes. Each patient pair contributed one independently reconstructed graph and defined the patient-level replication unit.

##### 14.11 External conventional Visium route–niche validation

The external cohort contained 26 conventional Visium sections from 19 patients in GSE293065: nine primary bone tumors, fourteen distant metastases and three local relapses. Counts were normalized to 10,000 per spot and log-transformed. The four mouse metastatic-route signatures were mapped to human genes and scored with `scanpy.tl.score_genes`, using `use_raw=FALSE`, control-set size $\min(50, n_{\text{genes}})$ and random seed 0. The osteoclast markers specified by the frozen source configuration were removed from route signatures; genes in route signatures were excluded from the macrophage panel.

The canonical osteoclast panel comprised CTSK, ACP5, MMP9, ATP6V0D2, NFATC1, DCSTAMP, OCSTAMP, CALCR, SIGLEC15, OSCAR, ATP6V0A3 and TCIRG1. The terminal panel comprised CTSK, ACP5, MMP9 and DCSTAMP. Within each section, the upper score quartile defined niche-high spots. Enrichment was the mean route score in niche-high spots minus its mean in the remaining spots. Macrophage residualization fitted each route score to an intercept and macrophage score by ordinary least squares within a section; the residuals entered the same canonical osteoclast enrichment calculation.

MR1-minus-MR3 differences across sections were evaluated with two-sided paired Wilcoxon signed-rank tests. The canonical, terminal and macrophage-residualized contrasts yielded $P = 0.0014439225$, 0.3940727115 and 0.0357243717, with positive differences in 20, 12 and 17 of 26 sections, respectively. These displayed $P$ values are unadjusted. Four-route comparisons with macrophage/monocyte, endothelial and T-cell niches used the same upper-quartile enrichment definition, with mean and SEM across sections. Enrichment measured spot-level expression associations in the external conventional Visium cohort.

### 15 Mouse Lung Myeloid-to-OLC Analysis

#### 15.1 SPARC-SS trajectory inference

The analysis used 7,681 lung myeloid cells assigned to 12 fixed region-cluster states. The existing 30-dimensional within-lung Harmony representation entered SPARC without re-estimating the clusters. Mono1 had the highest median CytoTRACE2 potency score among the 12 states (0.285) and was fixed as the single source. The cluster-excluded graph used the 49 nearest outside-cluster cells for each cell. All condition-permitted pairs were candidates, with same-condition connections reciprocal; only positive finite weights were traversable in a round. Connection strength $w_{ij}$ was converted to cost $C_{ij} = 1/w_{ij}$ before exact Dijkstra inference on the complete permitted graph.

The four requested endpoints were Mono7, Mono6, M$\varphi$4 and OLC. The modal paths were Mono1→Mono2→Mono3→Mono5→Mono7, Mono1→Mono2→Mono4→Mono6, Mono1→M$\varphi$3→M$\varphi$4 and Mono1→M$\varphi$3→OLC. Cell-sampling uncertainty was evaluated independently at 70%, 80% and 90%, with 1,000 rounds per fraction. Every round sampled the specified fraction without replacement within each cluster, rebuilt the outside-cluster neighbor graph, recalculated the complete connection-strength vector and ran exact Dijkstra with the fixed source, endpoints and permitted-edge set. The OLC route Mono1→M$\varphi$3→OLC was reachable and modal in all 3,000 rounds.

#### 15.2 Pseudotime, gene programs and human spatial scoring

For Mono1→M$\varphi$3→OLC, full-data covariance whitening was applied to the 30 Harmony dimensions. The fixed full-data projection reference was used for the 1,975 retained route cells. A discordant cell with an original Osteoclast annotation and a Mono1 reclustering assignment was excluded from downstream program analysis. Each retained cell was projected to its nearest adjacent cluster-center segment, and cumulative segment length defined pseudotime. Scanpy[5;43] gene scores were calculated for predefined osteoclast-resorption, fusion-commitment and AP-1 immediate-early programs; running means used a centered window of 98 cells. Nfatc1 log-normalized expression was summarized with the same window.

The spatial four-program reference used 5,042 mouse myeloid cells, grouped into early monocyte/inflammatory (3,485), macrophage-intermediate (635), osteoclastogenic pre-OLC (801) and mature-resorptive (121) categories. One-versus-rest Wilcoxon tests selected 30 positive markers per category with detection in more than 10% of group cells. The source grouping was My8/My5/My3, My0/My2, My7 and My11, respectively. Human ortholog matching recovered 13, 22, 19 and 23 genes in these categories. Bone and Lung counts were normalized to 10,000 and log-transformed before `scanpy.tl.score_genes` with `use_raw=FALSE`, control-set size $\min(50, n_{\text{genes}})$ and seed 0. The Lung four-program map used the argmax after averaging each score over 36 spatial neighbors. This broader expression reference and the 1,975-cell selected mouse route define their respective analysis inputs.

The neighborhood analysis used Lung tumor cells assigned to metastatic routes MR1–MR4. For each sender, neighbors within 25 or 50 $\mu$m were found from the tissue coordinates and the sender itself was excluded. Mature-resorptive OLC-high objects were defined by the upper decile of the four-program reference score across Lung. The plotted 25-micron comparison uses 127,778 malignant-lineage senders and excludes objects assigned to the osteoclast-rich region from this sender population. For each route and radius, enrichment compared the fraction of route-assigned senders with at least one neighboring object assigned to the osteoclast-rich region or at least one OLC-high neighbor against senders assigned to the other three routes. Observed-to-expected ratios, one-sided Fisher exact tests and Benjamini–Hochberg adjusted P values were reported.

### 16 TF regulon activity and cross-species concordance

#### 16.1 Expression inputs and curated regulatory network

The activity comparison used the full lung myeloid analysis pools. The mouse input contained 7,681 cells, labeled as 6,010 monocytes, 1,549 macrophages and 122 osteoclasts, with 22,489 genes. The human lung-metastasis input from GSE152048 contained 1,763 cells, labeled as 272 monocytes, 1,052 macrophages and 439 osteoclasts, with 13,764 genes. Mouse labels came from the annotated lung myeloid object; human labels came from its canonical-marker-based lung myeloid annotation. These are the activity-scoring pools, while the selected mouse trajectory used the separate 1,975-cell input described above.

For the human activity input, lung myeloid cells were clustered using 30 Harmony coordinates, 50 neighbors and Leiden resolution 1.0. Canonical signatures comprised FCN1, VCAN, S100A8, S100A9, CD14 and LYZ for monocytes; C1QA, C1QB, C1QC, APOE, MRC1, CD68, TREM2, MARCO and FABP4 for macrophages; and CTSK, ACP5, MMP9, OSCAR, CALCR, CA2 and ITGB3 for osteoclasts. Cell-level signature scores were averaged by cluster, standardized across clusters within each signature and assigned by the leading score. An osteoclast-labeled cluster with mean log-normalized resorption-panel expression below 1.0 was grouped with macrophages for this comparison.

The gene-by-cell matrices contained the stored log-normalized RNA expression. Raw counts had been normalized to 10,000 per cell and log1p-transformed; the exported matrices retained the genes available in each analysis object. Signed CollecTRI TF–target interactions[44] were retrieved from the OmniPath resource[45]. Empty gene symbols were excluded, and duplicate source–target symbol pairs retained their first record. An interaction annotated as inhibitory and not stimulatory received mode-of-regulation weight $-1$; all other retained records received $+1$. The deduplicated network contained 62,411 TF–target symbol pairs for 1,201 TF symbols.

For mouse scoring, each human TF and target symbol was converted to mouse-style capitalization by uppercasing the first character and lowercasing the remaining characters. The resulting symbols were intersected with the measured mouse genes. This implementation uses symbol correspondence for network transfer. Network and input provenance, together with the activity source tables, specify the analysis inputs.

##### 16.2 Univariate linear-model activity scores

decoupleR[46] estimated a per-cell TF activity score with `run_ulm`, using `source`, `target` and signed `mor` columns and `minsize=5`. A TF required at least five target genes represented in the expression matrix. For regulator $r$ and cell $c$, the ULM relates gene expression to its regulatory-weight vector:

$$x_{g,c} = \beta_{0,r,c} + \beta_{1,r,c} m_{r,g} + \epsilon_{g,c}, \qquad a_{r,c} = \frac{\widehat{\beta}_{1,r,c}}{\mathrm{SE}(\widehat{\beta}_{1,r,c})}.$$

The weight $m_{r,g}$ is the signed TF–target annotation and is zero for measured genes outside that regulon. The reported activity is the fitted slope t-statistic. No additional cross-cell z-standardization was applied to these scores. There were 749 scored mouse TFs and 668 scored human TFs. Scores were averaged within each species and each of its monocyte, macrophage and osteoclast annotation groups.

##### 16.3 Matching TF activity contrasts between species

For each scored TF, the contrast was the mean osteoclast-labeled activity minus the mean monocyte-labeled activity. TF names were converted to uppercase and matched by symbol across species. All matched contrasts finite in both species entered the direction comparison:

$$\Delta_r^{(s)} = \overline{a}_{r,\mathrm{OC}}^{(s)} - \overline{a}_{r,\mathrm{Mono}}^{(s)}, \qquad f_{\mathrm{same}} = \frac{1}{|\mathcal{R}_\cap|} \sum_{r \in \mathcal{R}_\cap} \mathbf{1}\left[\mathrm{sign}(\Delta_r^{(m)}) = \mathrm{sign}(\Delta_r^{(h)})\right].$$

The comparison contained 662 finite matched TF contrasts, with 417 sharing direction (63.0%) and 245 having opposite directions. All matched contrasts had nonzero differences. One plotted point represents one matched TF activity contrast. This readout describes directional correspondence in activity estimated from the transferred prior network.

#### 17 Cross-Resolution Lung Osteoclast Analyses

##### 17.1 Conventional Visium analysis

The conventional Visium analysis used only the Pair 4 lung-metastasis section and its $\lambda = 0.2$ BANKSY representation. Whole-section BANKSY clustering at resolution 2.0 identified complete myeloid-lineage clusters. The retained lineage was re-clustered at resolution 1.5 after evaluating resolutions 0.5–2.0 across random seeds 159, 271, and 907. The final input contained 30 macrophage-like spots, 99 osteoclast-precursor spots, and 145 mature- osteoclast spots. The macrophage-like cluster was the earliest reproducibly resolved state and supplied the starting population for the ordered analysis.

SPARC-SS used 30 components, $K = 19$ cluster-excluded outside-cluster neighbors, Leiden resolution 0.8, the complete condition-permitted candidate universe, and random seed 159. The highest median macrophage-minus-mature-osteoclast program contrast selected the macrophage root for this within-section, marker-anchored ordering. The resulting paths describe differentiation-state progression within one lung section.

##### 17.2 Segmented Visium HD analysis

The Lung Visium HD analysis used 6,483 Space Ranger segmented objects and 18,085 genes. The BANKSY representation was supplied to SPARC. BANKSY-Leiden clustering used 30 neighbors, resolution 0.55, and random seed 159. A monocyte-like early program selected LuM-C3 as the root. SPARC identified natural graph leaves before marker annotation; subsequent annotation identified LuM-C1 as both a natural leaf and the strongest osteoclast-program state. The osteoclast trajectory was LuM-C3 → LuM-C0 → LuM-C2 → LuM-C1 and contained 5,452 projected objects.

Terminal-state stability was assessed across four Leiden resolutions and three random seeds. The osteoclast-rich state remained a natural leaf in all 12 settings. Cross-resolution agreement between conventional Visium and Visium HD was evaluated using the direction and late-minus-early effect of shared canonical genes. H&E mapping used the original Lung whole-slide image and the Space Ranger coordinate transformation. Local arrows represent spatial pseudotime gradients.

#### 18 Data sources and access

##### 18.1 Osteosarcoma and additional applications

The paired mouse osteosarcoma scRNA-seq and patient-derived Visium HD data generated in this study have been deposited under GEO accessions GSE343530 and GSE343536, respectively. Published human osteosarcoma scRNA-seq data are available under GSE152048, and the conventional Visium cohort under GSE293065. Longitudinal HGSOC data are available under GSE165897.

##### 18.2 LARRY expression and clone assignments

The LK1, LK2 and LSK analyses used the `stateFate_inVitro` release in GSE140802, sample GSM4185642[24]. Processed normalized expression is supplied in `normed_counts.mtx.gz`, with `gene_names.txt.gz`. Cell-to-clone membership is supplied in `clone_matrix.mtx.gz`. The cell-barcode, library-name and metadata files specify the common row order, library identity, sampling time and experimental population. All filenames have the prefix `GSM4185642_stateFate_inVitro_`; they can be downloaded individually from the GEO sample page or from `GSE140802_RAW.tar`. Expression and lineage information were joined using these shared cell identifiers and row metadata.

##### 18.3 Tumor Cas9 expression and lineage releases

KP-Tracer inputs came from the authors' version-0.1 `KPTracer-Data.tar.gz` release at https://doi.org/10.5281/zenodo.5847462. The raw-count layer of `expression/adata_processed.combined.h5ad` supplied expression. `KPTracer_meta.csv` records whether a cell has RNA and TargetSite measurements and identifies its metastatic family. The `trees/` directory contains released character matrices and reconstructed Newick trees.

macsGESTALT expression matrices, feature names and cell barcodes came from `GSE173958_RAW.tar`. The M1 and M2 archives in Mendeley Data version 1, https://doi.org/10.17632/t98pjcd7t6.1, supplied the expression-matching metadata, static-clone assignments, editing records, full edge lists and reconstructed clone phylogenies. The two mice were matched and evaluated separately.

Quinn A549 xenograft inputs came from GSE161363. RNA samples GSM4905335, GSM4905337, GSM4905339 and GSM4905341 correspond to M5k, M30k, M10k and M100k. Their lineage samples are GSM4905334, GSM4905336, GSM4905338 and GSM4905340, respectively. The RNA releases contain expression matrices, gene lists, barcodes and cell metadata. The paired lineage releases contain allele tables, character matrices and tree archives.

##### 18.4 Simulation and study-generated data

The four-generator mixed within/cross-condition design includes the complete complex reference topologies, transition modes, descendant allocations, expression regimes, generator parameters and seed-specific unit manifests. The design contains 256 datasets, each with 8,000 retained cells and 2,000 simulated genes. Seeds 159, 161 and 162 define 96 calibration datasets, and 164–168 define 160 test datasets. The conventional Splatter topology, difficulty and cell-number experiments retain their own separate configuration and unit manifests. Generator scripts and the common observation-layer code document how the count matrices and reference annotations were produced.

Synthetic count matrices can be regenerated from the supplied generator scripts, reference topologies, parameter files and seed manifests. These materials, empirical path-frequency estimates, benchmark evaluation results and numerical figure source data are available from the authors during peer review and will be released on GitHub upon acceptance of the manuscript.

Extended gene-program comparisons for MR1–MR4 and routes to L6 and L7, including all-gene dynamics and pathway-scoring tables, are available from the authors during peer review and will be released on GitHub upon acceptance of the manuscript.

Gene-level trajectory results and the numerical outputs needed to interpret and reproduce the reported comparisons will accompany the versioned study release. These supporting data will be available from the authors during peer review and released through the GitHub repository specified above upon acceptance.
